# Multimodal Molecular Mapping of the Vasculature in Human Cortex Reveals Lipid Markers of Cerebral Amyloid Angiopathy

**DOI:** 10.64898/2026.03.13.711741

**Authors:** Cody R. Marshall, Felipe A. Moser, Claire F. Scott, Lissa Ventura-Antunes, Wilber Romero-Fernandez, Łukasz G. Migas, Léonore E. M. Tideman, Madeline E. Colley, Martin Dufresne, Matthew S. Schrag, Raf Van de Plas, Jeffrey M. Spraggins

**Author notes:** These authors contributed equally to this work (Cody R. Marshall, Felipe A. Moser).

## Abstract

Cerebral amyloid angiopathy (CAA) commonly co-occurs with Alzheimer’s disease (AD), yet the molecular changes that accompany vascular β-amyloid deposition in human tissue remain incompletely defined. Herein, we use a novel imaging approach that combines matrix-assisted laser desorption/ionization imaging mass spectrometry (IMS) with immunofluorescence microscopy on the same sections of postmortem human frontal cortex to map the lipid microenvironment of leptomeningeal vasculature in cases with and without CAA. Autofluorescence-guided regions-of-interest were imaged by IMS in both negative and positive ion modes and registered to post-IMS-acquired microscopy images. Immunofluorescence microscopy using collagen IV, α-smooth muscle actin (αSMA), and thiazine red enabled automated segmentation of total, amyloid-positive, and amyloid-negative vasculature regions. A CAA index, the ratio of amyloid-positive area to total vasculature area in a region imaged by IMS, was used to define vasculature and classify each case into having CAA, or CAA-present and not having CAA, or CAA-absent. An interpretable machine learning approach (XGBoost models with Shapley additive explanations for interpretation) was trained on pixel-level spectra and identified lipid signatures of vascular identity shared across groups as well as class-specific marker candidates that distinguished CAA-present from CAA-absent vasculature. CAA-present vessels were enriched for gangliosides (*e.g.*, GM1), whereas CAA-absent vessels were characterized by higher contributions from phosphatidylserines (*e.g.*, long-chain polyunsaturated PS species). Univariate differences were inconsistent between the two groups, but multivariate models in negative mode yielded stable discriminatory features. These results define spatial lipid correlates of vascular amyloid pathology in the human brain and establish a multimodal framework for mechanistically linking lipid metabolism, vascular integrity, and CAA in AD.

## 1 Introduction

Alzheimer’s disease (AD) is a complex neurological disorder and while its most famous neuropathological features are β-amyloid plaques and neurofibrillary tangles, most autopsy studies suggest that a wide range of pathologies contribute to disease progression.[1] Emerging drug treatment strategies for AD, focusing on clearing β-amyloid from the brain using monoclonal antibodies, have had some success in slowing AD symptoms, but they have not yet been successful at preventing disease onset or progression.[2] Furthermore, patients with concurrent vascular deposition of β-amyloid, a condition called cerebral amyloid angiopathy (CAA), treated with anti-β-amyloid immunotherapies have tended to have a high rate of side effects from these new treatments.[3][4][5] These complexities suggest a more elaborate molecular picture of AD than previously supposed and have increased the priority of understanding the molecular underpinnings of CAA and other common AD co-pathologies. A systems biology approach will enhance our understanding of associated pathologies, such as CAA, and their interconnectedness with AD.

CAA frequently co-occurs with AD, with 80% of patients with AD having some level of CAA pathology.[6] CAA is independently associated with cognitive impairment and intracerebral hemorrhage,[7][8] and has so far been molecularly defined by β-amyloid deposition within the walls of the cerebrovasculature.[9] However, several researchers have reported β-amyloid deposition occurring at spatially distinct regions away from vessel wall breakdown, suggesting the relationship between β-amyloid and vascular degeneration may be indirect.[8][10] The molecular mechanisms involved in the transition from β-amyloid deposition to cognitive decline and hemorrhage remain largely unknown. Genomic studies show that CAA-risk is increased in patients carrying one or more ε4 alleles of apolipoprotein E (ApoE4), a gene likely involved in lipid metabolism.[11] In addition, large-scale lipidomic studies suggest that aberrant lipid metabolism plays a critical role in AD and CAA progression.[12][13][14][15][16][17] Furthermore, prior reports have also noted that various matrix metalloproteinases play a role in vascular degeneration within the context of CAA.[18][19] Vessels containing CAA degenerate over time, as observed by a loss of vascular smooth muscle (VSM).[20] This process hinders the vessel’s ability to autoregulate blood flow, exacerbating the brain’s ability to mitigate disease.[21] Collectively, these observations suggest a more complicated molecular picture of how β-amyloid deposition leads to CAA’s effects and, ultimately, vessel rupture. Deeper characterization of the vascular microenvironment is needed to provide further molecular insight into AD and CAA pathogenesis.

Spatial multi-omics and molecular imaging are rapidly growing strategies to expand beyond traditional bulk omics by adding spatial context to molecular information and thereby enhancing our understanding of the tissue microenvironment.[22] By linking genomic clues to lipid metabolism, differential lipidomic expression in bulk mass spectrometry analysis, and the role of vascular smooth muscle cells, spatial lipidomic analysis combined with imaging of cell types in the vasculature could reveal important information regarding the microenvironment of AD and AD-associated pathologies like CAA.

While the importance of lipids in AD and CAA is clear, imaging a wide variety of lipid species is often not feasible through traditional imaging approaches, such as immunofluorescence microscopy, due to lack of specificity and the need for a prohibitively large number of highly specific markers for a molecular profile that is unknown a priori. In contrast, matrix-assisted laser desorption/ionization (MALDI) imaging mass spectrometry (IMS) is a molecular imaging technology uniquely suited to imaging lipids as the use of probes is not necessary.[23][24][25][26][27][28][29] MALDI IMS has been demonstrated in the brain as a powerful tool for imaging a wide range of molecules,[30][31][32][33][34][35] with some researchers successfully using it to characterize amyloid isoform composition within neuritic plaques.[35][36][37] MALDI IMS has even been used to image the spatial lipidomic environment surrounding amyloid plaques.[38][39][40][41] To leverage the lipid imaging capabilities of MALDI IMS in the study of AD-associated pathologies like CAA, multimodal imaging approaches are needed to correlate IMS data with vasculature features observed in immunofluorescence microscopy. For example, newly developed integration strategies combining MALDI IMS and immunofluorescence microscopy have linked lipid information to specific cell types within tissues, such as the human kidney.[42]

Herein, we utilize an integrated multimodal molecular imaging integration approach to distinguish lipid profiles between healthy and CAA-diseased tissue features and cell types. We focus specifically on the leptomeninges, vascular tissue most vulnerable to CAA,[43] to link spatial lipid markers with histopathology. Using autofluorescence microscopy-guided MALDI IMS, we imaged leptomeningeal vessels at 10 µm pixel size. This was then registered to post-IMS-acquired immunofluorescence images of the same sections, based on which vasculature and CAA features were segmented automatically. We propose a quantitative CAA index (β-amyloid/vasculature area) to stratify tissues and cases into CAA-present and CAA-absent vasculature subgroups, enabling pixel-specific comparisons within vasculature rather than using bulk tissue. We then employ an interpretable machine learning approach to identify, among hundreds of measured lipid species, those that exhibit marker potential for vasculature in general as well as for CAA presence or absence.[44][45] This approach uses eXtreme Gradient Boosting (XGBoost) models to mine these large, high-dimensional datasets for such relationships, and Shapley Additive exPlanations (SHAP) as a model interpretability method to determine which molecular species inputs hold marker potential.[46][47][48] The resulting spatial lipid signatures suggest the loss or redistribution of structural phospholipids in CAA and a potential mechanistic link between lipid metabolism and β-amyloid deposition. Together, this multimodal framework establishes a basis for identifying potential biomarkers and therapies at the neurovascular interface in AD.

## 2 Materials and Methods

### 2.1 Materials

Human brain samples were obtained from an ongoing tissue resource at Vanderbilt University Medical Center, with supervision from the institutional review board, IRB #180287. Tissue cases and/or their surrogates provided written informed consent to permit tissue use in research applications. All specimens were thoroughly de-identified to limit any risk to patient privacy. Ammonium formate, tris-buffered saline (TBS), fish gelatin, 100 mM glycine, 0.1% Triton X-100, bovine serum albumin, and 0.05% Tween-20 were purchased from Sigma-Aldrich (St. Louis, MO). The matrix aminocinnamic acid (ACA) and 4% paraformaldehyde (PFA) were bought from Thermo Scientific (Waltham, MA). High-performance liquid chromatography (HPLC)-grade acetone was purchased from Fisher Scientific (Pittsburg, PA). Alpha smooth muscle actin (αSMA) (AB124964) and normal donkey serum (AB7475) were purchased from Abcam (Cambridge, UK). Collagen IV (M3F7) was purchased from NeoBiotechnologies (Union City, CA). Thiazine red was purchased from Chemsavers (Bluefield, VA). DAPI fluoromount-G was purchased from Southern Biotech (Homewood, Alabama)

### 2.2 Sample preparation

The frontal cortex of postmortem human brains was frozen in liquid nitrogen, embedded in 15% fish gelatin, and stored at −80 *^◦^*C. A total of 13 human cases with varying levels of AD and CAA, as determined by the Vonsattel scale,[49] were selected for the imaging study. A total of 6 serial tissue sections were collected from each of 13 human cases. Four sections were omitted due to technical difficulties with sample handling, resulting in 74 vasculature sections being imaged (Figures S1-S3). Three out of the six serial sections per case were used as technical replicates for negative ion mode IMS. The other three were used as technical replicates for positive ion mode IMS. The tissue was cryosectioned at 10 µm thickness using a CM3050 S cryostat (Leica Biosystems, Wetzlar, Germany). The sections were thaw-mounted on a warm plate at 30 *^◦^*C onto indium tin oxide (ITO) coated glass slides (Delta Technologies, Loveland, CO). Autofluorescence microscopy images of the samples were collected immediately before subsequent vacuum sealing and preservation at −80 *^◦^*C, and before any subsequent imaging workflows.

The autofluorescence microscopy employed standard DAPI, eGFP, and DSRed fluorescent filters using a Zeiss AxioScan.Z1 slide scanner (Carl Zeiss Microscopy GmbH, Oberkochen, Germany), equipped with a Colibri7 LED light source. Tissue sections were then washed three times with chilled 150 mM ammonium formate at 4 *^◦^*C for 45 seconds each to remove endogenous salts and then dried with nitrogen gas to remove excess solution. An in-house developed sublimation device was used to sublimate 10 mg of aminocinnamic acid (5 mg/ml in acetone) onto three sections per sublimation run.[50] The matrix sublimation temperature reached up to 200 *^◦^*C for 15 minutes, while the samples were simultaneously cooled to −78 *^◦^*C using a dry ice and acetone slurry.[51] After sublimation, the slides were placed on a 100 *^◦^*C hot plate for 15 seconds to anneal the matrix, facilitating further recrystallization. After MALDI IMS data acquisition (see below), autofluorescence microscopy images were also acquired post-IMS using a Zeiss AxioScan.Z1 fluorescence slide scanner, using eGFP and DSRed fluorescence filters and a monochromatic brightfield image.

### 2.3 MALDI IMS Acquisition

MALDI IMS experiments were performed on a prototype MALDI timsTOF (Bruker Daltonik, Bremen, Germany) in both negative and positive ionization Q-TOF mode.[29] Autofluorescence microscopy images were imported into flexImaging and used to manually define regions of interest (ROIs) containing leptomeningeal tissue. Each established ROI comprises approximately 50,000 MALDI IMS pixels for each polarity. In both negative and positive ion modes, the IMS data were collected with the beam scan off, a 10 µm pitch, and a 10 µm spot diameter. The laser power was set to 35% with 40 shots per pixel, in both modalities. See Tab. S1 for more information on MALDI IMS instrument parameters.

### 2.4 Immunofluorescence Microscopy

After acquiring the post-IMS autofluorescence microscopy image, immunofluorescence microscopy was performed on the same tissue section. The MALDI matrix was removed from the tissue with a series of ethanol (EtOH) washes (70% to 90% EtOH). The tissue was fixed in 4% PFA solution for 15 minutes. The PFA was removed using a series of increasing sucrose solutions (10% to 30% sucrose). The tissue was placed in a sealed petri dish with 1X TBS and photobleached for 48 hours using an LED lamp (BESTVA DC Series 1200W LED Grow Light Full Spectrum) at 4 *^◦^*C. The sections were then washed with a 100mM glycine/1X TBS/0.1% Triton X-100 buffer for 30 minutes. Sections were blocked for 60 minutes at 37 *^◦^*C in a solution containing 10% normal donkey serum in 1X TBS/0.1% bovine serum albumin. Primary antibodies for collagen IV, αSMA, and a thiazine red stain were used to stain the tissue. Tissue was then washed 4 times for 10 minutes each in 1X TBS/0.05% Tween-20 and then cover slipped with a DAPI fluoromount.

### 2.5 Multimodal Image Registration and Data Analysis

To analyze the different types of images acquired from the same section in a multimodal manner, each modality was registered to a common spatial coordinate system using in-house developed registration software. The immunofluorescence microscopy images and pre-IMS autofluorescence microscopy images were co-registered with post-IMS autofluorescence microscopy images using the wsireg software, employing a combination of rigid and affine transformations.[52] The pre-IMS autofluorescence microscopy image was used as a pass-through modality for the registration of the immunofluorescence microscopy image onto the post-IMS autofluorescence microscopy image. The MALDI IMS pixels were then registered to the post-IMS autofluorescence microscopy image by manually matching a subset of these pixels to the corresponding laser ablation marks using the in-house developed tool image2image.[53][54][55] The full-resolution, co-registered images were stored in a vendor-neutral pyramidal OME-TIFF file format. After registering all modalities to their corresponding post-IMS autofluorescence microscopy image, spatial delineations (*i.e.*, segmentations) of the vasculature based on the immunofluorescence microscopy modality were used for further analysis.

### 2.6 Tissue Feature Annotations

To perform tissue segmentation within the leptomeningeal vasculature, we first defined a ROI delineating vasculature within the leptomeninge, overlapping with the area where MALDI IMS was acquired based on post-autofluorescence data. This ROI established the boundaries for all subsequent analyses, cropping out the surrounding area. Then, a leptomeninge mask was generated to enable exclusion of all non-leptomeninge tissue. This mask was obtained through an initial prediction using the Segment Anything Model (SAM),[56][57][58] and was further manually curated in the open-source digital pathology software QuPath.[59] The images and leptomeninges masks were then cropped using the vasculature ROI.

Next, vasculature segmentation was performed on the Cy5 (collagen IV) channel of each cropped image using the Moran quadrant map method proposed by Tideman et al.[60] For this, each image was first downsampled by a factor of 4 to reduce noise and minimize artifacts caused by IMS-related tissue damage (e.g., laser ablation damage). Then, the local Moran’s I was calculated using a cumulative fifth-order Queen contiguity criterion. Pixels that exhibit a positive local Moran’s I together with above average intensity were designated as vasculature with a value of 1, while the rest were designated as background with a value of 0. This vasculature segmentation mask was then upsampled back to the original pixel size using a bilinear interpolation, and binarized using a 0.5 threshold.

Finally, the thiazine red (amyloid) and Cy7 (smooth muscle actin) channels were binarized using manual thresholding and then combined with the vasculature masks to produce segmentations of the plaque and actin regions. This was then refined by filling small holes (<200 pixels, 84.5 µm^2^) and removing small objects (<200 pixels, 84.5 µm^2^), mitigating artifacts related to damage introduced during the MALDI IMS acquisition.

### 2.7 MALDI IMS Data Preprocessing

The MALDI IMS measurements were exported from the Bruker timsTOF format (.d) into a custom binary format for downstream analysis. Each pixel reported centroided mass spectral peaks spanning the full acquisition range, and these were reconstructed into pseudo-profile spectra using Bruker’s SDK (v2.21). We performed mass alignment on these profiles using the msalign library (v0.2.0),[61] using at least six reference peaks present in at least 50% of all pixels. Then, the dataset’s mass axis was calibrated to approximately *±*1 ppm accuracy using theoretical *m/z* values of at least five reference peaks. After calibration, spectra were intensity normalized using a standard total ion current (TIC) metric. Subsequently, an average mass spectrum was computed for each tissue section, aggregating all pixel spectra within that section’s MALDI IMS ROI. Peak picking of these average spectra yielded 639 and 471 distinct *m/z* features in negative and positive ionization modes, respectively. Notably, isotopic peaks were retained and not excluded from the subsequent classification workflow.

### 2.8 LC-MS/MS

Liquid chromatography with tandem mass spectrometry (LC-MS/MS) was used to identify lipid species that could not be confidently identified from MALDI IMS measurements, which includes isomeric and isobaric species. For each case, 3-4 sections of 10 µm thickness were allocated to glass vials in addition to 5 µL of 100 µg/mL internal standards Avanti Equisplash, C24:1 mono-sulfo galactosyl(ß) ceramide-d7 (d18:1/24:1) (Avanti), and 18:2 Cardiolipin-d5 (Avanti) and methanol was added until the tissue was submerged. Metal beads were added to each vial, and the tissue was homogenized with a vortex for 1 minute. Vials were placed on dry ice for 5 minutes and sonicated on ice for 30 minutes. Lipids were extracted from the solution with 1600 µL of ice-cold MTBE and 400 µL of ice-cold water, then diluted to 400 µL in ice-cold methanol. All vials were then centrifuged at 100*×*g for 10 minutes at 4°C, then held on ice for 10 minutes. The top layer, which was 600 µL, was extracted to a new vial and dried under nitrogen. The sample was resuspended in 300 µL of methanol. Reverse-phase separation was performed on a Waters Premier UHPLC with a 2.1*×*100 mm Waters Premier CSH-C18 column heated to 60 *^◦^*C. Mobile phase A consisted of 60:40 ACN:H_2_O with 10 mM ammonium formate and 0.1% formic acid. Mobile Phase B consisted of 90:10 IPA:ACN with 10 mM ammonium formate and 0.1% formic acid. The total analysis time was 25 minutes, followed by a 10-minute equilibration time. A timsTOF fleX mass spectrometer used parallel accumulation-serial fragmentation (PASEF) and MS/MS stepping to determine the collision cross section (CCS) of each species, enabling simultaneous measurement of both low-mass fragments and higher-molecular-weight lipids.

### 2.9 Univariate Molecular Comparison

The MALDI IMS ion images were normalized by TIC and subsequently inter-sample normalized using the mean-fold change method.[62] This normalization approach ensured consistent median intensities across the cohort while preserving relative biological signatures. Ion intensities were then extracted from defined vasculature masks to compare CAA-present and CAA-absent samples. For each ion, a single representative value per sample was calculated by averaging the intensities of all pixels within the respective mask. Statistical significance was determined using a univariate analysis with bootstrap resampling (10,000 iterations). This process produced robust molecular distributions from which *p*-values were derived. Multiple testing correction was performed using the Benjamini-Hochberg procedure (multipletests from statsmodels.stats.multitest Python library) to control the False Discovery Rate (FDR), yielding q-values. The relationship between effect size and statistical significance was visualized using volcano plots, with log_2_(fold change) plotted against −log_10_(*p*-value). Molecular features were classified as significantly enriched in the CAA-present group (*p ≤* 0.05; *fold*-*change >* 1), significantly depleted (*p ≤* 0.05; *fold*-*change <* 1), or non-significant.

### 2.10 Supervised Machine Learning and Interpretation by Shapley Additive Explanations

To identify lipid species predictive of vascular identity and amyloid-associated pathology, we implemented the interpretable supervised machine learning workflow described in Tideman et al.[44] This approach uses XGBoost models to search for relationships between IMS-observed ion species and the microscopy-derived spatial mask of a tissue structure, cell type, or region of interest.[46] After training an XGBoost classification model with strong predictive performance, SHAP is used as a model interpretability method to determine which, out of the hundreds of ion species measured, are driving the model’s class prediction and, thus which molecular species hold marker potential for recognizing the tissue region of interest.[47][48]

Two classification tasks were defined: (1) distinguishing vasculature from background regions within the MALDI IMS ROI, and (2) differentiating CAA-present vasculature from CAA-absent vasculature based on immunofluorescence microscopy segmentation masks. These classification tasks were performed separately for both positive and negative ion mode datasets. Each MALDI IMS pixel, defined by its detected peak features (see Sec. 2.7), served as an input feature vector, while the spatial segmentation masks from collagen IV, labeled using αSMA+ or thiazine red+ threshold results, provided the class labels for model training.

For each classification task, we employed a one-versus-all strategy using an ensemble of 10 XGBoost models, each initialized with a different random seed. The training data for each model consisted of randomly sampled pixels from the relevant masks, with random under-sampling of the majority class (e.g., background or CAA-absent) and random over-sampling of the minority class (e.g., vasculature) to address class imbalance. Ensemble modeling helped reduce the risk of overfitting and ensured that biologically meaningful patterns were preserved while minimizing spurious correlations. Model performance was evaluated using out-of-sample predictions, and pixels not used during training were held out for testing. Specifically, 66% of the available samples were randomly assigned to the training set, while the remaining 34% were reserved as an independent test set. See Tab. S2 for the model performance metrics of each task.

To interpret the models trained for the different recognition scenarios, we employed SHAP to quantify the contribution of each ion species to the models’ predictions. For each molecular feature (ion), SHAP provided local importance scores (Shapley values) per pixel, and tissue-wide importance scores were calculated by averaging the local Shapley values across all pixels in the dataset. Final global SHAP scores for each ion were computed as the mean of the tissue-wide importance scores across the 10-model ensemble. These global SHAP scores were used to rank the IMS-reported features by predictive relevance to vascular classification or CAA status, and the top-ranked ions were considered candidate molecular markers. Annotation of these ion species was done manually after SHAP analysis using an LC-MS/MS library (see Sec. 2.8) and lipidmaps.org.[63][64] Final manual annotations are summarized in Tables S3 & S4.

To assess the relationship between lipid abundance and marker potential (*i.e.*, SHAP-based feature importance), we computed Spearman rank-order correlation coefficients (ρ) between ion intensity values and corresponding local Shapley values. This analysis was used to infer the directionality of a lipid’s association with a classification outcome. If this correlation is positive, increased presence of the lipid species in a pixel tends to increase the probability that this pixel is part of the tissue structure of interest (*i.e.*, the lipid species is a positive marker candidate). Negative correlation suggests that a decreased presence of the lipid species increases the probability that the tissue structure of interest is present (*i.e.*, the lipid species is a negative marker candidate). Overall, features with |ρ| > 0.2 were considered to show a meaningful monotonic relationship between ion intensity and classification probability. In visualizations such as summary bubble plots, the global SHAP score determined marker size, while Spearman correlation determined marker color, providing an interpretable view of both predictive importance and biological association.

### 2.11 MALDI IMS Automatic Identification

The average mass spectrum was scaled between 0 and 1, peak picked, deisotoped, and filtered using a signal-to-noise ratio threshold of 0.03 (calculated as the fraction of the base peak of the mass spectrum). Features below this threshold were excluded from the identification process. IMS identification was performed using in-house-developed software that associated detected peaks with tentative lipids from the LIPID MAPS Structure Database and custom LC-MS/MS database (discussed above). Parameters for annotating peaks include [M + H]^+^, [M + Na]^+^, and [M + K]^+^ adducts in positive mode and [M *−* H]*^−^* and [M *−* CH_3_]*^−^* in negative mode and a search window of *±*5 ppm. To provide more confident identification, IMS identification was subsequently associated with LC-MS/MS identifications.

## 3 Results

### 3.1 Multimodal Molecular Imaging Workflow Enables Vasculature Focused Analysis

To investigate spatial lipidomic signatures and changes associated with the vasculature and CAA features, we applied the multimodal molecular imaging workflow described in Fig. 1 to the leptomeninges in postmortem human frontal cortex tissue. We analyzed 13 cases with varying presence of AD and CAA pathology. Preliminary characterization of CAA was done according to the National Alzheimer’s Coordinating Center (NACC) (Tab. S5) to include sufficient control cases and cases with CAA pathology. CAA is staged according to the Vonsattel criteria defined by four grades: absent, mild, moderate, or severe. We viewed cases with moderate to severe CAA as having clinically relevant CAA.[49] For each case, we examined six serial sections - three technical replicates for negative ion mode MALDI IMS lipid imaging, and three technical replicates for positive ion mode imaging (Fig. 1a). Additional tissue sections were collected for downstream LC-MS/MS (Fig. 1e), to enable confident lipid identification and to support annotation of MALDI IMS peaks through the online database at lipidmaps.org.[63][64] Autofluorescence microscopy images acquired prior to MALDI IMS imaging (Fig. 1b) guided the selection of ROIs enriched for clearly visible leptomeninges, which are cortex-related structures particularly vulnerable to vasculature amyloid deposition, resulting in CAA (Fig. 2a, Figures S1-S3).[8][65][66][67] Four ROIs were removed in analysis due to technical difficulties. This workflow, using autofluorescence microscopy-directed MALDI IMS imaging [68] allowed us to expand our study across a large cohort of tissues.

**Figure 1.**
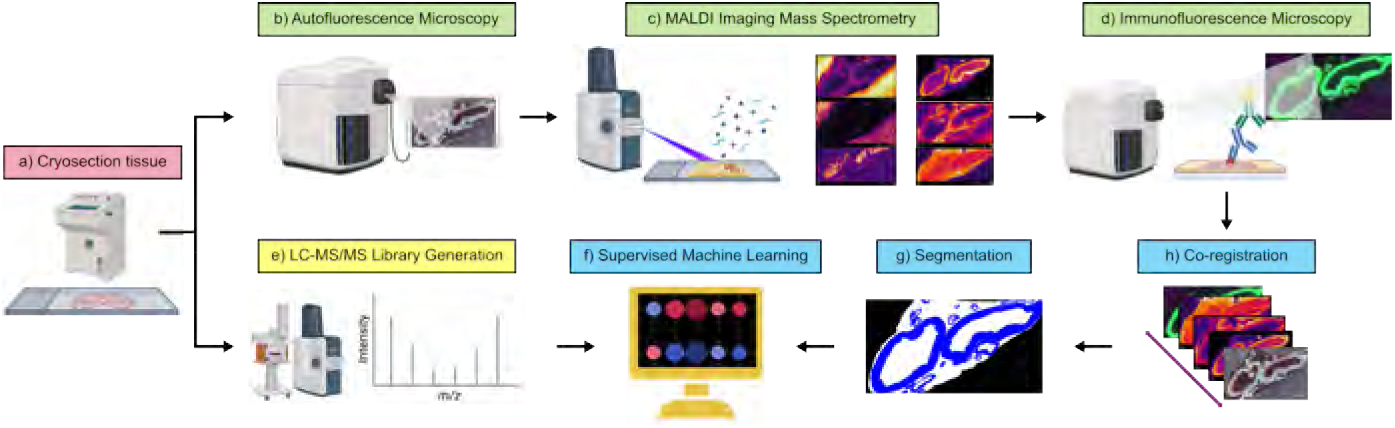
Multimodal Molecular Imaging Workflow. Sample preparation and tissue handling is shown in red. Data acquisition and imaging is shown in green. LC-MS/MS completed in parallel is shown in yellow. Data curation, integration, and analysis is shown in blue. (a) Post-mortem human frontal cortex tissue cryosectioned at 10 µm thickness. (b) Autofluorescence microscopy acquired on Zeiss AxioScan.Z1 to locate leptomeninges. (c) Matrix 4-aminocinnamic acid sublimated onto the samples (5 mg/ml) using an in-house developed sublimation device. IMS data acquired on a MALDI timsTOF flex mass spectrometer. (d) Matrix removed with ethanol washes. Antibody markers collagen IV, actin smooth muscle, and thiazine red stain were used to mark vasculature and pathogenic features. (e) Lipid IDs validated using in-house generated LC-MS/MS library and lipidmaps.org. (f) Peak alignment, mass calibration, intensity normalization, peak identification and supervised ML analysis. (g) Automated segmentation of leptomeninge vasculature, plus healthy and CAA-diseased regions. (h) Co-registration of IMS, autofluorescence and immunofluorescence microscopy data.

**Figure 2.**
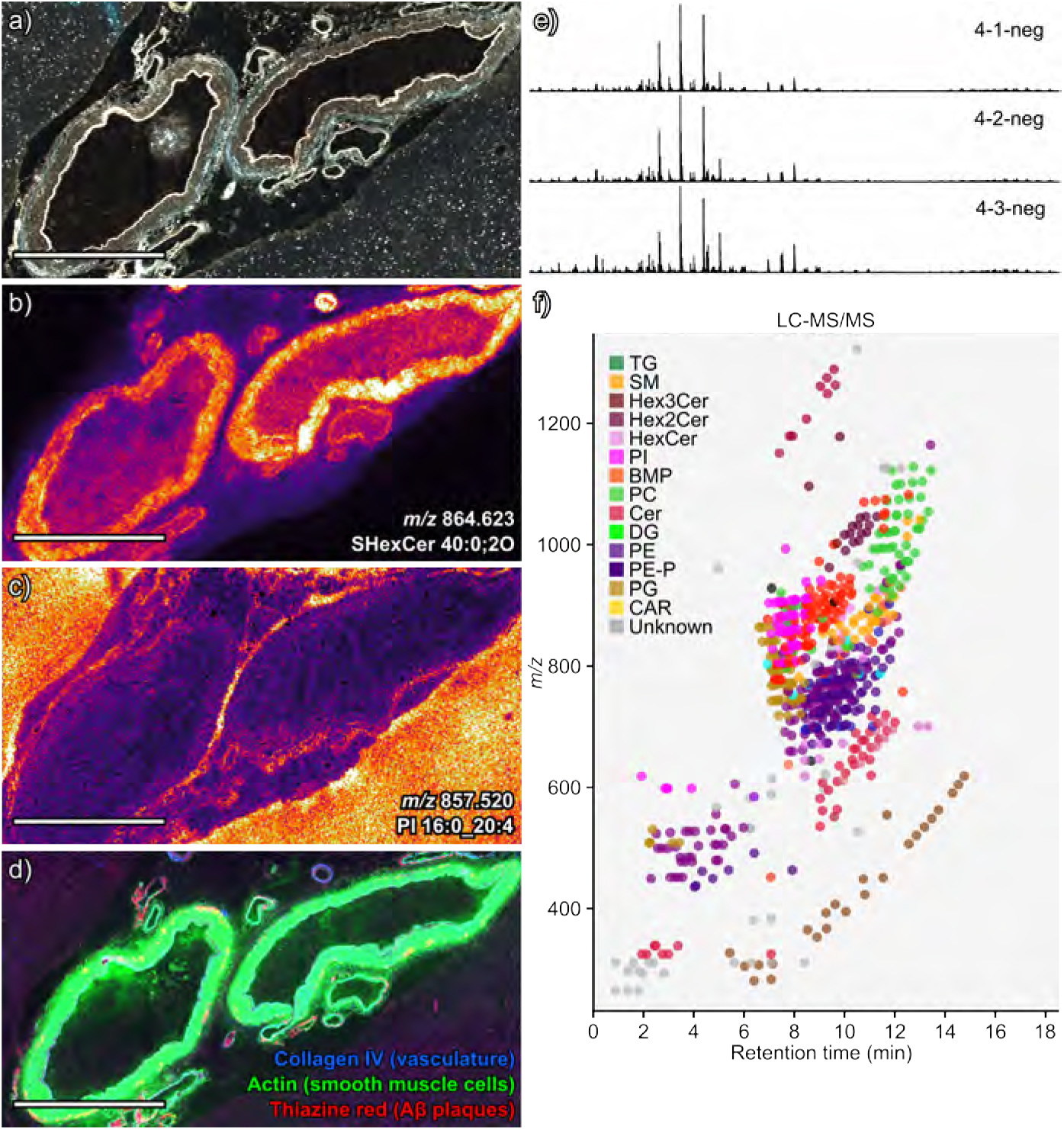
Data Acquisition. MALDI IMS coupled with immunofluorescence microscopy and LC-MS/MS were acquired on 13 donors with and without cerebral amyloid angiopathy (CAA) disease pathology. (a) Autofluorescence microscopy enables directed imaging of leptomeninges where CAA is most prominent. MALDI IMS heat maps of lipids (b) SHexCer 40:0;2O and (c) PI 16:0_20:4 are shown. (d) Immunofluorescence microscopy with collagen IV, smooth muscle cells, and amyloid markers was used to identify vasculature features and CAA. (e) The average spectrum of negative mode imaging data across 3 technical replicates for case 4. The spectral pattern of this MALDI IMS region (shown in a through d) is similar across three technical replicates. (f) Summary plot of lipid classes found from LC-MS/MS data captured on sections from all 13 cases shows a wide range of lipids within the sample. Scale bar is equal to 1 mm.

Overall, the MALDI IMS imaging dataset consisted of 74 ROIs (Fig. 1c). MALDI IMS at 10 µm pixel size enabled sensitive and reproducible lipid detection within the vasculature of the leptomeninges (Fig. 2b & 2c), preserving antigenicity for downstream immunofluorescence microscopy imaging on the same sections (Fig. 2d). Average mass spectra (Fig. 2e, Fig. S4) and Levey-Jennings plots (Fig. S5) display consistent signal to noise ratios and total ion currents across technical replicates. Automated IMS identification using in-house-developed software (see Methods-MALDI IMS Tentative Identification) detected 419 lipid features in negative ion mode (Fig. S6a), while another 537 lipids were detected in positive ion mode (Fig. S6b). Most of these molecular species exhibited distinct distributions within the brain tissue, including a subset that displayed spatial distributions matching vasculature patterns observed in the autofluorescence microscopy image (Fig. 2b & 2c). Notable lipids in negative mode such as SHexCer 40:0;2O (*m/z* 864.623) and PI 16:0_20:4 (*m/z* 857.520) showed clear localization to vasculature elements in the leptomeninges. Overall, the LC-MS/MS negative mode workflow confirmed the presence of at least 15 lipid classes within the dataset, including phosphatidylserines (PS), phosphatidylethanolamines (PE), ceramides (Cer), and inositol-containing lipids (PI), among others (Fig. 2f). The LC-MS/MS positive ion mode workflow confirmed the presence of an additional 14 lipid classes, including triglycerides (TG), sphingomyelins (SM), phosphatidylinositols (PI), and phosphatidylcholines (PC), among others (Fig. S7).

Following MALDI IMS, the same tissue sections were processed for immunofluorescence microscopy (Fig. 1d), allowing for segmentation of specific vasculature and CAA features (Fig. 1g). All image co-registration (Fig. 1h) was performed on modalities that measured the same tissue section. Co-registration of the MALDI IMS image with the post-IMS autofluorescence microscopy image was highly accurate (Fig. S8), enabling precise spatially resolved molecular analysis of structurally annotated vasculature and CAA features (Fig. 1f).

### 3.2 Automated Segmentation Enables Molecular Analysis of Healthy Vasculature and CAA-diseased Features

Combining immunofluorescence microscopy with MALDI IMS enables us to derive the presence or absence of CAA pathology from the microscopy and provide contextual information for spatial molecular data obtained via IMS. Collagen IV staining highlights the vascular basement membrane within the leptomeninges and serves as a broader marker of the total vasculature.[67] Smooth muscle actin (αSMA) was chosen to identify smooth muscle cells and areas of healthy vasculature. This builds on prior research suggesting that vasculature β-amyloid deposition occurs in a location similar to that of smooth muscle cells, potentially replacing the smooth muscle cell layer within the vasculature tissue in severely diseased vessels.[8][67] Collagen IV signal is continuously present across vasculature, while αSMA is variably expressed depending on vessel type and vasculature integrity.[67] Thiazine red, which binds β-sheet-rich amyloid aggregates, reveals regions of vascular amyloid deposition consistent with CAA pathology.[67] Representative immunofluorescence microscopy images show clear demarcation between amyloid-positive and amyloid-negative areas in the vasculature based on the location of thiazine red (Fig. 3a & 3b). This is critical for determining whether the vasculature across cases is CAA-present (diseased) or CAA-absent (healthy). This pathogenic heterogeneity evident in the immunofluorescence microscopy data allows for image segmentation and spatial lipidomic comparisons between diseased or healthy vascular tissue.

**Figure 3.**
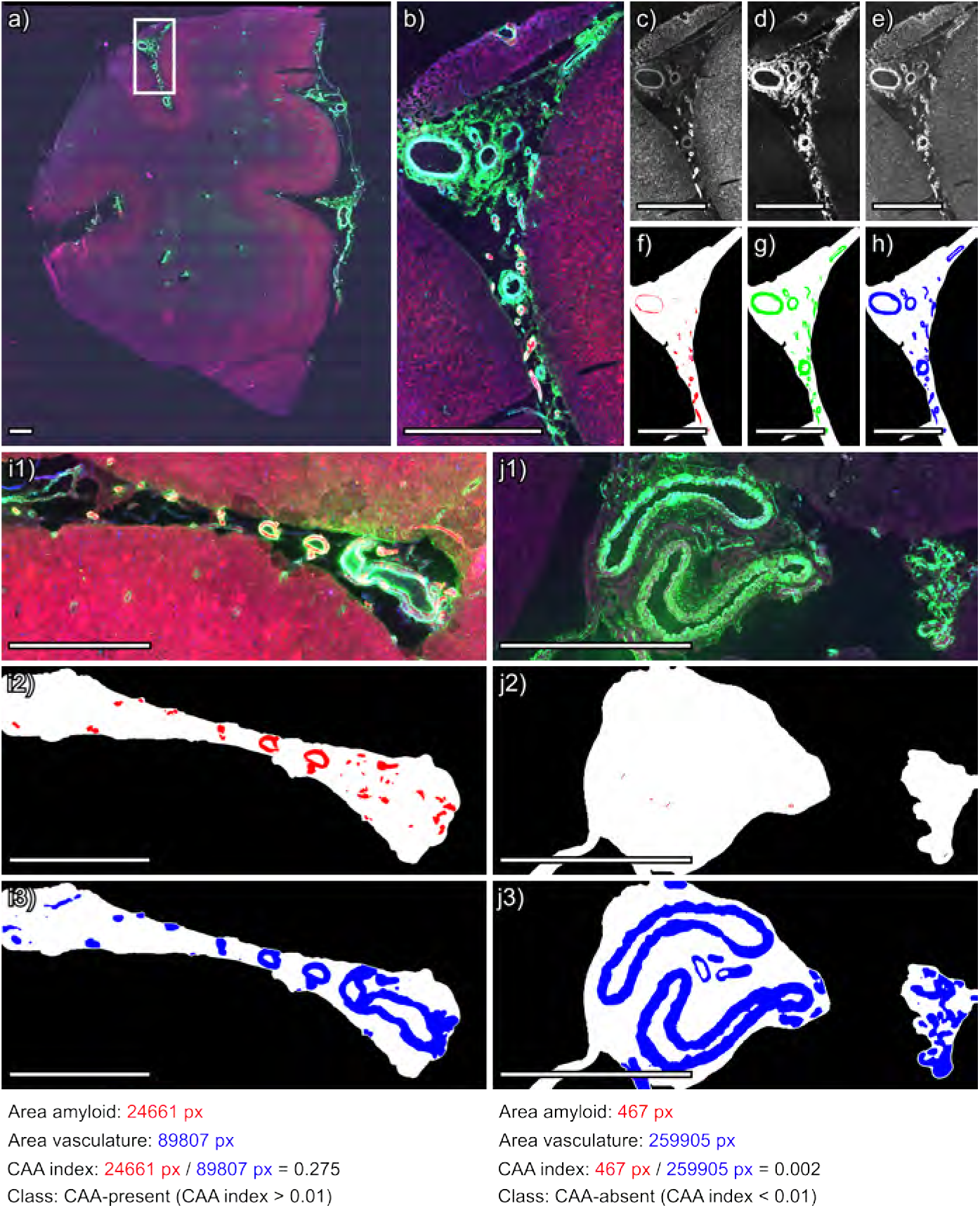
Workflow for CAA Index. (a) Whole slide immunofluorescence microscopy image. (b) Zoomed in immunofluorescence image of region acquired by MALDI IMS. (c) Thiazine red stains amyloid plaques, (d) smooth muscle actin stains smooth muscle, and (e) collagen IV stains the basement membrane of vasculature. Segmentation masks obtained for (f) amyloid, (g) smooth muscle cells, and (h) collagen IV. Below are examples of tissue sections above the CAA index (i1-i3) and below the CAA index (j1-j3). The IF image (i1 and j1), amyloid mask (i2 and j2) and collagen IV mask (i3 and j3) are shown. Calculated values for the area amyloid, the area vasculature, the CAA index (ratio of amyloid-positive/total vasculature), and the class (CAA-present above 0.01 CAA index and CAA-absent below 0.01) are shown below. Scale bar is equal to 1 mm.

To develop a robust method for automatically, rapidly, and reliably annotating the vasculature across a dataset, we first used the collagen IV signal (Fig. 3e) to define a broad vasculature mask (Fig. 3h). This was done using a custom spatial segmentation method (see Methods-Tissue Feature Annotations). Within this vasculature mask, we utilized the thiazine red marker (Fig. 3c) and the actin smooth muscle cell marker (Fig. 3d) to define two additional regions using a thresholding segmentation method (see Methods-Tissue Feature Annotations). Taken together, we defined a total of three regions: total vasculature (based on collagen IV) (Fig. 3h), amyloid-positive vasculature (based on thiazine red+ only) (Fig. 3f), and amyloid-negative vasculature (based on αSMA+ only) (Fig. 3g). All immunofluorescence microscopy images and segmentation results for the cohort are available in the supplement (Figures S9-S82). Because CAA is a pathology defined by a complex vasculature microenvironment,[67] we chose not to restrict our search to tissue that directly colocalized with amyloid deposits. Instead, we focused our analysis on molecular data within the broad vasculature, as defined by collagen IV, using information from its amyloid plaque area to classify the entire vasculature mask as either CAA-present (a more CAA-diseased region) or CAA-absent (a healthier region). To accomplish this, we define a CAA index based on the ratio of amyloid plaque area to total vasculature area per each IMS ROI in the section. A threshold of 0.01 (amyloid-positive vasculature area/total vasculature area), representing the ratio of amyloid plaque area to collagen IV vasculature area, was determined to be optimal for defining CAA within the vasculature mask based on comparisons with traditional pathological classification and input from expert neuropathologists. For example, Fig. 3i shows a vascular tissue area with a high CAA index. The first image shows immunofluorescence microscopy (Fig. 3i1) with a high degree of amyloid. Below it is an image showing the results of the generated amyloid mask (Fig. 3i2), which has a relatively high area overlapping with the total collagen IV mask (Fig. 3i3). That example’s CAA index was calculated as 0.275, well above the established 0.01 threshold. Therefore, this section was defined as CAA-present. Figure 3j shows a vascular tissue area with a lower CAA index. The first image shows immunofluorescence microscopy (Fig. 3j1) with little to no amyloid present. Below it is an image showing the results of the generated amyloid mask (Fig. 3j2), which has a low area overlapping with the collagen IV mask (Fig. 3j3). The CAA index for this example was calculated as 0.002, well below the 0.01 threshold, and was therefore defined as CAA-absent.

This workflow was repeated for each of the six sections acquired per case. The raw CAA index values for each ROI across the entire dataset can be visualized in the CAA index heat map (Fig. S83a). All ROIs with index values above the threshold of 0.01 were categorized as CAA-present, while all sections below the threshold were categorized as CAA-absent (Fig. S83b). The average value across the sections was used to assign a case to the CAA-present or CAA-absent group. In total, 7 cases were categorized as CAA-present, and 6 as CAA-absent. The final grouping is shown in Tab. S6. The categorizations for each case are also in agreement with the preliminary characterization by NACC (Tab. S5).

Importantly, the CAA index is not intended to quantify disease burden within each IMS ROI. Because CAA severity exists on a continuum, the index is used to classify each vasculature mask as relatively more CAA-like or non-CAA-like. This grouping enables direct downstream comparisons, particularly with binary classification models, even if the index does not precisely measure disease severity.

### 3.3 Molecular Signatures Differentiate CAA-present and CAA-absent Vasculature

With the sections classified as diseased or healthy, we can now search for a link between these labels and corresponding IMS observations, i.e., we can pursue identification of disease-associated lipid changes. We first performed a traditional univariate comparison between MALDI IMS ion intensities that stem from CAA-present or CAA-absent vasculature segments. We specifically examined ion intensity within the vasculature segmentation mask generated by collagen IV, thereby excluding confounding signal contributions from the perivascular space or adjacent grey and white matter. Since both CAA-present and CAA-absent segments represent the same cell types in the leptomeninges, we anticipated substantial overlap in their molecular profiles. Because we separated the two groups of similar vasculature tissue strictly by their ratio of amyloid-positive/total vasculature (see Sec. 3.2), any observed differences would be expected to reflect lipid features that positively or negatively associate with amyloid deposition rather than cell-type composition.

Group-level molecular differences were analyzed using a difference spectrum derived from subtraction of the mean normalized spectra of CAA-absent vasculature from CAA-present vasculature across all *m/z* features (Fig. S84a & S84b). In these diagrams, mass spectra shown in blue represent ions with higher intensity in the average mass spectrum of the entire CAA-absent vasculature mask group, while mass spectra in orange represent ions with higher intensity across the average mass spectrum of the entire CAA-present group. These differences were subtle, which is consistent with the expectation that amyloid-associated changes occur against a largely conserved vascular lipid profile.

We also accounted for inter-donor variability, extending this analysis to donor-level univariate comparisons using average spectra representing CAA-present and CAA-absent vascular regions from individual donors. Across both negative (Fig. S85) and positive (Fig. S86) ion modes, these comparisons similarly revealed only marginal differences between CAA-present and CAA-absent vasculature. Collectively, these results suggest that CAA-associated lipid changes are not dominated by large, isolated abundance changes in individual molecules, but instead may manifest as more subtle multivariate shifts in the vascular lipidome as molecules respond in concert to amyloid deposition.

To capture these multivariate patterns, we transitioned from univariate analysis to an interpretable machine learning framework that combines classification by eXtreme Gradient Boosting (XGBoost) and Shapley Additive exPlanations (SHAP).[44][45][46][47][48] XGBoost models integrate information from many ions simultaneously, enabling multivariate classification of vascular regions with or without CAA. SHAP complements this approach by quantifying the contribution of each ion species to the model predictions, thereby identifying lipid features whose combined effects most strongly distinguish CAA from healthy vasculature.

As an initial validation of this approach, we trained an XGBoost classifier to distinguish IMS pixels within the vasculature mask from those in the surrounding background tissue (i.e., non-vasculature regions). The performance metrics for the model can be seen in Tab. S2. This analysis established lipid profiles that broadly define vascular tissue and that differentiate it from non-vascular tissue. We initially compared the vasculature masks from the CAA-present group with the corresponding background masks (Fig. S87a). The top-ranked lipid features that drove the classification model to differentiate between CAA-present vasculature and other surrounding tissue regions are shown in Fig. S87b. The ions *m/z* 864.623 (SHexCer 40:0;2O) and *m/z* 1027.687 (unknown) ranked among the top differentiating features and were also found to have significantly higher intensity within the univariate comparison (Fig. S85b).

We performed a comparable analysis for the CAA-absent vasculature group (Tab. S2, Fig. S88a), which revealed a similar set of lipid marker candidates (Fig. S88b). Ion images for the key lipids found to define vasculature in general (i.e., both CAA-present and CAA-absent vasculature) are shown in Fig. 4. The substantial overlap in lipid identifications between these two analyses highlights the strong molecular similarity between CAA-present and CAA-absent vessels. These shared molecular species most likely correspond to structural lipids necessary for vasculature integrity. This is expected since the primary driver here is the difference between the cell types of the vasculature (e.g., endothelial and vascular smooth muscle cells) and the other cell types of the brain. Looking beyond the top-ranked features, we do see more well-defined changes when comparing features with lower SHAP scores. For example, *m/z* 788.545 (PS 18:0_18:1) seems to be a stronger marker candidate for the CAA-absent group compared to CAA-present vasculature (Fig. S89).

**Figure 4.**
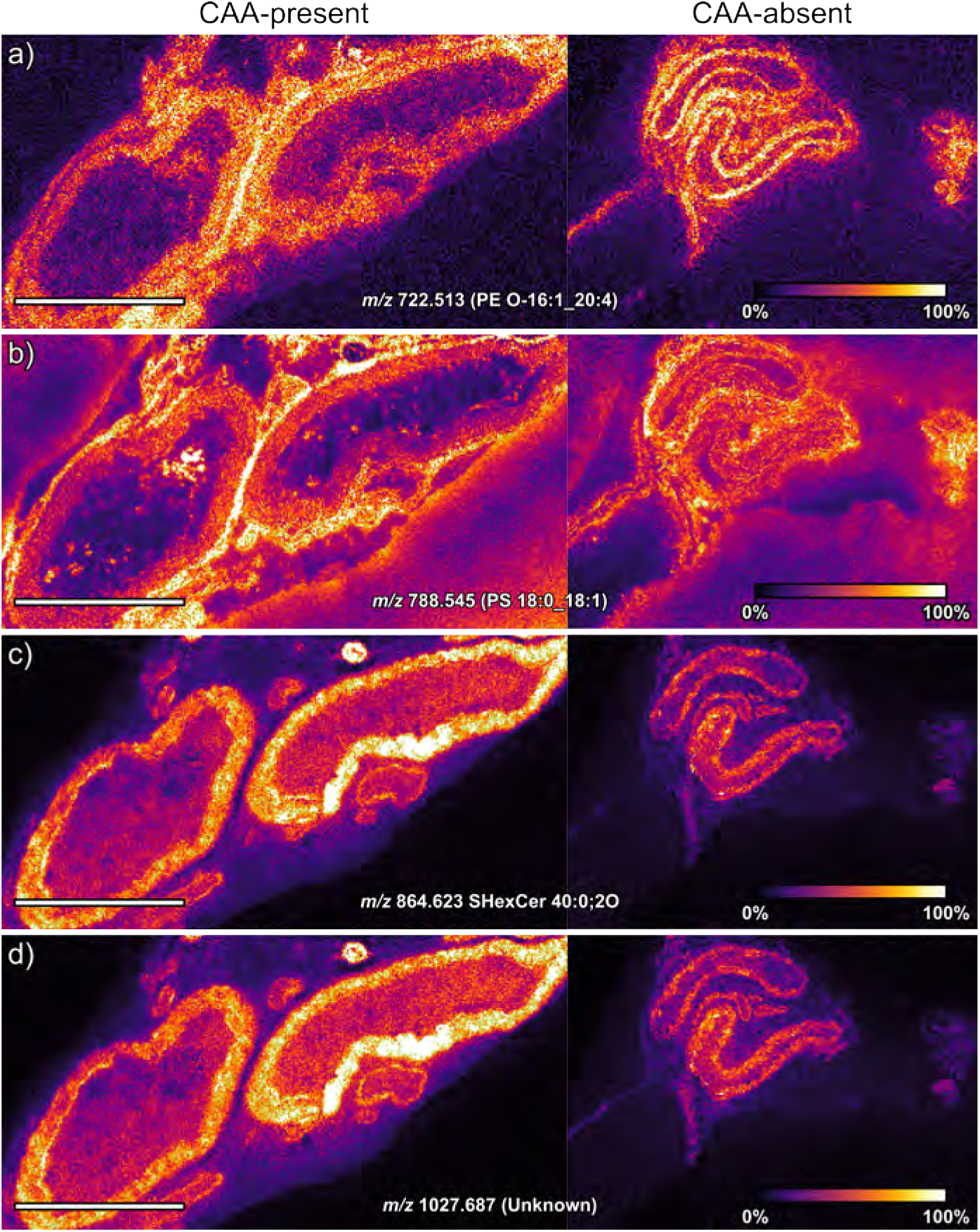
Molecular Marker Candidates for Vascular in General. Interpretable machine learning using SHAP on negative mode data identified markers that map the vasculature. Top hits are shown, including (a) *m/z* 722.513 (PE O-16:1_20:4), (b) *m/z* 788.545 (PS 18:0_18:1), (c) *m/z* 864.623 SHexCer 40:0;2O, (d) *m/z* 1027.687 (Unknown). Scale bar is equal to 1 mm.

Next, we performed the same task on the positive ion mode dataset cohort. When training a classifier to differentiate CAA-present vasculature masks from non-vasculature background masks (Tab. S2, Fig. S90a), we identified several lipid marker candidates that positively correlated with vasculature (Fig. S90b). This workflow was repeated to differentiate between CAA-absent vasculature masks and non-vasculature background masks (Tab. S2, Fig. S91a), revealing a similar set of top-ranked lipid marker candidates (Fig. S91b). Overlapping marker candidates between CAA-present and CAA-absent vasculature, such as *m/z* 703.575 (SM 16:1;O2/18:0), 705.584 (SM 34:0;O2), *m/z* 701.559 (SM 20:2;O2/14:0), *m/z* 766.574 (PE P-17:0_22:4), and *m/z* 741.530 (Unidentified), most likely play key roles in vasculature structure and integrity, similar to the overlapping marker candidates in the negative mode dataset.

To determine lipidomic differences specifically between vessels with amyloid and healthy vessels, we performed a second type of classification task that was limited to pixels associated with vascular masks. MALDI IMS data associated with CAA-present and CAA-absent vasculature were used to train a classification model to distinguish between the two conditions in both negative and positive ion modes, while ignoring non-vascular related pixels (see Tab. S2 for performance metrics).

The negative ion mode analysis revealed a set of features with high-ranking SHAP scores, correlating either with CAA-present tissue regions (in red) or CAA-absence (in blue) (Fig. 5a & 5b). Molecular features that were positively correlated to CAA-present vessels include *m/z* 1544.867 (GM1 36:1;O2), *m/z* 774.544 (PE O-18:1_22:6), *m/z* 834.529 (PS 18:0_22:6), *m/z* 568.268 (LPS 22:6), *m/z* 760.514 (PS 16:0_18:1), and *m/z* 746.513 (PE O-16:1_22:6), with *m/z* 1544.867 (GM1 36:1;O2) standing out in terms of marker potential. Molecular features that were positively correlated to CAA-absent vessels include *m/z* 838.559 (PS 18:0_22:4), *m/z* 836.539 (PS 18:0_22:5), *m/z* 888.624 (SHexCer 42:2;2O), *m/z* 883.533 (PI 18:1_20:4), 786.529 (PS 18:1_18:1), *m/z* 788.545 (PS 18:0_18:1), and *m/z* 794.570 (PE 18:0_22:4). Interestingly, PS 18:0_22:4 and PS 18:0_22:5 had the largest global SHAP importance scores, indicating that out of all molecular features measured in these experiments these two lipids hold the most potential to serve as markers for distinguishing healthy and diseased vasculature. This suggested relationship was confirmed in their localizations, which correlate distinctly to CAA-absent vascular regions. When examining the individual ion images, these lipids show uniform high intensity throughout the grey matter regions across all cases, but more focused analysis within the vascular regions makes it clear that these lipids are found with greater intensity and specific localization to vasculature elements in the CAA-absent cases compared to the CAA-present cases (Fig. 5c). All negative mode annotations can be seen in Tab. S3.

**Figure 5.**
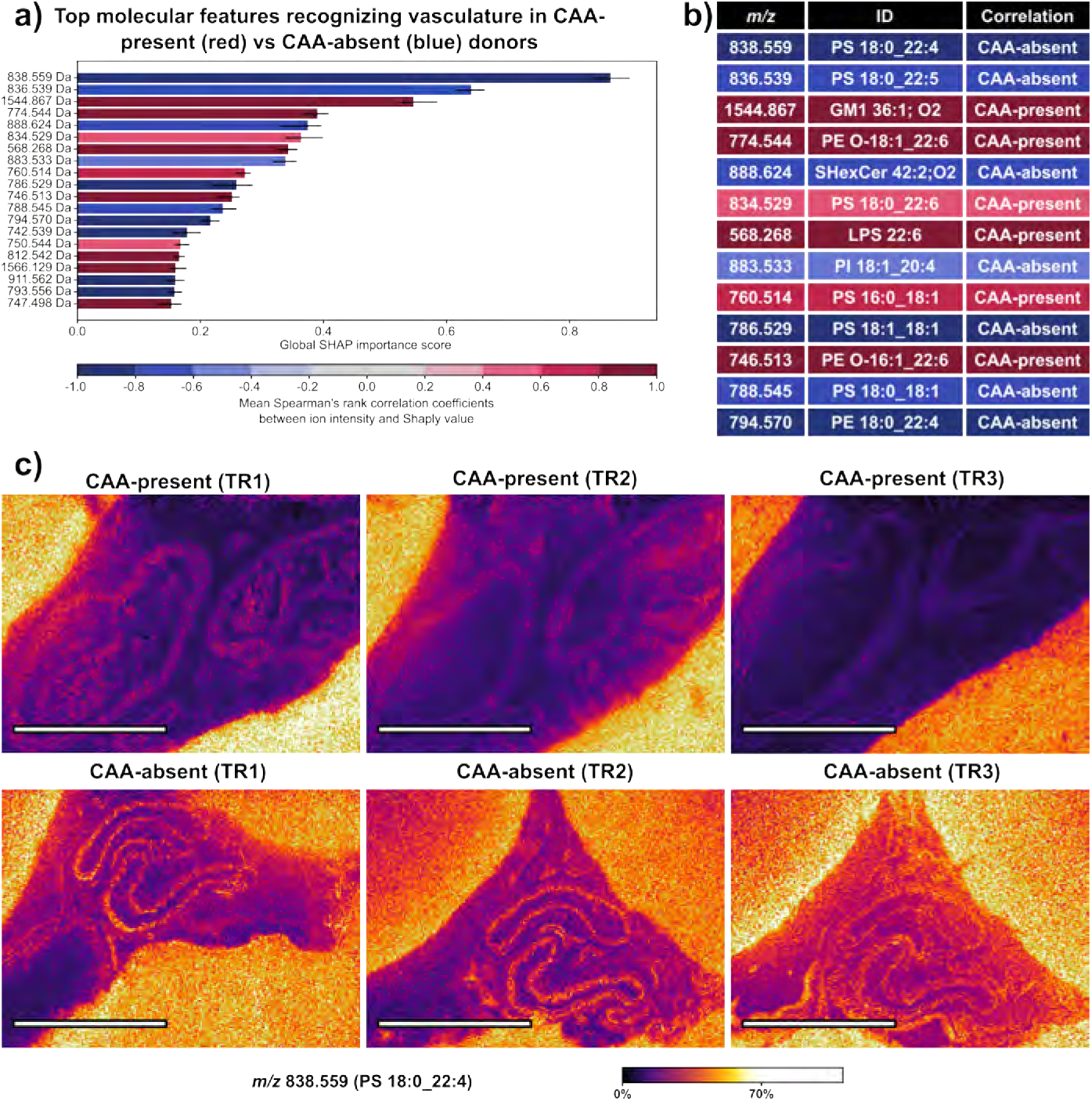
Molecular Marker Candidates for CAA. (a) Ranked list of molecular features distinguishing between CAA-present (red) and CAA-absent (blue) vasculature masks. (b) Ranked list of molecular features and corresponding lipid annotations. (c) Ion image of top-ranked lipid, *m/z* 838.559 (PS 18:0_22:4), comparing three replicates of CAA-present tissue (above) and CAA-absent tissue (below), with higher intensity in CAA-absent vasculature. Scale bar is equal to 1 mm.

Lipid features in positive ion mode that correlate to CAA-present vasculature included *m/z* 731.606 (SM 36:1;O2), *m/z* 729.590 (SM 18:1;O2/18:1), *m/z* 812.611 (PC 38:3), *m/z* 798.541 (unknown), *m/z* 703.575 (SM 16:1;O2/18:0), and *m/z* 787.688 (SM 40:1;O2) (Tab. S2, Fig. S92). Lipids *m/z* 786.600 (PE 39:2) and 759.638 (SM 20:1;O2/18:0) correlated to CAA-absent vasculature (Tab. S2, Fig. S92). Overall, in positive ion mode, our analysis did not find standout molecular features with clear marker potential. All positive mode annotations can be seen in Tab. S4.

## 4 Discussion

CAA is a common yet understudied vascular pathology that is closely associated with Alzheimer’s disease, and the molecular mechanisms linking β-amyloid deposition to vasculature degeneration remain poorly understood.[8][9][34][66][69][70] In this study, we employed a multimodal molecular imaging approach to investigate the spatial lipidomic profiles of vasculature in postmortem human cortex from cases with and without CAA pathology. By integrating MALDI IMS with immunofluorescence microscopy and applying our SHAP-based interpretable supervised machine learning pipeline, we identified distinct lipid signatures associated with CAA-present vasculature and CAA-absent vasculature (Fig. 1). These findings reveal new insights into the molecular microenvironment of CAA and highlight lipid dysregulation as a key feature of vascular amyloid pathology.

Our use of autofluorescence microscopy-guided MALDI IMS allowed for precise imaging of leptomeningeal vessels, a niche particularly vulnerable to CAA pathology. High-resolution lipid imaging across 13 cases captured 956 total lipid features (Fig. S6), demonstrating broad lipidomic coverage within these vascular structures. Importantly, our imaging strategy preserved tissue antigenicity, enabling downstream immunofluorescence microscopy and the segmentation of vasculature into vasculature regions as well as into amyloid-positive and amyloid-negative subsets (Fig. 3). Highly precise multimodal image co-registration enabled lipid signals to be assigned to discrete vascular features, which is an advantage over traditional bulk lipidomics and non-spatial omics techniques.[22] Automated vasculature segmentation using the Moran quadrant map, a method developed by Tideman et al.,[60] allowed for accurate segmentation of vasculature tissue features. Combined with thiazine red staining, this enabled robust binary classification of vessels based on amyloid presence, and it allowed us to define a quantitative CAA index.

By more precisely focusing our analysis on vasculature-specific ROIs, we further refined our assessment of lipid changes associated with CAA pathology. The specificity of our approach allowed us to implement the interpretable supervised machine learning method described in Tideman et al.[44] This involved training an XGBoost classification model to differentiate CAA-present samples from CAA-absent samples, using only the IMS-measured lipid species as inputs. Once the model successfully differentiated, SHAP was used to probe the decision process of the model, which in turn allowed us to rank all lipid features according to their relevance in differentiating. This implicitly identified potential marker candidates for both vascular identity and pathology, while minimizing confounding chemical signals from grey matter, white matter, perivascular space in the leptomeninges, or other regions of the frontal cortex. Classifiers trained to distinguish vasculature from surrounding tissue identified consistent lipid marker candidates of vascular structure across human donors, such as *m/z* 864.623 (SHexCer 40:0;2O) and *m/z* 703.575 (SM 16:1;O2/18:0). Notably, some of these marker candidates, particularly in negative ion mode, showed reduced predictive strength in CAA-present vasculature compared to CAA-absent vasculature, suggesting that core structural lipids may be diminished or redistributed during CAA-associated vascular degeneration.

When classifiers were trained to distinguish CAA-present from CAA-absent vasculature, a distinct lipid signature was suggested. In amyloid-absent vessels, we observed enrichment of certain phosphatidylserines, *m/z* 838.559 (PS 18:0_22:4) and *m/z* 836.539 (PS 18:0_22:5) (Fig. 5). We also saw *m/z* 788.545 (PS 18:0_18:1) lose marker candidate strength in the CAA-present vasculature compared to CAA-absent vasculature when classifying against the background(Fig. S89), suggesting that its association with vasculature might be selectively weakened in the presence of amyloid. Phosphatidylserines are known to contribute to membrane curvature, apoptosis signaling, and coagulation pathways, and their loss may reflect early lipidomic alterations that predispose vessels to amyloid deposition or reflect loss of vascular integrity.[71][72][73] Conversely, CAA-present vasculature showed increased abundance of GM1 gangliosides (GM1 36:1;O2). This lipid has been previously implicated in amyloid pathology and oxidative stress.[70],[74][75][76][77][78][79][80][81] This proposes a potential role of GM1 in vascular amyloid deposition and supports the idea that some overlapping lipid-mediated mechanisms may contribute to both neuritic and vascular β-amyloid aggregation.[82]

Together, these findings provide molecular context for prior histological observations of vascular smooth muscle cell loss and extracellular matrix remodeling in CAA.[20][67][83][84] The altered abundance of phosphatidylserines may underlie the degeneration of vascular integrity, while the accumulation of glycosphingolipids may either promote amyloid aggregation or arise as a consequence of CAA. Our findings align with genomic associations linking CAA risk and APOE ε4, a key regulator of lipid metabolism, and with prior bulk lipidomic analyses linking lipid dysregulation to cerebrovascular disease.[85][86][87][88] Unlike these prior studies, however, our work provides spatially resolved molecular profiles at the level of individual vessels, enabling direct correlation of molecular features with histopathological features of CAA.

Finally, the integration of spatial lipidomics with immunofluorescence microscopy-based segmentation provides a platform for future investigation of other pathological processes in CAA, including inflammatory cell infiltration, matrix metalloproteinase activity, and perivascular clearance mechanisms. Future studies incorporating spatial transcriptomics, proteomics, or other highly multiplexed imaging modalities (e.g., co-detection by indexing immunofluorescence microscopy) could further link lipid alterations to specific cell types and molecular pathways.[89][90][91] Moreover, the observed heterogeneity across cases suggests that lipid alterations in CAA may vary with genetic background, comorbid pathology, or treatment history, which warrants further investigation in larger, clinically annotated cohorts. Lastly, while accurate mass and LC-MS/MS confirmation support lipid assignments, some molecular features remain unannotated, and functional validation of lipid candidates will be necessary to determine their roles in CAA pathogenesis.

## 5 Author Contributions

C.R.M. and F.A.M. contributed equally to this work and led the study. C.R.M. conceived the project, performed the experiments, curated and analyzed the data, and wrote the manuscript. F.A.M. contributed to project conceptualization, data analysis, and manuscript writing. C.F.S. contributed to study conceptualization. L.V.-A. and W.R.-F. contributed to conceptualization, experimental design, and provided methodological guidance and domain expertise. L.G.M. and L.E.M.T. contributed to data analysis and the development of analytical methods. M.E.C. acquired experimental data and contributed to the Methods section. M.D. contributed to methodological development and experimental methods. M.S.S. contributed to study design and experimental planning and provided clinical and pathological expertise. R.V.d.P. contributed to data analysis and study conceptualization. J.M.S. supervised the study, contributed to manuscript writing, and secured funding. All authors reviewed, edited, and approved the final manuscript.

## 6 Acknowledgements

This work was supported by grants of the National Institutes of Health (NIH)’s National Institute on Aging (R01AG078803), National Institute of Diabetes and Digestive and Kidney Diseases (U54DK134302 and U01DK133766), and National Cancer Institute (U01CA294527), awarded to JMS. It was furthermore supported in part by grant numbers 2021-240339 and 2022-309518 (LGM and RV) of the Chan Zuckerberg Initiative DAF, an advised fund of Silicon Valley Community Foundation. CRM was supported by an Interdisciplinary Alzheimer’s Disease T32 training grant (5T32AG058524-07). The content is solely the responsibility of the authors and does not necessarily represent the official views of the National Institutes of Health.

## 7 Conclusion

In conclusion, we have developed a multimodal molecular imaging framework that integrates spatially resolved lipid IMS with fluorescence microscopy and interpretable supervised machine learning to enable vessel-specific molecular profiling of cerebral amyloid angiopathy in the human brain. Using a custom, quantitative CAA index consistent with neuropathological assessment, this approach directly links lipid profiles to leptomeningeal vessels with and without CAA. Our findings demonstrate that CAA is associated with coordinated lipid remodeling rather than isolated molecular changes, characterized by reduced phosphatidylserine levels and enrichment of glycosphingolipids within diseased vessels. These vasculature lipid signatures provide unique insights into lipid metabolism, vascular degeneration, and amyloid deposition, which cannot be captured adequately by bulk or univariate analyses. While larger cohorts and complementary spatial omics will be required to establish causality and clinical relevance, this work establishes spatial lipidomics as a powerful platform for interrogating cerebrovascular pathology and offers a molecular toolbox for uncovering lipidomic features of CAA in Alzheimer’s disease.

