## Supplementary Material for "Multimodal Molecular Mapping of the Vasculature in Human Cortex Reveals Lipid Markers of Cerebral Amyloid Angiopathy"

| | Negative Ion Mode | Positive Ion Mode |
|---|---|---|
| **Transfer** | | |
| MALDI Plate Offset | -70.0 V | 70.0 V |
| Deflection Delta Sum | -120.0 V | 120.0 V |
| Funnel 1 RF | 350.0 Vpp | 350.0 Vpp |
| IsCID Energy | 0.0 eV | 5.0 eV |
| Funnel 2 RF | 400.0 Vpp | 300.0 Vpp |
| Multipole RF | 500.0 Vpp | 400.0 Vpp |
| **Collision Cell** | | |
| Collision Energy | -10.0 eV | 10.0 eV |
| Collision RF | 3000.0 Vpp | 2000.0 Vpp |
| **Quadrupole** | | |
| Ion Energy | -5.0 eV | 5.0 eV |
| Low Mass | $m/z$ 500.00 | $m/z$ 400.00 |
| **Focus Pre TOF** | | |
| Transfer Time | 120.0 µs | 70.0 µs |
| Pre Pulse Storage | 12.0 µs | 10.0 µs |

**Table S1** Instrument parameters for negative and positive ion mode qTOF MALDI IMS experiments.

| Polarity | Classification | Balanced accuracy | Precision | Recall | F1 |
|---|---|---|---|---|---|
| Positive | CAA-present vs rest | 0.960 ± <0.001 | 0.941 ± 0.001 | 0.950 ± 0.001 | 0.945 ± 0.001 |
| Positive | CAA-absent vs rest | 0.965 ± <0.001 | 0.941 ± 0.001 | 0.948 ± 0.001 | 0.945 ± 0.001 |
| Positive | CAA-present vs CAA-absent | 0.994 ± <0.001 | 0.996 ± <0.001 | 0.995 ± <0.001 | 0.995 ± <0.001 |
| Negative | CAA-present vs rest | 0.954 ± <0.001 | 0.943 ± 0.001 | 0.937 ± 0.001 | 0.940 ± 0.001 |
| Negative | CAA-absent vs rest | 0.952 ± <0.001 | 0.933 ± 0.001 | 0.923 ± 0.001 | 0.928 ± 0.001 |
| Negative | CAA-present vs CAA-absent | 0.982 ± <0.001 | 0.987 ± 0.001 | 0.983 ± 0.001 | 0.985 ± 0.001 |

**Table S2** Performance metrics for each classification task. Each classifier consisted of an ensemble of 10 XGBoost models, each initialized with a different random seed. Reported metrics were calculated from out-of-sample predictions and are presented as mean ± standard deviation across ensemble members.

| IMS $m/z$ | Theoretical $m/z$ | IMS ppm error | Identification |
|---|---|---|---|
| 568.268 | 568.2681 | -0.176 | LPS 22:6* |
| 619.2895 | 619.2889 | 0.969 | LPI 20:4 |
| 722.513 | 722.513 | 0.0 | PE O-16:1_20:4 |
| 746.513 | 746.513 | 0.0 | PE O-16:1_22:6 |
| 760.514 | 760.5134 | 0.789 | PS 16:0_18:1 |
| 774.544 | 774.5443 | -0.387 | PE O-18:1_22:6 |
| 786.529 | 786.5291 | -0.127 | PS 18:1_18:1 |
| 788.545 | 788.5447 | 0.380 | PS 18:0_18:1 |
| 790.541 | 790.5366 | 5.566 | PE 18:0_22:6 |
| 794.570 | 794.5705 | -0.629 | PE 18:0_22:4 |
| 834.529 | 834.5291 | -0.120 | PS 18:0_22:6 |
| 836.539 | 836.5447 | -6.814 | PS 18:0_22:5 |
| 838.559 | 838.5604 | -1.670 | PS 18:0_22:4 |
| 857.520 | 857.5186 | 1.633 | PI 16:0_20:4 |
| 864.623 | 864.624 | -1.157 | SHexCer 40:0;O2 |
| 866.633 | N/A | N/A | Unidentified |
| 883.533 | 883.5342 | -1.358 | PI 18:1_20:4 |
| 888.624 | 888.624 | 0 | SHexCer 42:2;O2 |
| 973.722 | N/A | N/A | Unidentified |
| 1027.687 | N/A | N/A | Unidentified |
| 1029.696 | N/A | N/A | Unidentified |
| 1544.867 | 1544.8694 | -1.553 | GM1 36:1;O2* |

**Table S3** List of selected molecules highlighted in this work that were detected in negative ion mode. Asterisks indicate molecules not validated by LC-MS/MS.

| IMS *m/z* | Theoretical *m/z* | IMS ppm error | Identification |
|---|---|---|---|
| 701.559 | 701.5592 | -0.285 | SM 20:2;O2/14:0 |
| 703.575 | 703.5748 | 0.284 | SM 16:1;O2/18:0 |
| 705.584 | 705.5793 | 6.661 | SM 34:0;O2 |
| 729.590 | 729.5905 | -0.685 | SM 18:1;O2/18:1 |
| 731.606 | 731.6061 | -0.137 | SM 36:1;O2 |
| 734.569 | 734.5694 | -0.545 | PC 16:0_16:0 |
| 741.530 | N/A | N/A | Unidentified |
| 759.638 | 759.6374 | 0.790 | SM 20:1;O2/18:0 |
| 766.574 | 766.5745 | -0.652 | PE P-17:0_22:4 |
| 786.600 | 786.6007 | -0.890 | PE 39:2 |
| 787.668 | 787.6687 | -0.889 | SM 40:1;O2 |
| 798.541 | N/A | N/A | Unidentified |
| 812.611 | 812.6164 | -6.645 | PC 38:3 |

**Table S4** List of selected molecules highlighted in this work that were detected in positive ion mode. Asterisks indicate molecules not validated by LC-MS/MS.

| Cases | Age at death / Sex | Primary clinical categorization | AD Neuropathology | CAA grade | PMI | Other neuropathologies |
|---|---|---|---|---|---|---|
| Case 1 | 81/F | Alzheimer's disease | Severe (A3, B3, C3) | Severe CAA | 35H53' | NONE |
| Case 2 | 68/F | Alzheimer's disease | Severe (A3, B3, C3) | Severe CAA | 9h | None |
| Case 3 | 68/F | Alzheimer's disease | Severe (A3, B3, C3) | Low CAA | 15h | None |
| Case 4 | 70/M | Alzheimer's disease | Severe (A3, B3, C3) | Severe CAA | 11h | Tau in middle and superior temporal gyrus |
| Case 5 | 71/M | Alzheimer's disease | Severe (A3, B3, C3) | Severe CAA | 24h | Hemorrhages |
| Case 6 | 66/F | Alzheimer's disease | Severe (B5, Thal 4, C3) | Mild CAA | 25h | Small subdural |
| Case 7 | 73/M | Alzheimer's disease | Severe (A3, B3, C3) | Moderate CAA | 7h55' | None |
| Case 8 | 68/M | Alzheimer's disease | Moderate (A3, B2, C3) | Mild CAA | 18h40' | FTD TDP43, 4R-tau grains |
| Case 9 | 68/M | Neurological control | None | None | 24h | None |
| Case 10 | 66/M | Neurological control | None | None | 23h38' | None |
| Case 11 | 75/F | Alzheimer's disease | Severe (A3, B3, C2) | Mild CAA | 6h36' | Moderate vascular disease |
| Case 12 | 73/M | Alzheimer's disease | Moderate (A3, B2, C2) | None | 15h | LBD, stroke |
| Case 13 | 70/F | Neurological control | None | None | 35h15' | ARTAG |

**Table S5** Preliminary pathology characterization by the National Alzheimer's Coordinating Center (NACC).

| **CAA-present** | **CAA-absent** |
|---|---|
| Case 1 | Case 2 |
| Case 3 | Case 6 |
| Case 4 | Case 9 |
| Case 5 | Case 10 |
| Case 7 | Case 12 |
| Case 8 | Case 13 |
| Case 11 | |

**Table S6** List of donors labeled as CAA-present and CAA-absent according to our CAA index.

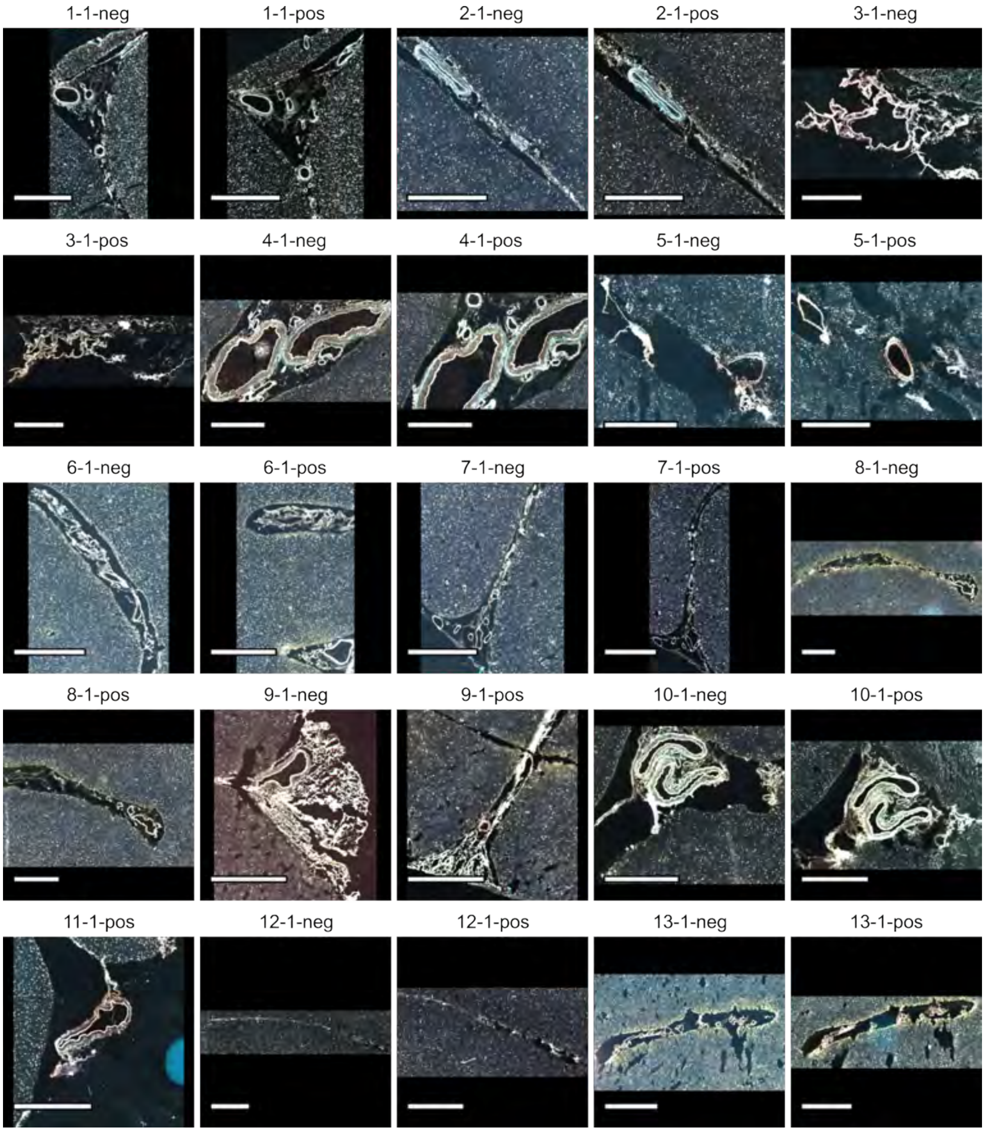


**Figure S1** Autofluorescence images of technical replicate 1. Labels are in order of case number, technical replicate, and MALDI IMS polarity. Case 11 (negative ion mode) was omitted from analysis due to technical difficulties. Per-channel standardization was applied across the entire dataset to ensure that color intensities are directly comparable across Figures S1-S3. Scale bar is equal to 1 mm.

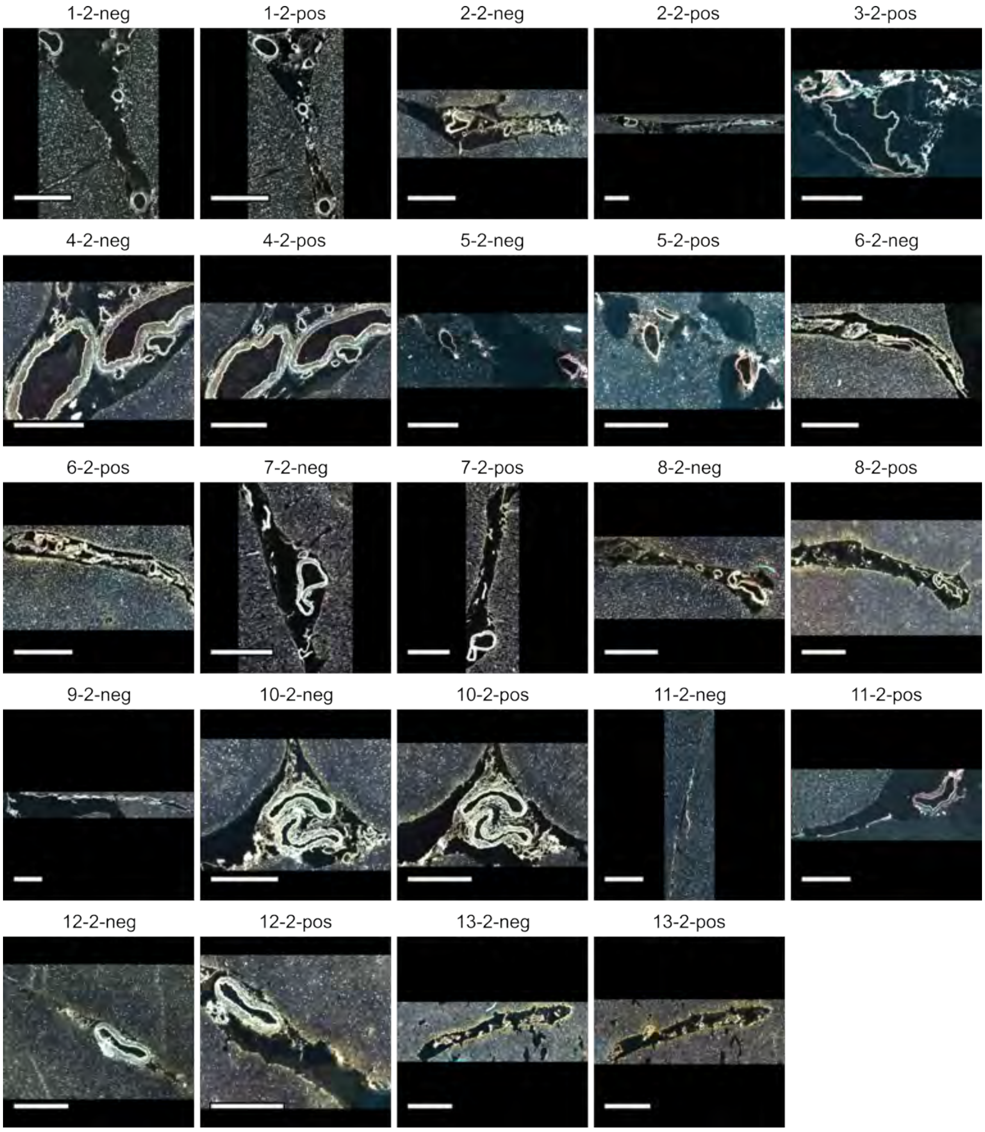


**Figure S2** Autofluorescence images of technical replicate 2. Labels are in order of case number, technical replicate, and MALDI IMS polarity. Case 3 (negative ion mode) and 9 (positive ion mode) were omitted from analyses due to technical difficulties. Per-channel standardization was applied across the entire dataset to ensure that color intensities are directly comparable across Figures S1-S3. Scale bar is equal to 1 mm.

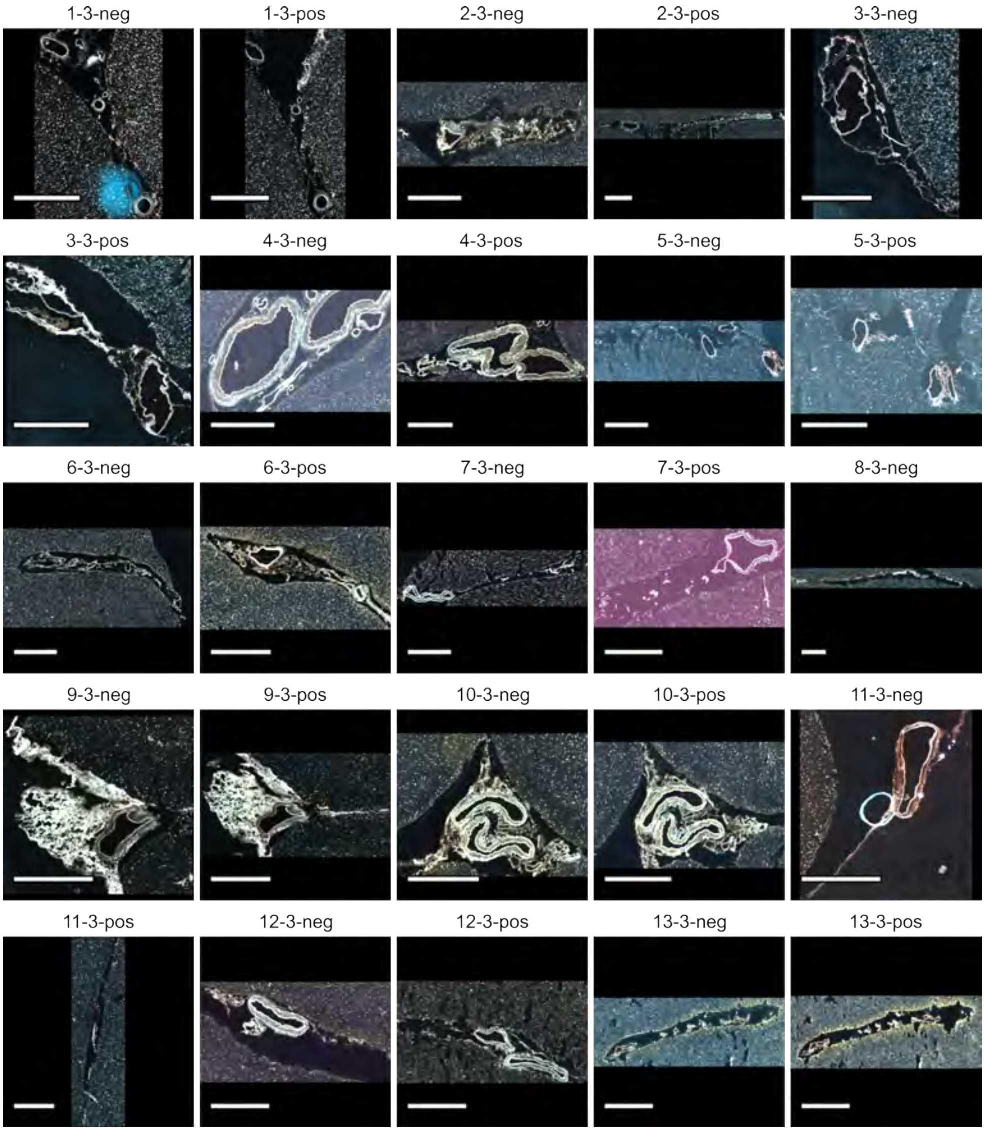


**Figure S3** Autofluorescence images of technical replicate 3. Labels are in order of case number, technical replicate, and MALDI IMS polarity. Case 8 (positive ion mode) was omitted from analyses due to technical difficulties. Per-channel standardization was applied across the entire dataset to ensure that color intensities are directly comparable across Figures S1-S3. Scale bar is equal to 1 mm.

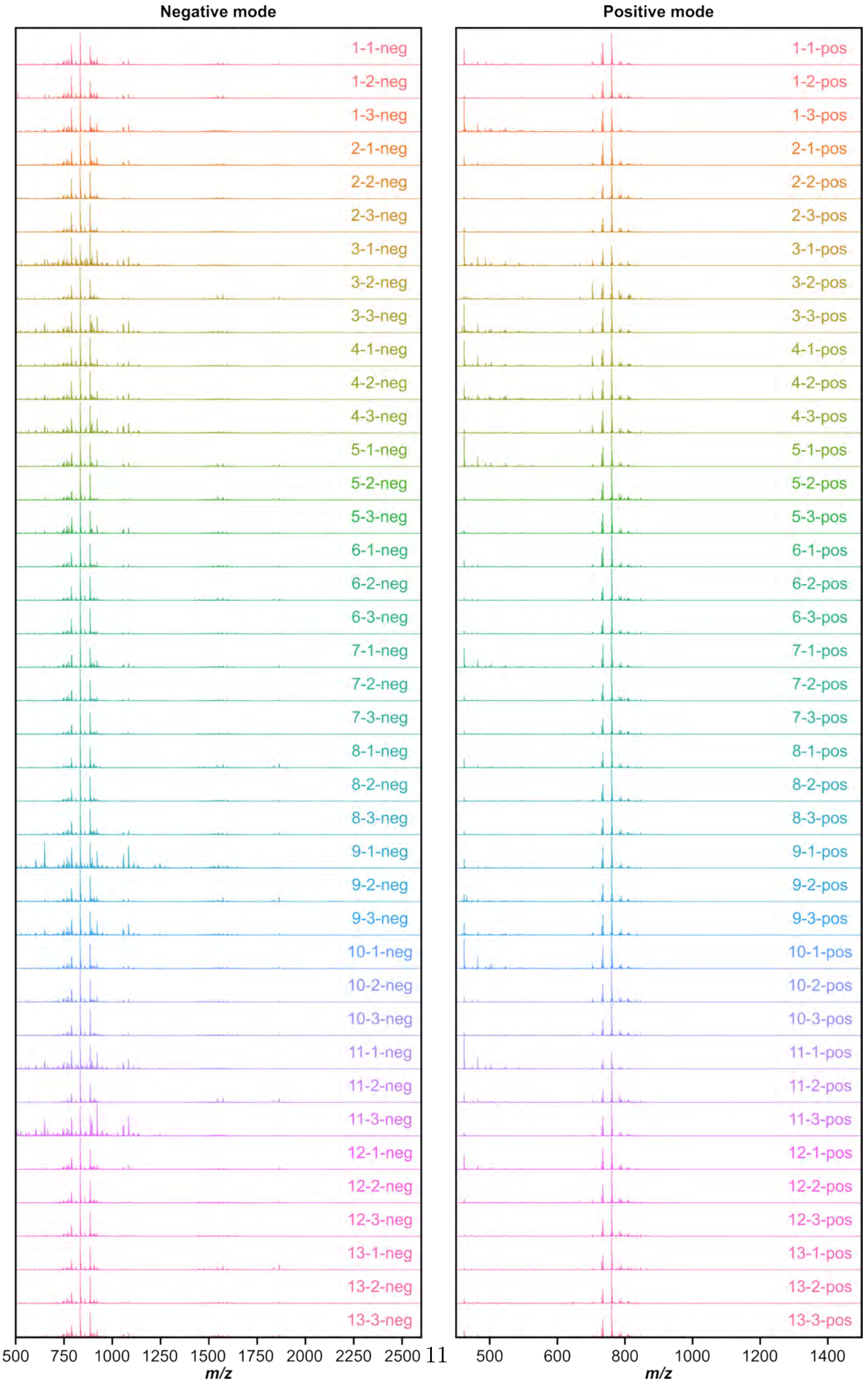


**Figure S4** Average spectra for all cases in negative and positive ion modes. Labels are in order of case number, technical replicate, and polarity.

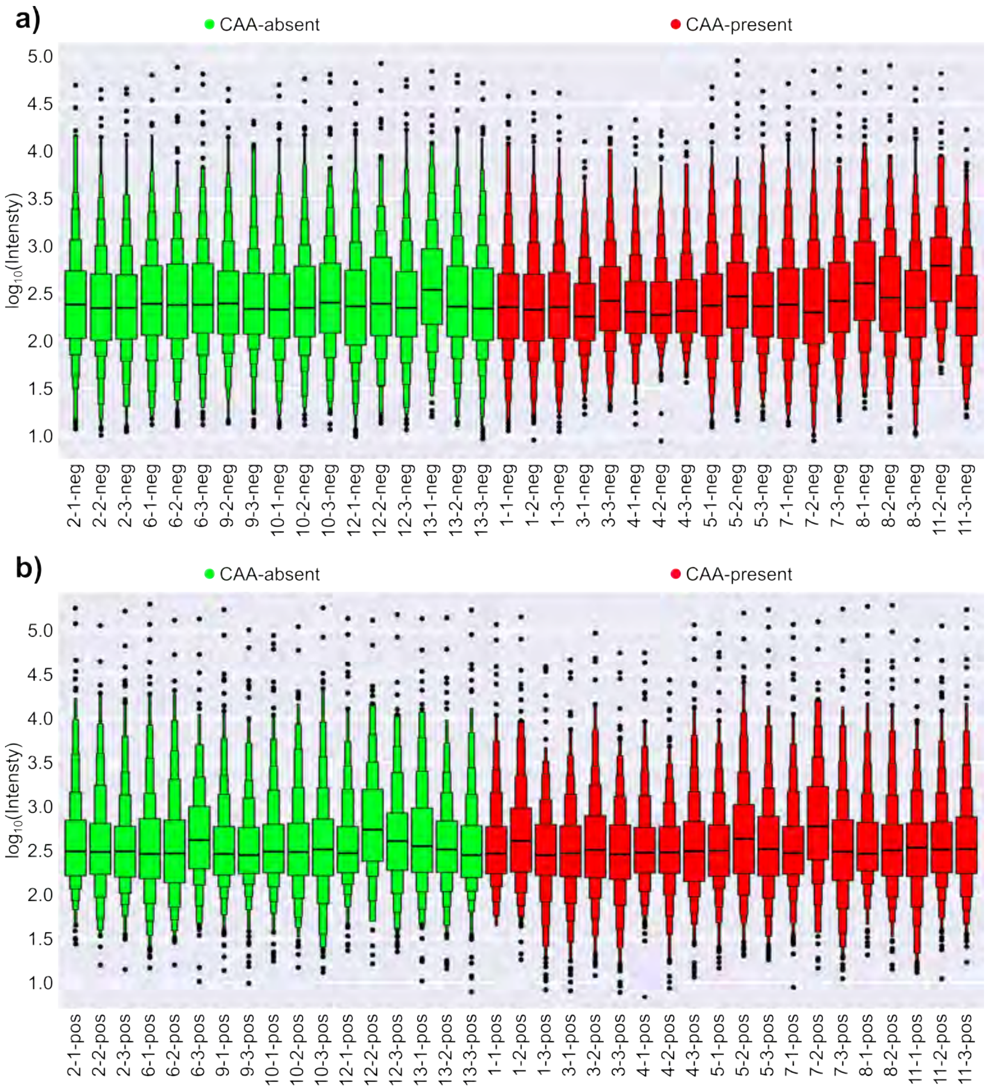


**Figure S5** Levey-Jennings plots highlighting intensity variation across all replicates for negative ion mode (a) and positive ion mode (b). Labels are in order of case number, technical replicate, and polarity.

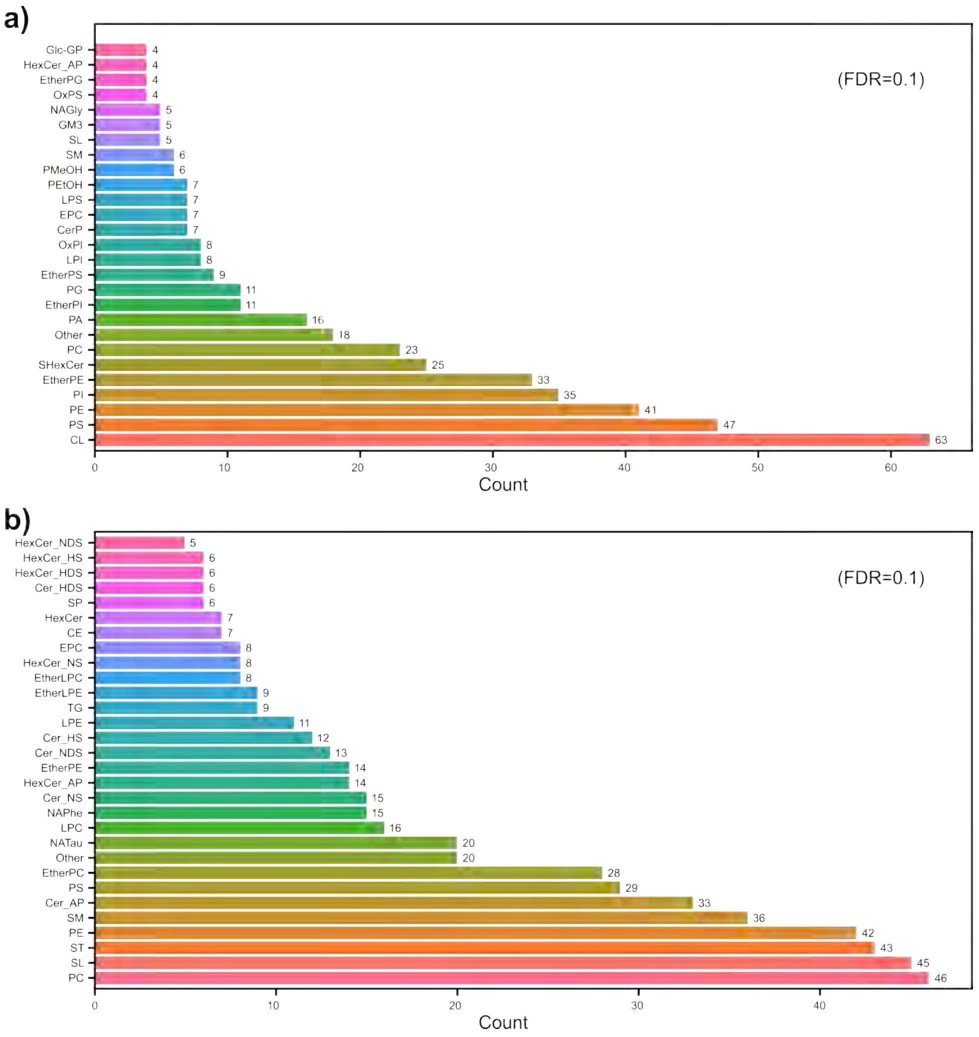


**Figure S6** Total annotated lipids per molecular class for negative ion mode (a) and positive ion mode (b). Tentative identifications were made using an in-house LC-MS/MS library and lipidmaps.org.

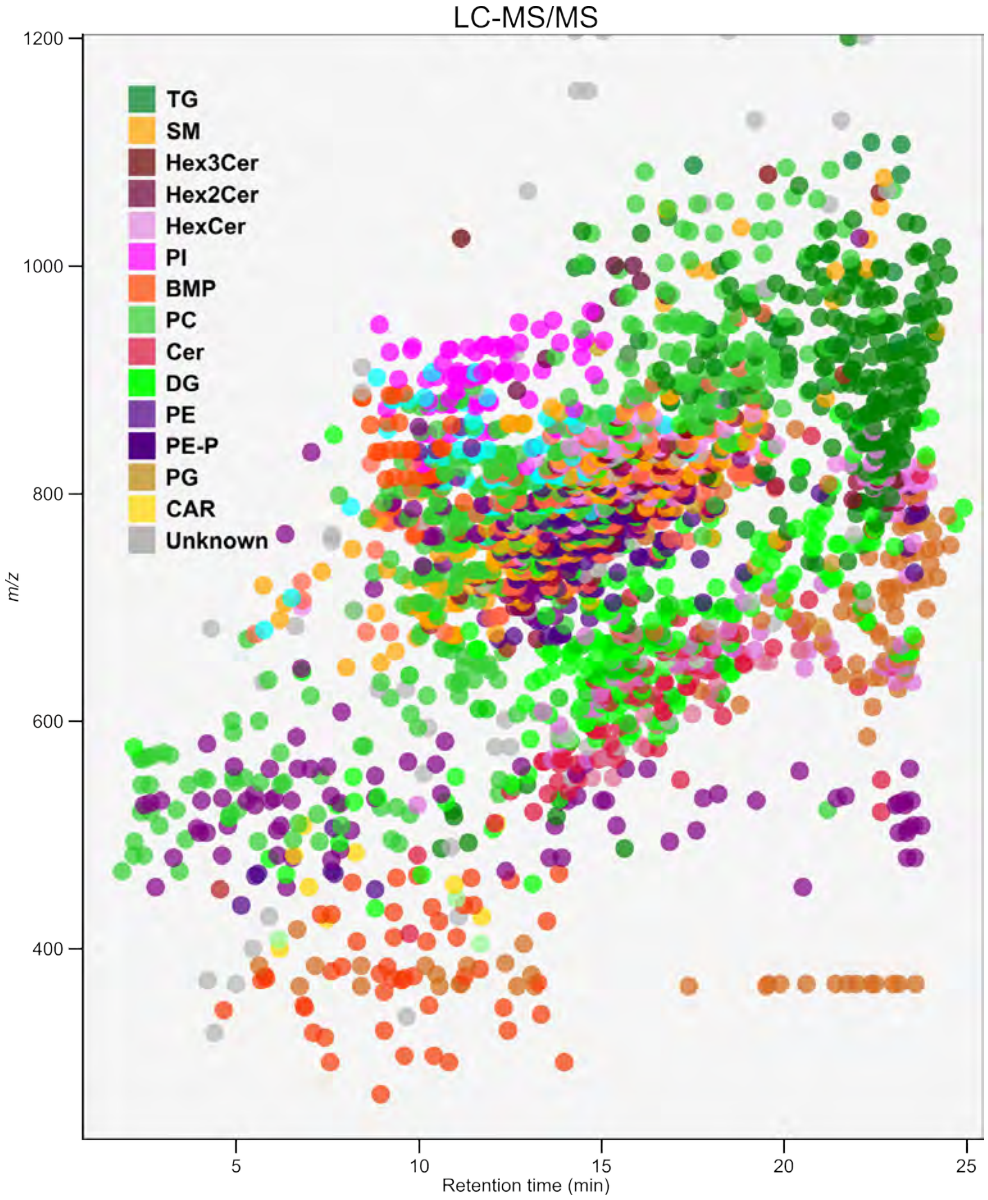


**Figure S7** Positive ion mode LC-MS/MS retention time vs. *m/z* plot.

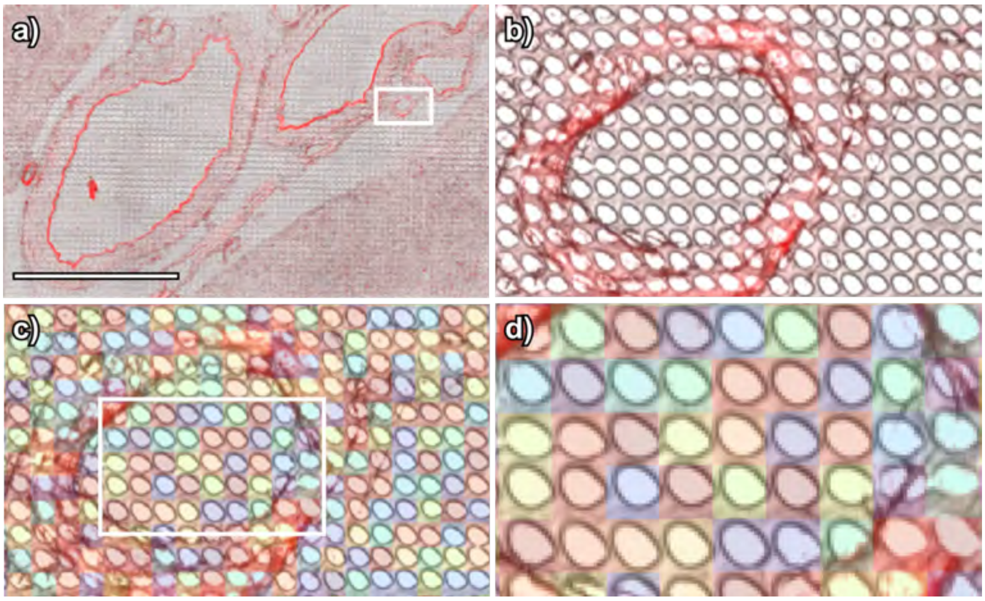


**Figure S8** Visual example of co-registration accuracy for case 4. (a) Negative ion mode, technical replicate 3, brightfield and autofluorescence image of MALDI IMS region. (b) Close-up view of the white box region of a). (c) Overlay of MALDI IMS onto the magnified brightfield image. Each IMS pixel is randomly colored to clearly visualize each pixel individually. (d) Close-up view of the white box region of c), with MALDI IMS overlay showing accuracy of $<$1 µm error. Each colored pixel is 10x10 µm and the scale bar in (a) is 1 mm.

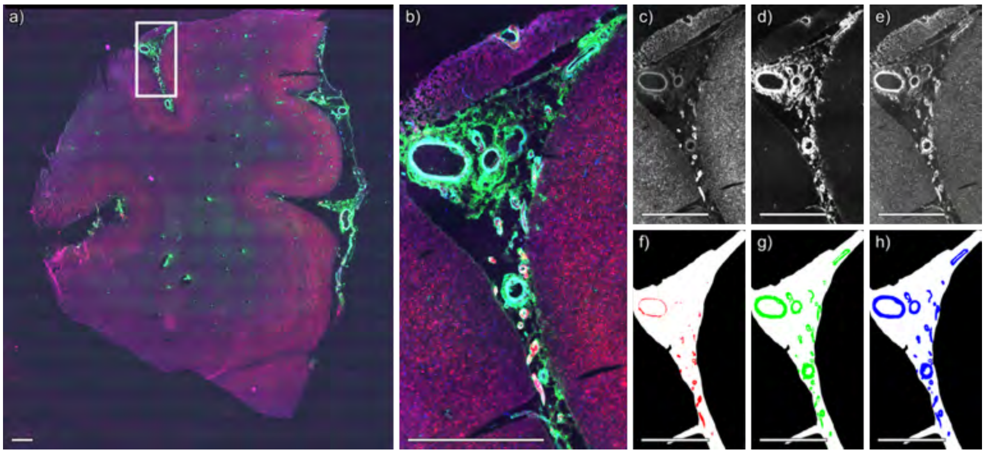


**Figure S9** Segmentation of immunofluorescence microscopy image of case 1, technical replicate 1, negative ion mode. (a) Whole slide immunofluorescence microscopy image. (b) Immunofluorescence image of region acquired by MALDI IMS, shown in a) as a white box. (c) Thiazine red stains amyloid, (d) anti-actin antibodies label smooth muscle, and (e) anti-collagen IV antibodies label vascular basement membranes. Segmentations for (f) amyloid, (g) smooth muscle cells, and (h) collagen IV. Per-channel standardization was applied across the entire dataset to ensure that color intensities are directly comparable across Figures S9-S82. Scale bar is equal to 1 mm.

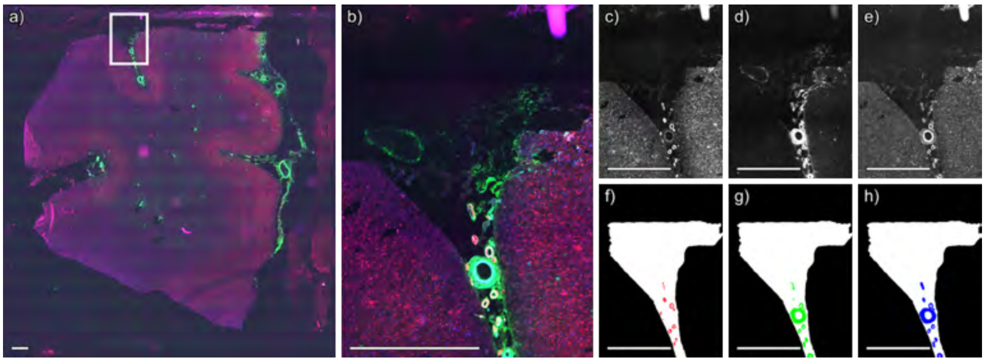


**Figure S10** Segmentation of immunofluorescence microscopy image of case 1, technical replicate 1, positive ion mode. (a) Whole slide immunofluorescence microscopy image. (b) Immunofluorescence image of region acquired by MALDI IMS, shown in a) as a white box. (c) Thiazine red stains amyloid, (d) anti-actin antibodies label smooth muscle, and (e) anti-collagen IV antibodies label vascular basement membranes. Segmentations for (f) amyloid, (g) smooth muscle cells, and (h) collagen IV. Per-channel standardization was applied across the entire dataset to ensure that color intensities are directly comparable across Figures S9-S82. Scale bar is equal to 1 mm.

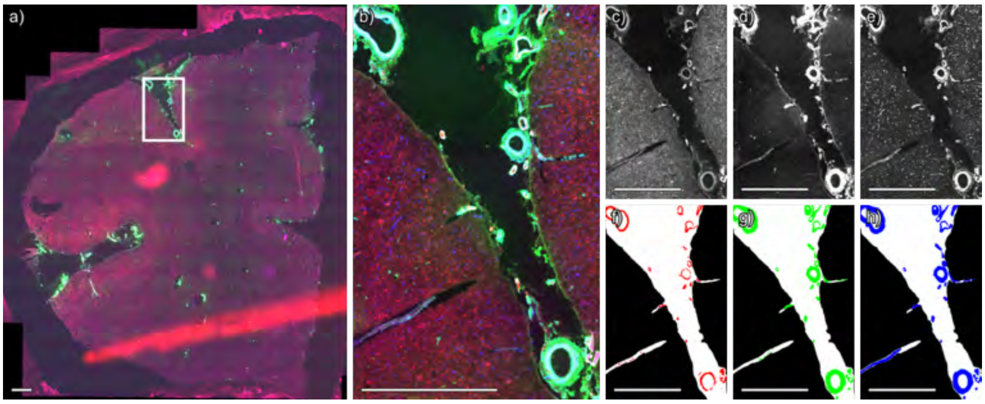


**Figure S11** Segmentation of immunofluorescence microscopy image of case 1, technical replicate 2, negative ion mode. (a) Whole slide immunofluorescence microscopy image. (b) Immunofluorescence image of region acquired by MALDI IMS, shown in a) as a white box. (c) Thiazine red stains amyloid, (d) anti-actin antibodies label smooth muscle, and (e) anti-collagen IV antibodies label vascular basement membranes. Segmentations for (f) amyloid, (g) smooth muscle cells, and (h) collagen IV. Per-channel standardization was applied across the entire dataset to ensure that color intensities are directly comparable across Figures S9-S82. Scale bar is equal to 1 mm.

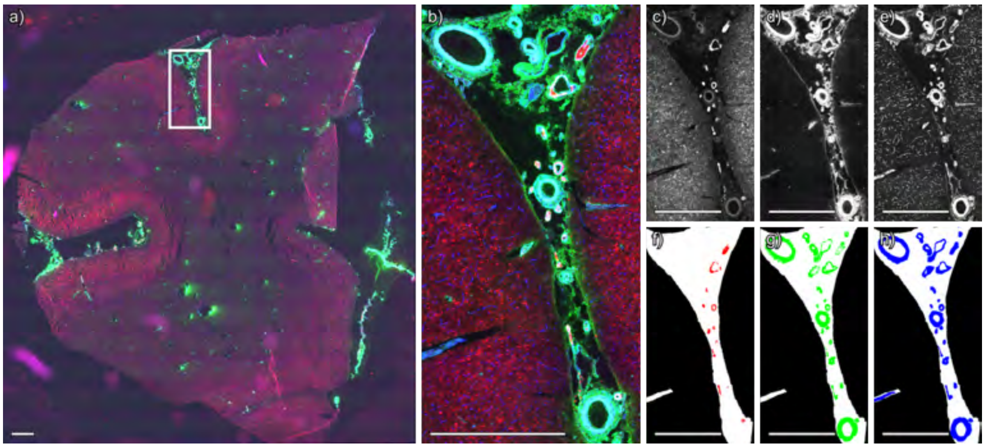


**Figure S12** Segmentation of immunofluorescence microscopy image of case 1, technical replicate 2, positive ion mode. (a) Whole slide immunofluorescence microscopy image. (b) Immunofluorescence image of region acquired by MALDI IMS, shown in a) as a white box. (c) Thiazine red stains amyloid, (d) anti-actin antibodies label smooth muscle, and (e) anti-collagen IV antibodies label vascular basement membranes. Segmentations for (f) amyloid, (g) smooth muscle cells, and (h) collagen IV. Per-channel standardization was applied across the entire dataset to ensure that color intensities are directly comparable across Figures S9-S82. Scale bar is equal to 1 mm.

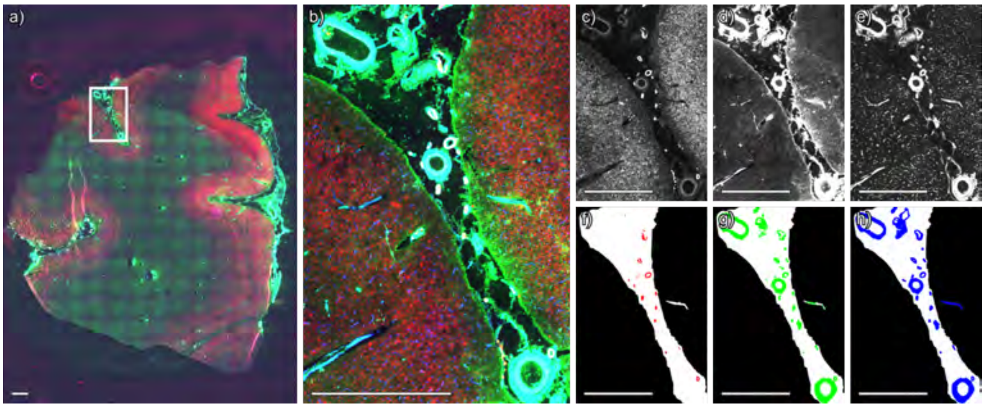


**Figure S13** Segmentation of immunofluorescence microscopy image of case 1, technical replicate 3, negative ion mode. (a) Whole slide immunofluorescence microscopy image. (b) Immunofluorescence image of region acquired by MALDI IMS, shown in a) as a white box. (c) Thiazine red stains amyloid, (d) anti-actin antibodies label smooth muscle, and (e) anti-collagen IV antibodies label vascular basement membranes. Segmentations for (f) amyloid, (g) smooth muscle cells, and (h) collagen IV. Per-channel standardization was applied across the entire dataset to ensure that color intensities are directly comparable across Figures S9-S82. Scale bar is equal to 1 mm.

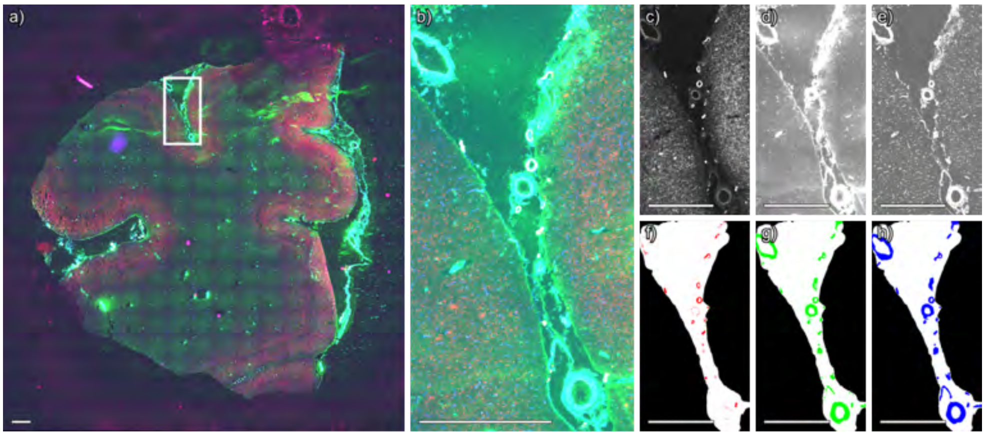


**Figure S14** Segmentation of immunofluorescence microscopy image of case 1, technical replicate 3, positive ion mode. (a) Whole slide immunofluorescence microscopy image. (b) Immunofluorescence image of region acquired by MALDI IMS, shown in a) as a white box. (c) Thiazine red stains amyloid, (d) anti-actin antibodies label smooth muscle, and (e) anti-collagen IV antibodies label vascular basement membranes. Segmentations for (f) amyloid, (g) smooth muscle cells, and (h) collagen IV. Per-channel standardization was applied across the entire dataset to ensure that color intensities are directly comparable across Figures S9-S82. Scale bar is equal to 1 mm.

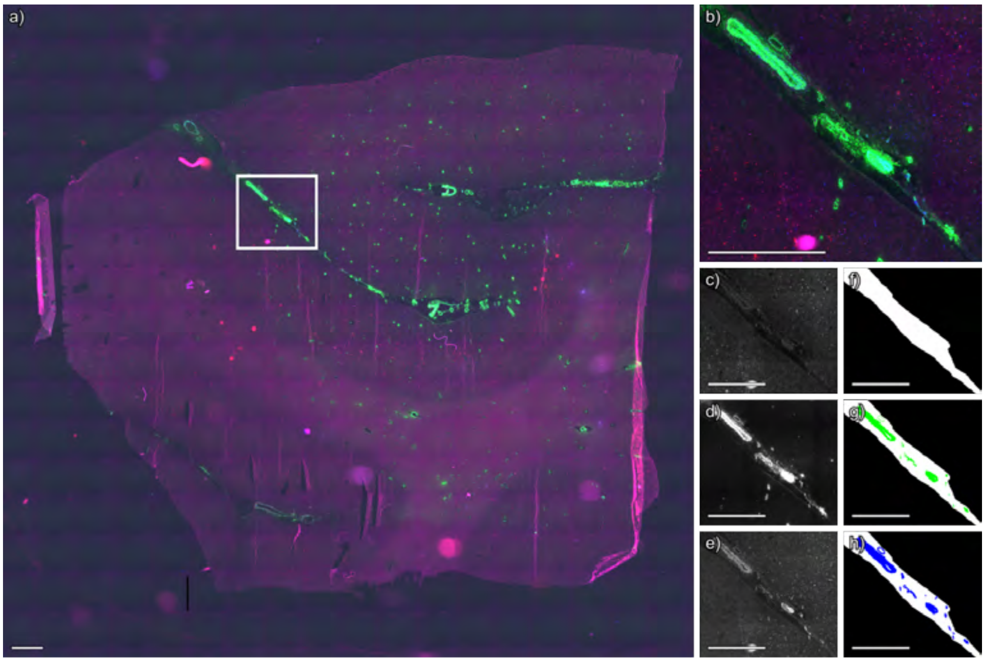


**Figure S15** Segmentation of immunofluorescence microscopy image of case 2, technical replicate 1, negative ion mode. (a) Whole slide immunofluorescence microscopy image. (b) Immunofluorescence image of region acquired by MALDI IMS, shown in a) as a white box. (c) Thiazine red stains amyloid, (d) anti-actin antibodies label smooth muscle, and (e) anti-collagen IV antibodies label vascular basement membranes. Segmentations for (f) amyloid, (g) smooth muscle cells, and (h) collagen IV. Per-channel standardization was applied across the entire dataset to ensure that color intensities are directly comparable across Figures S9-S82. Scale bar is equal to 1 mm.

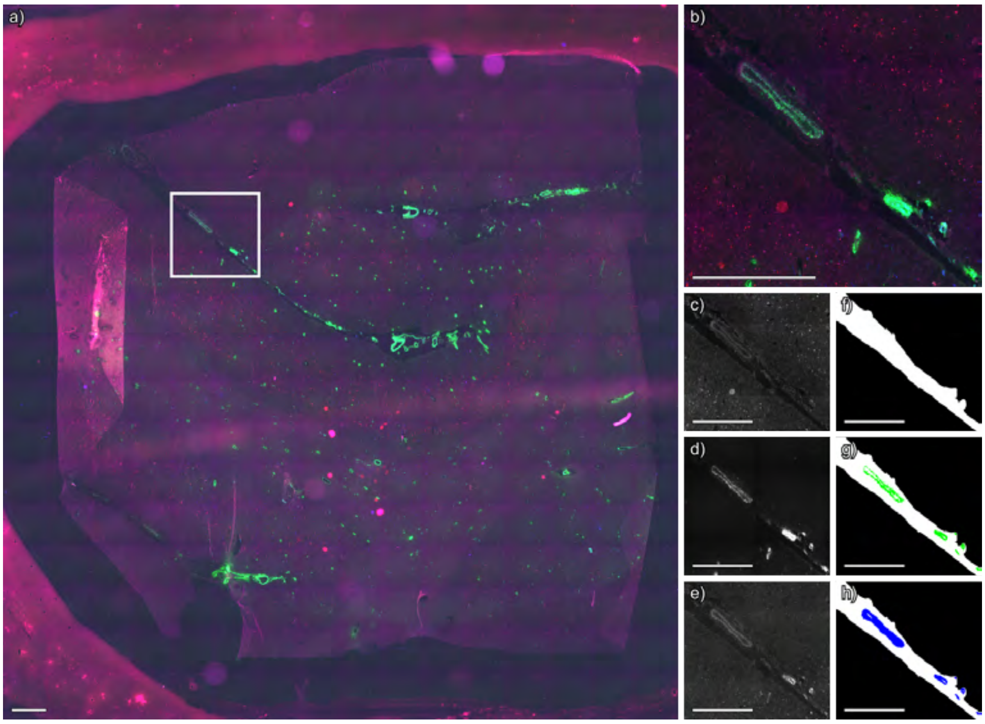


**Figure S16** Segmentation of immunofluorescence microscopy image of case 2, technical replicate 1, positive ion mode. (a) Whole slide immunofluorescence microscopy image. (b) Immunofluorescence image of region acquired by MALDI IMS, shown in a) as a white box. (c) Thiazine red stains amyloid, (d) anti-actin antibodies label smooth muscle, and (e) anti-collagen IV antibodies label vascular basement membranes. Segmentations for (f) amyloid, (g) smooth muscle cells, and (h) collagen IV. Per-channel standardization was applied across the entire dataset to ensure that color intensities are directly comparable across Figures S9-S82. Scale bar is equal to 1 mm.

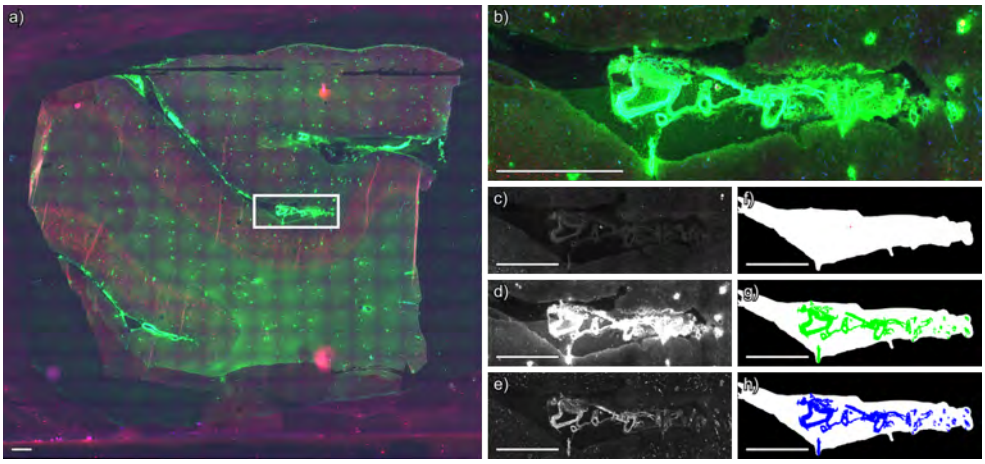


**Figure S17** Segmentation of immunofluorescence microscopy image of case 2, technical replicate 2, negative ion mode. (a) Whole slide immunofluorescence microscopy image. (b) Immunofluorescence image of region acquired by MALDI IMS, shown in a) as a white box. (c) Thiazine red stains amyloid, (d) anti-actin antibodies label smooth muscle, and (e) anti-collagen IV antibodies label vascular basement membranes. Segmentations for (f) amyloid, (g) smooth muscle cells, and (h) collagen IV. Per-channel standardization was applied across the entire dataset to ensure that color intensities are directly comparable across Figures S9-S82. Scale bar is equal to 1 mm.

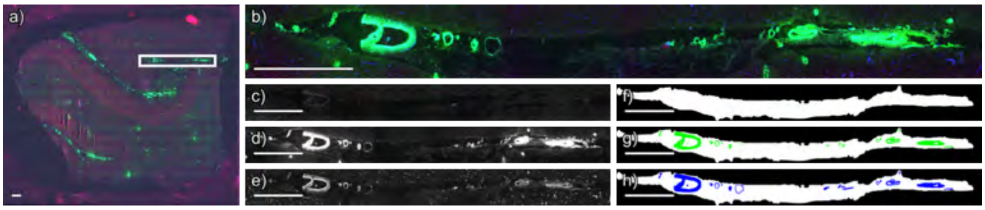


**Figure S18** Segmentation of immunofluorescence microscopy image of case 2, technical replicate 2, positive ion mode. (a) Whole slide immunofluorescence microscopy image. (b) Immunofluorescence image of region acquired by MALDI IMS, shown in a) as a white box. (c) Thiazine red stains amyloid, (d) anti-actin antibodies label smooth muscle, and (e) anti-collagen IV antibodies label vascular basement membranes. Segmentations for (f) amyloid, (g) smooth muscle cells, and (h) collagen IV. Per-channel standardization was applied across the entire dataset to ensure that color intensities are directly comparable across Figures S9-S82. Scale bar is equal to 1 mm.

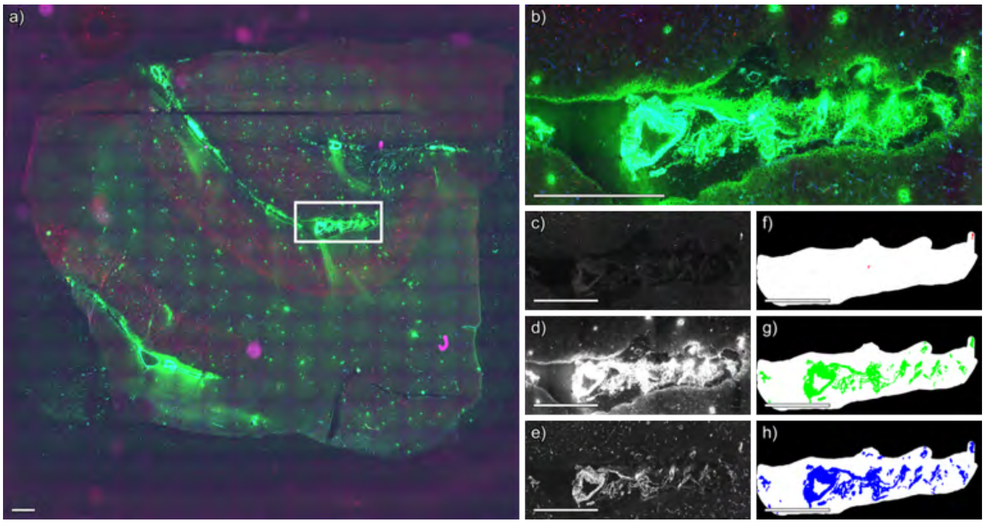


**Figure S19** Segmentation of immunofluorescence microscopy image of case 2, technical replicate 3, negative ion mode. (a) Whole slide immunofluorescence microscopy image. (b) Immunofluorescence image of region acquired by MALDI IMS, shown in a) as a white box. (c) Thiazine red stains amyloid, (d) anti-actin antibodies label smooth muscle, and (e) anti-collagen IV antibodies label vascular basement membranes. Segmentations for (f) amyloid, (g) smooth muscle cells, and (h) collagen IV. Per-channel standardization was applied across the entire dataset to ensure that color intensities are directly comparable across Figures S9-S82. Scale bar is equal to 1 mm.

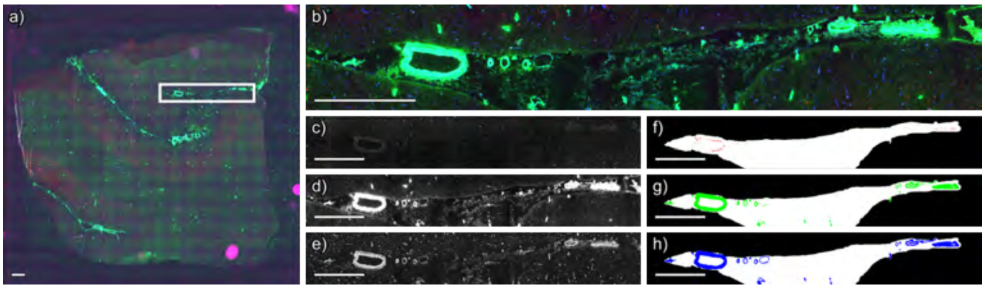


**Figure S20** Segmentation of immunofluorescence microscopy image of case 2, technical replicate 3, positive ion mode. (a) Whole slide immunofluorescence microscopy image. (b) Immunofluorescence image of region acquired by MALDI IMS, shown in a) as a white box. (c) Thiazine red stains amyloid, (d) anti-actin antibodies label smooth muscle, and (e) anti-collagen IV antibodies label vascular basement membranes. Segmentations for (f) amyloid, (g) smooth muscle cells, and (h) collagen IV. Per-channel standardization was applied across the entire dataset to ensure that color intensities are directly comparable across Figures S9-S82. Scale bar is equal to 1 mm.

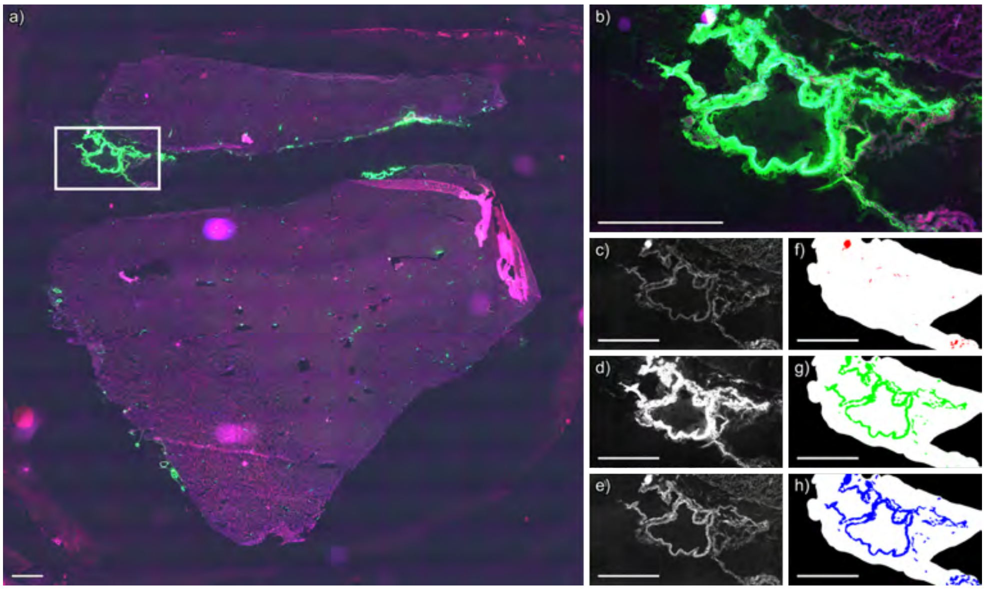


**Figure S21** Segmentation of immunofluorescence microscopy image of case 3, technical replicate 1, negative ion mode. (a) Whole slide immunofluorescence microscopy image. (b) Immunofluorescence image of region acquired by MALDI IMS, shown in a) as a white box. (c) Thiazine red stains amyloid, (d) anti-actin antibodies label smooth muscle, and (e) anti-collagen IV antibodies label vascular basement membranes. Segmentations for (f) amyloid, (g) smooth muscle cells, and (h) collagen IV. Per-channel standardization was applied across the entire dataset to ensure that color intensities are directly comparable across Figures S9-S82. Scale bar is equal to 1 mm.

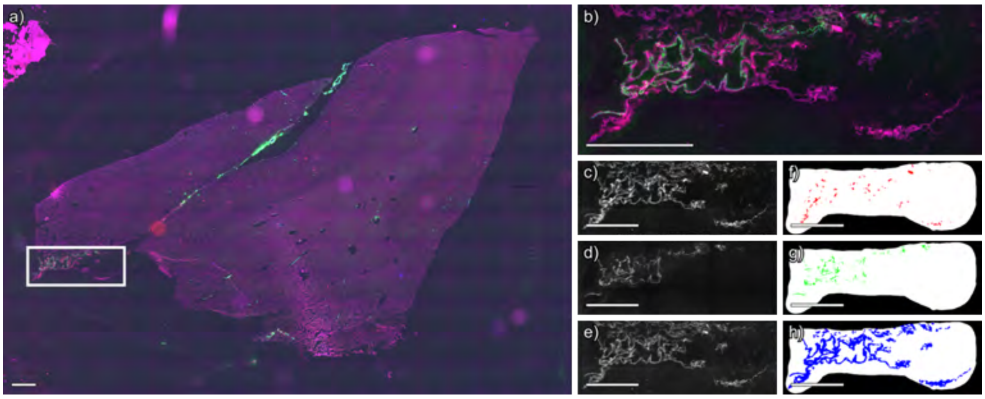


**Figure S22** Segmentation of immunofluorescence microscopy image of case 3, technical replicate 1, positive ion mode. (a) Whole slide immunofluorescence microscopy image. (b) Immunofluorescence image of region acquired by MALDI IMS, shown in a) as a white box. (c) Thiazine red stains amyloid, (d) anti-actin antibodies label smooth muscle, and (e) anti-collagen IV antibodies label vascular basement membranes. Segmentations for (f) amyloid, (g) smooth muscle cells, and (h) collagen IV. Per-channel standardization was applied across the entire dataset to ensure that color intensities are directly comparable across Figures S9-S82. Scale bar is equal to 1 mm.

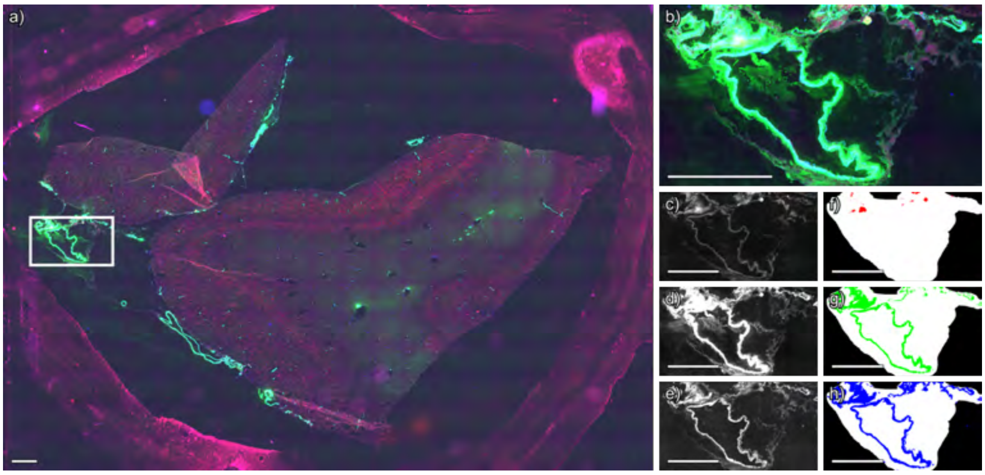


**Figure S23** Segmentation of immunofluorescence microscopy image of case 3, technical replicate 2, positive ion mode. (a) Whole slide immunofluorescence microscopy image. (b) Immunofluorescence image of region acquired by MALDI IMS, shown in a) as a white box. (c) Thiazine red stains amyloid, (d) anti-actin antibodies label smooth muscle, and (e) anti-collagen IV antibodies label vascular basement membranes. Segmentations for (f) amyloid, (g) smooth muscle cells, and (h) collagen IV. Per-channel standardization was applied across the entire dataset to ensure that color intensities are directly comparable across Figures S9-S82. Scale bar is equal to 1 mm.

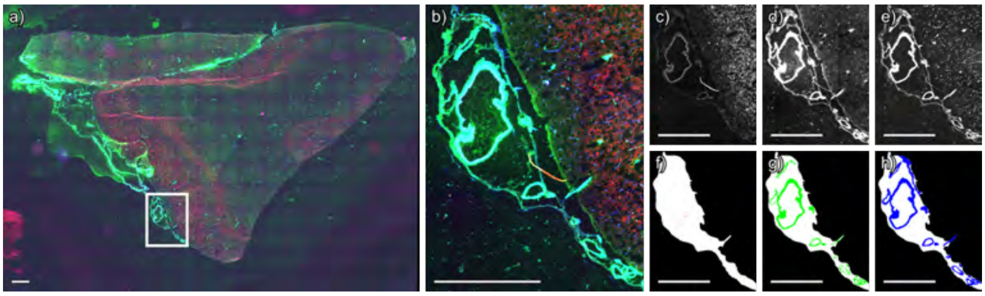


**Figure S24** Segmentation of immunofluorescence microscopy image of case 3, technical replicate 3, negative ion mode. (a) Whole slide immunofluorescence microscopy image. (b) Immunofluorescence image of region acquired by MALDI IMS, shown in a) as a white box. (c) Thiazine red stains amyloid, (d) anti-actin antibodies label smooth muscle, and (e) anti-collagen IV antibodies label vascular basement membranes. Segmentations for (f) amyloid, (g) smooth muscle cells, and (h) collagen IV. Per-channel standardization was applied across the entire dataset to ensure that color intensities are directly comparable across Figures S9-S82. Scale bar is equal to 1 mm.

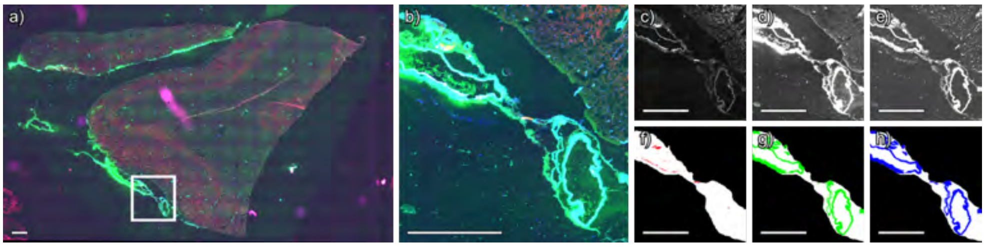


**Figure S25** Segmentation of immunofluorescence microscopy image of case 3, technical replicate 3, positive ion mode. (a) Whole slide immunofluorescence microscopy image. (b) Immunofluorescence image of region acquired by MALDI IMS, shown in a) as a white box. (c) Thiazine red stains amyloid, (d) anti-actin antibodies label smooth muscle, and (e) anti-collagen IV antibodies label vascular basement membranes. Segmentations for (f) amyloid, (g) smooth muscle cells, and (h) collagen IV. Per-channel standardization was applied across the entire dataset to ensure that color intensities are directly comparable across Figures S9-S82. Scale bar is equal to 1 mm.

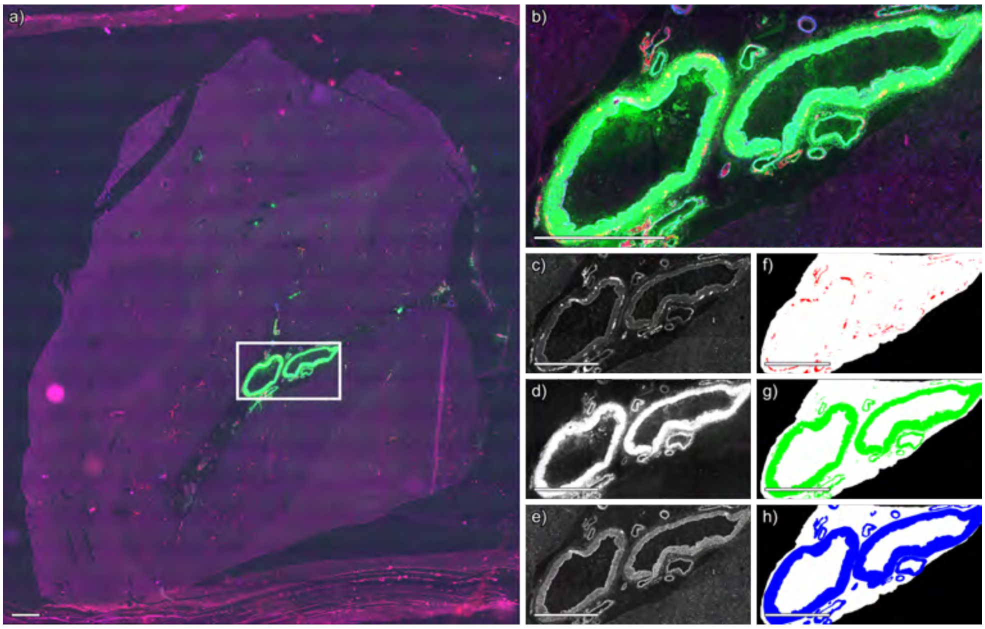


**Figure S26** Segmentation of immunofluorescence microscopy image of case 4, technical replicate 1, negative ion mode. (a) Whole slide immunofluorescence microscopy image. (b) Immunofluorescence image of region acquired by MALDI IMS, shown in a) as a white box. (c) Thiazine red stains amyloid, (d) anti-actin antibodies label smooth muscle, and (e) anti-collagen IV antibodies label vascular basement membranes. Segmentations for (f) amyloid, (g) smooth muscle cells, and (h) collagen IV. Per-channel standardization was applied across the entire dataset to ensure that color intensities are directly comparable across Figures S9-S82. Scale bar is equal to 1 mm.

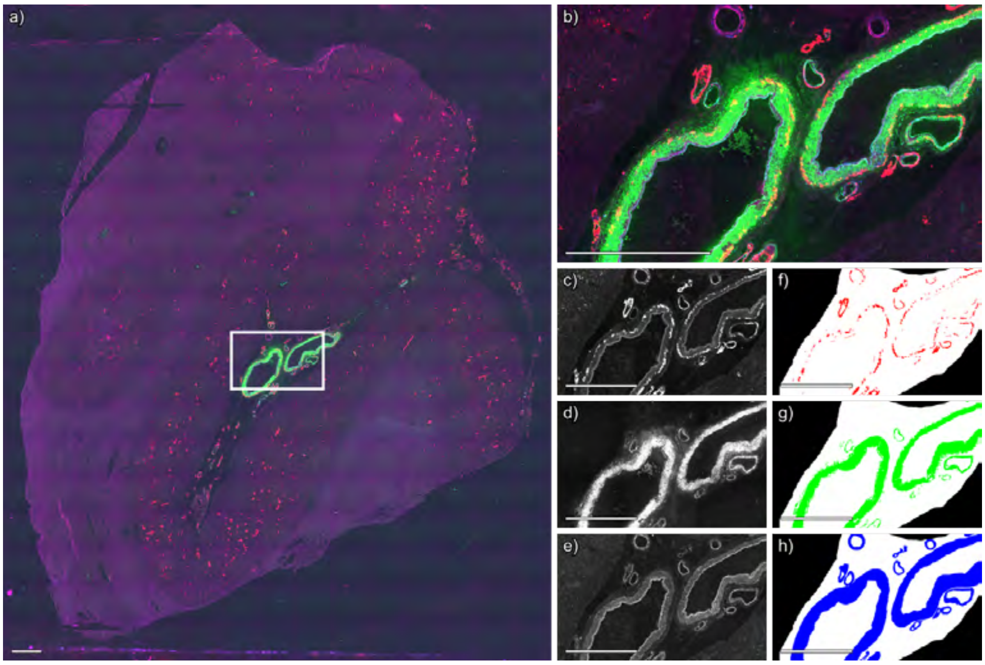


**Figure S27** Segmentation of immunofluorescence microscopy image of case 4, technical replicate 1, positive ion mode. (a) Whole slide immunofluorescence microscopy image. (b) Immunofluorescence image of region acquired by MALDI IMS, shown in a) as a white box. (c) Thiazine red stains amyloid, (d) anti-actin antibodies label smooth muscle, and (e) anti-collagen IV antibodies label vascular basement membranes. Segmentations for (f) amyloid, (g) smooth muscle cells, and (h) collagen IV. Per-channel standardization was applied across the entire dataset to ensure that color intensities are directly comparable across Figures S9-S82. Scale bar is equal to 1 mm.

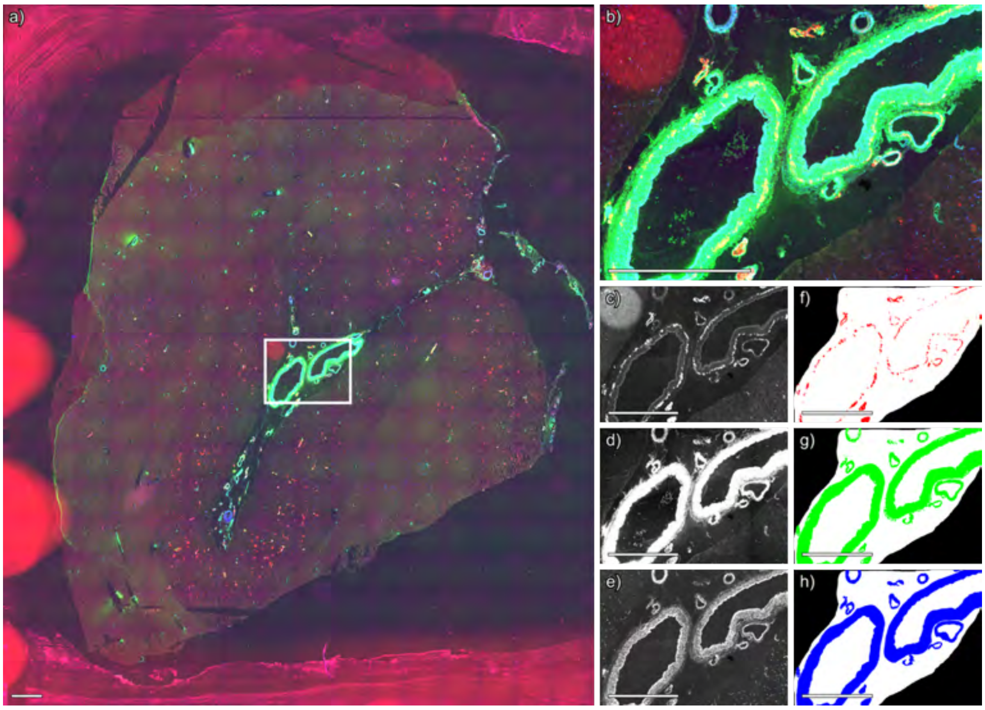


**Figure S28** Segmentation of immunofluorescence microscopy image of case 4, technical replicate 2, negative ion mode. (a) Whole slide immunofluorescence microscopy image. (b) Immunofluorescence image of region acquired by MALDI IMS, shown in a) as a white box. (c) Thiazine red stains amyloid, (d) anti-actin antibodies label smooth muscle, and (e) anti-collagen IV antibodies label vascular basement membranes. Segmentations for (f) amyloid, (g) smooth muscle cells, and (h) collagen IV. Per-channel standardization was applied across the entire dataset to ensure that color intensities are directly comparable across Figures S9-S82. Scale bar is equal to 1 mm.

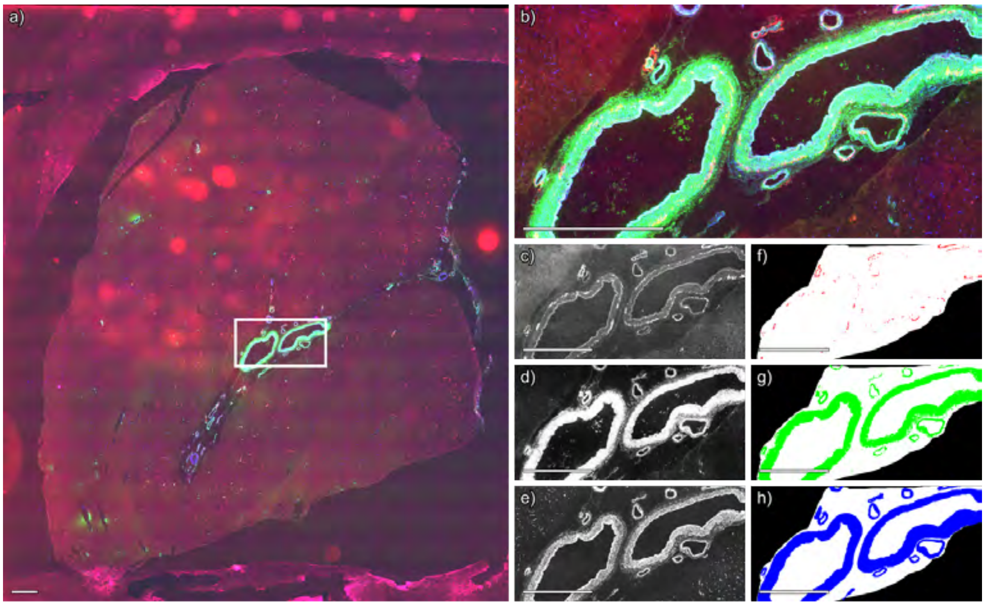


**Figure S29** Segmentation of immunofluorescence microscopy image of case 4, technical replicate 2, positive ion mode. (a) Whole slide immunofluorescence microscopy image. (b) Immunofluorescence image of region acquired by MALDI IMS, shown in a) as a white box. (c) Thiazine red stains amyloid, (d) anti-actin antibodies label smooth muscle, and (e) anti-collagen IV antibodies label vascular basement membranes. Segmentations for (f) amyloid, (g) smooth muscle cells, and (h) collagen IV. Per-channel standardization was applied across the entire dataset to ensure that color intensities are directly comparable across Figures S9-S82. Scale bar is equal to 1 mm.

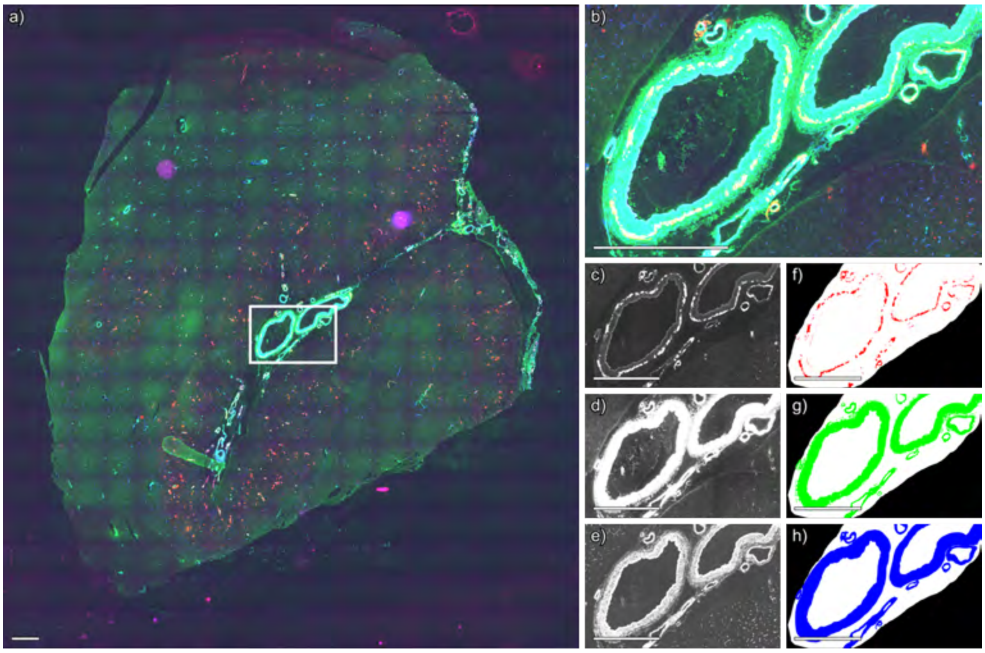


**Figure S30** Segmentation of immunofluorescence microscopy image of case 4, technical replicate 3, negative ion mode. (a) Whole slide immunofluorescence microscopy image. (b) Immunofluorescence image of region acquired by MALDI IMS, shown in a) as a white box. (c) Thiazine red stains amyloid, (d) anti-actin antibodies label smooth muscle, and (e) anti-collagen IV antibodies label vascular basement membranes. Segmentations for (f) amyloid, (g) smooth muscle cells, and (h) collagen IV. Per-channel standardization was applied across the entire dataset to ensure that color intensities are directly comparable across Figures S9-S82. Scale bar is equal to 1 mm.

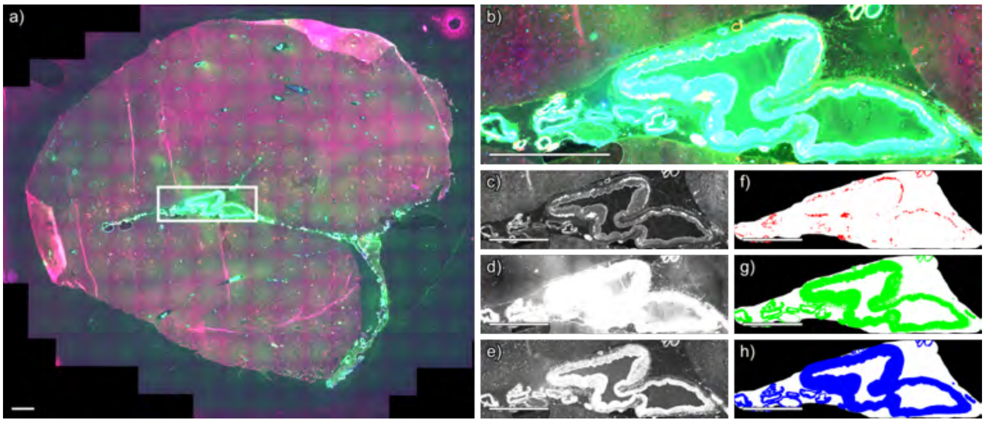


**Figure S31** Segmentation of immunofluorescence microscopy image of case 4, technical replicate 3, positive ion mode. (a) Whole slide immunofluorescence microscopy image. (b) Immunofluorescence image of region acquired by MALDI IMS, shown in a) as a white box. (c) Thiazine red stains amyloid, (d) anti-actin antibodies label smooth muscle, and (e) anti-collagen IV antibodies label vascular basement membranes. Segmentations for (f) amyloid, (g) smooth muscle cells, and (h) collagen IV. Per-channel standardization was applied across the entire dataset to ensure that color intensities are directly comparable across Figures S9-S82. Scale bar is equal to 1 mm.

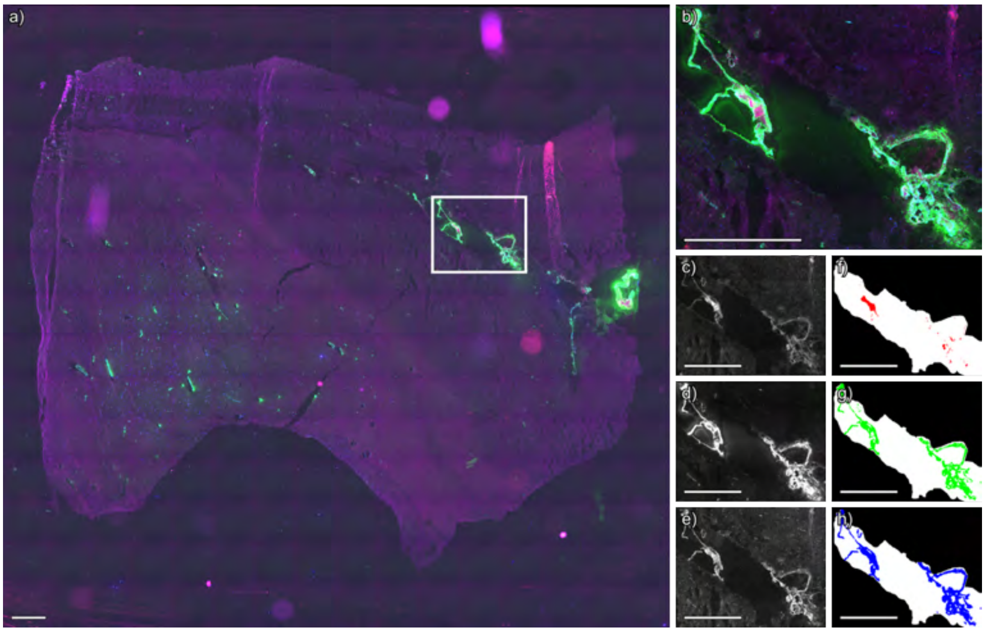


**Figure S32** Segmentation of immunofluorescence microscopy image of case 5, technical replicate 1, negative ion mode. (a) Whole slide immunofluorescence microscopy image. (b) Immunofluorescence image of region acquired by MALDI IMS, shown in a) as a white box. (c) Thiazine red stains amyloid, (d) anti-actin antibodies label smooth muscle, and (e) anti-collagen IV antibodies label vascular basement membranes. Segmentations for (f) amyloid, (g) smooth muscle cells, and (h) collagen IV. Per-channel standardization was applied across the entire dataset to ensure that color intensities are directly comparable across Figures S9-S82. Scale bar is equal to 1 mm.

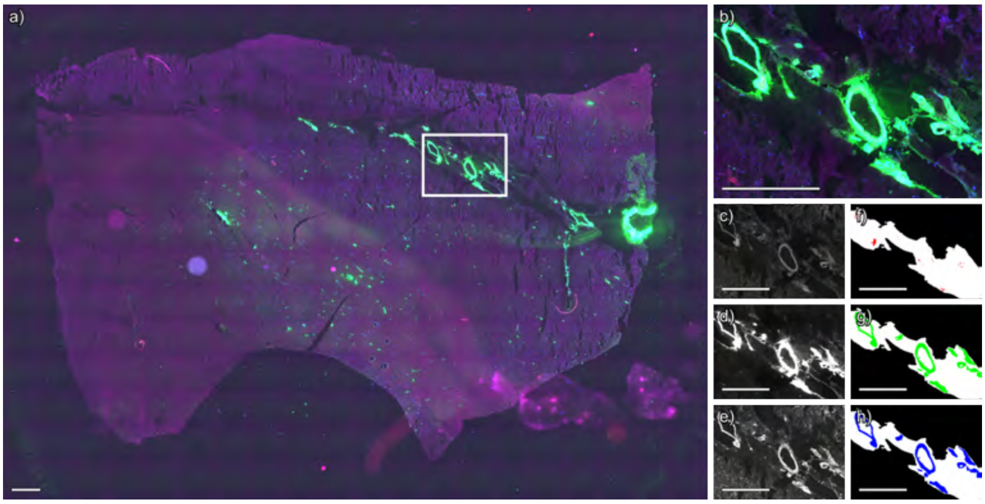


**Figure S33** Segmentation of immunofluorescence microscopy image of case 5, technical replicate 1, positive ion mode. (a) Whole slide immunofluorescence microscopy image. (b) Immunofluorescence image of region acquired by MALDI IMS, shown in a) as a white box. (c) Thiazine red stains amyloid, (d) anti-actin antibodies label smooth muscle, and (e) anti-collagen IV antibodies label vascular basement membranes. Segmentations for (f) amyloid, (g) smooth muscle cells, and (h) collagen IV. Per-channel standardization was applied across the entire dataset to ensure that color intensities are directly comparable across Figures S9-S82. Scale bar is equal to 1 mm.

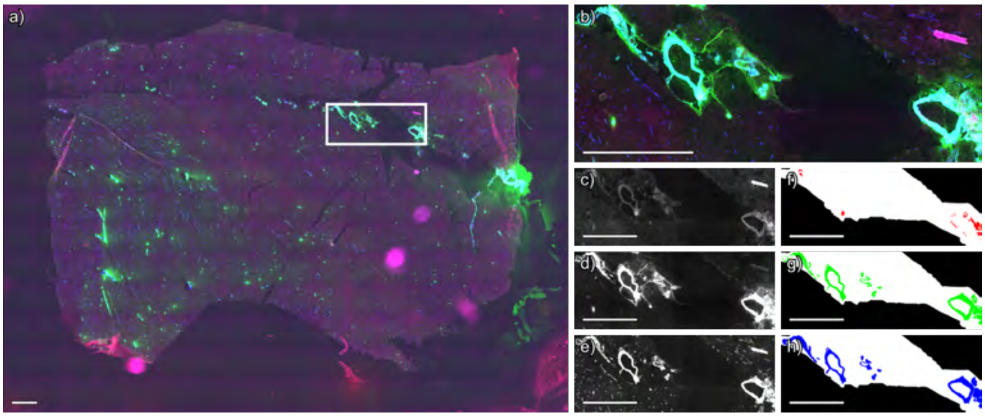


**Figure S34** Segmentation of immunofluorescence microscopy image of case 5, technical replicate 2, negative ion mode. (a) Whole slide immunofluorescence microscopy image. (b) Immunofluorescence image of region acquired by MALDI IMS, shown in a) as a white box. (c) Thiazine red stains amyloid, (d) anti-actin antibodies label smooth muscle, and (e) anti-collagen IV antibodies label vascular basement membranes. Segmentations for (f) amyloid, (g) smooth muscle cells, and (h) collagen IV. Per-channel standardization was applied across the entire dataset to ensure that color intensities are directly comparable across Figures S9-S82. Scale bar is equal to 1 mm.

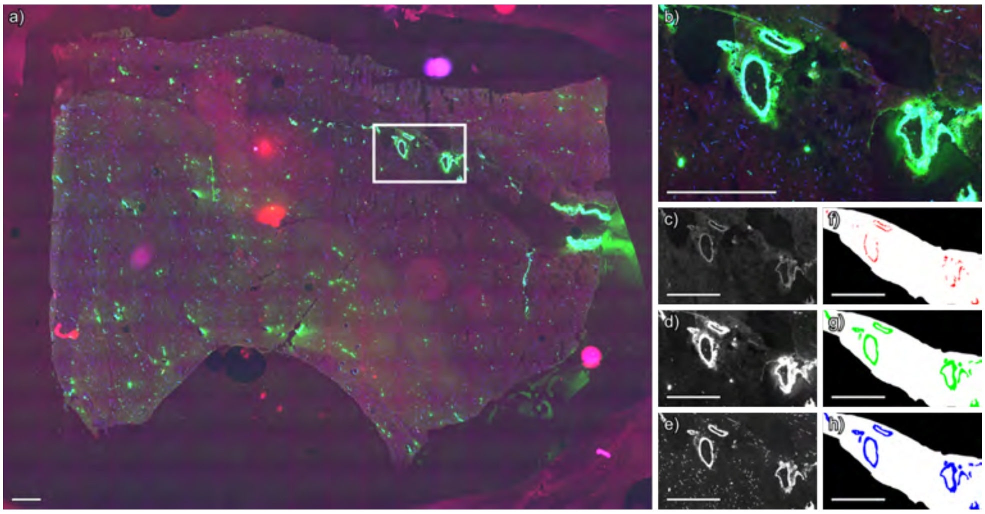


**Figure S35** Segmentation of immunofluorescence microscopy image of case 5, technical replicate 2, positive ion mode. (a) Whole slide immunofluorescence microscopy image. (b) Immunofluorescence image of region acquired by MALDI IMS, shown in a) as a white box. (c) Thiazine red stains amyloid, (d) anti-actin antibodies label smooth muscle, and (e) anti-collagen IV antibodies label vascular basement membranes. Segmentations for (f) amyloid, (g) smooth muscle cells, and (h) collagen IV. Per-channel standardization was applied across the entire dataset to ensure that color intensities are directly comparable across Figures S9-S82. Scale bar is equal to 1 mm.

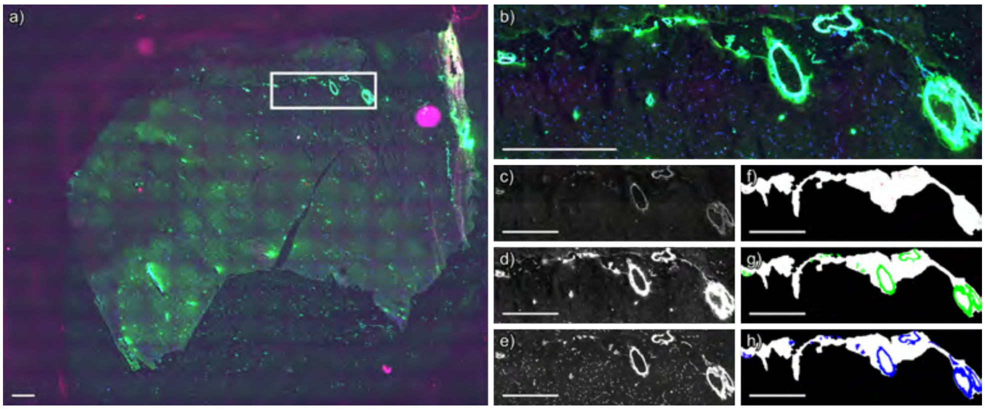


**Figure S36** Segmentation of immunofluorescence microscopy image of case 5, technical replicate 3, negative ion mode. (a) Whole slide immunofluorescence microscopy image. (b) Immunofluorescence image of region acquired by MALDI IMS, shown in a) as a white box. (c) Thiazine red stains amyloid, (d) anti-actin antibodies label smooth muscle, and (e) anti-collagen IV antibodies label vascular basement membranes. Segmentations for (f) amyloid, (g) smooth muscle cells, and (h) collagen IV. Per-channel standardization was applied across the entire dataset to ensure that color intensities are directly comparable across Figures S9-S82. Scale bar is equal to 1 mm.

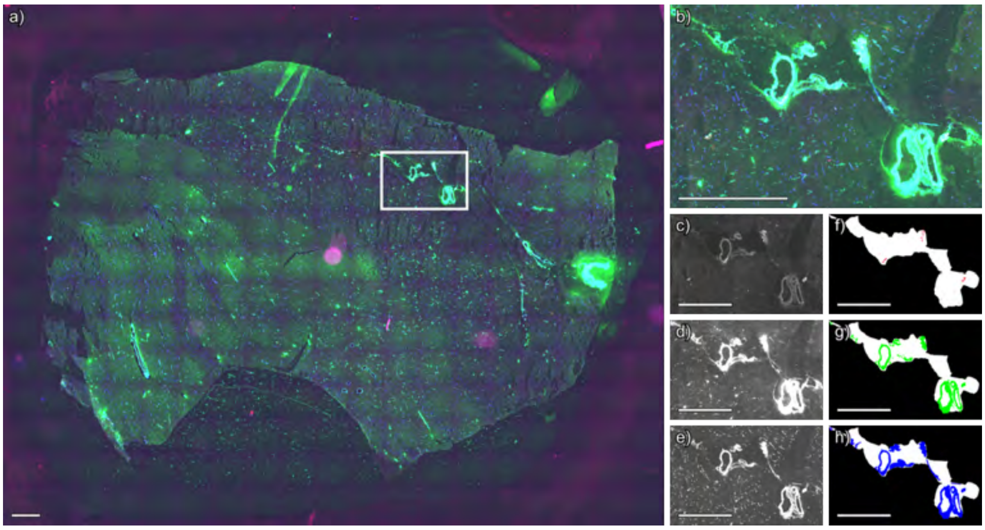


**Figure S37** Segmentation of immunofluorescence microscopy image of case 5, technical replicate 3, positive ion mode. (a) Whole slide immunofluorescence microscopy image. (b) Immunofluorescence image of region acquired by MALDI IMS, shown in a) as a white box. (c) Thiazine red stains amyloid, (d) anti-actin antibodies label smooth muscle, and (e) anti-collagen IV antibodies label vascular basement membranes. Segmentations for (f) amyloid, (g) smooth muscle cells, and (h) collagen IV. Per-channel standardization was applied across the entire dataset to ensure that color intensities are directly comparable across Figures S9-S82. Scale bar is equal to 1 mm.

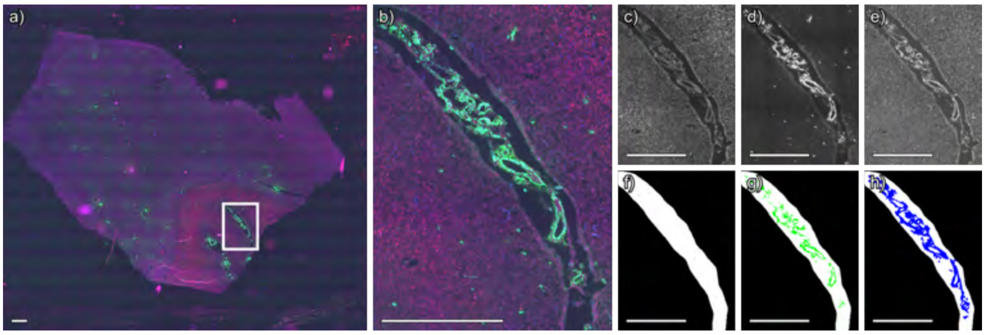


**Figure S38** Segmentation of immunofluorescence microscopy image of case 6, technical replicate 1, negative ion mode. (a) Whole slide immunofluorescence microscopy image. (b) Immunofluorescence image of region acquired by MALDI IMS, shown in a) as a white box. (c) Thiazine red stains amyloid, (d) anti-actin antibodies label smooth muscle, and (e) anti-collagen IV antibodies label vascular basement membranes. Segmentations for (f) amyloid, (g) smooth muscle cells, and (h) collagen IV. Per-channel standardization was applied across the entire dataset to ensure that color intensities are directly comparable across Figures S9-S82. Scale bar is equal to 1 mm.

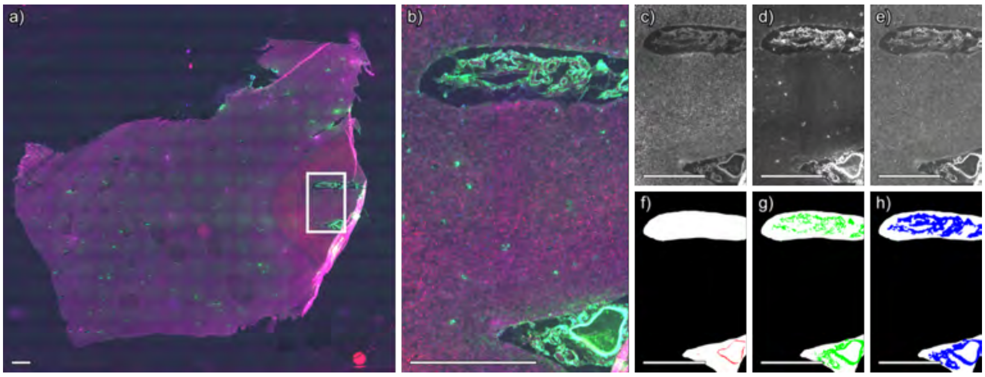


**Figure S39** Segmentation of immunofluorescence microscopy image of case 6, technical replicate 1, positive ion mode. (a) Whole slide immunofluorescence microscopy image. (b) Immunofluorescence image of region acquired by MALDI IMS, shown in a) as a white box. (c) Thiazine red stains amyloid, (d) anti-actin antibodies label smooth muscle, and (e) anti-collagen IV antibodies label vascular basement membranes. Segmentations for (f) amyloid, (g) smooth muscle cells, and (h) collagen IV. Per-channel standardization was applied across the entire dataset to ensure that color intensities are directly comparable across Figures S9-S82. Scale bar is equal to 1 mm.

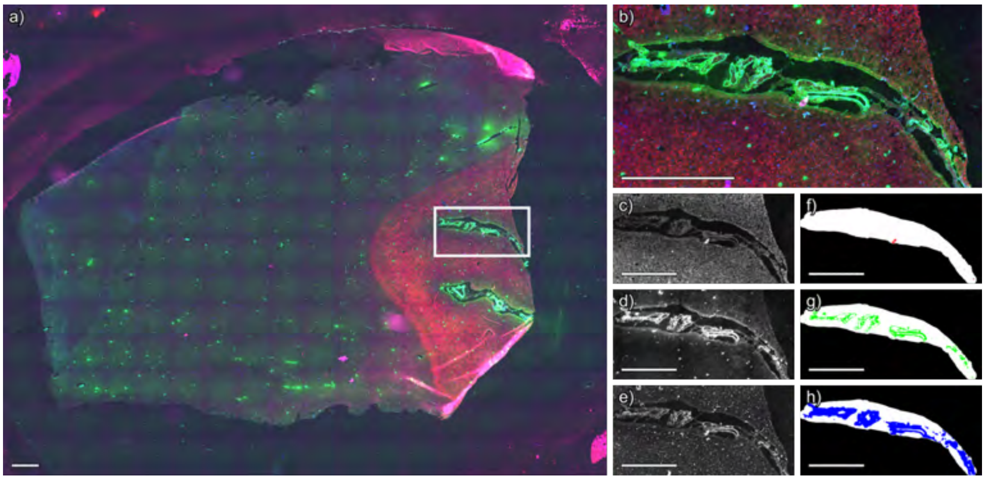


**Figure S40** Segmentation of immunofluorescence microscopy image of case 6, technical replicate 2, negative ion mode. (a) Whole slide immunofluorescence microscopy image. (b) Immunofluorescence image of region acquired by MALDI IMS, shown in a) as a white box. (c) Thiazine red stains amyloid, (d) anti-actin antibodies label smooth muscle, and (e) anti-collagen IV antibodies label vascular basement membranes. Segmentations for (f) amyloid, (g) smooth muscle cells, and (h) collagen IV. Per-channel standardization was applied across the entire dataset to ensure that color intensities are directly comparable across Figures S9-S82. Scale bar is equal to 1 mm.

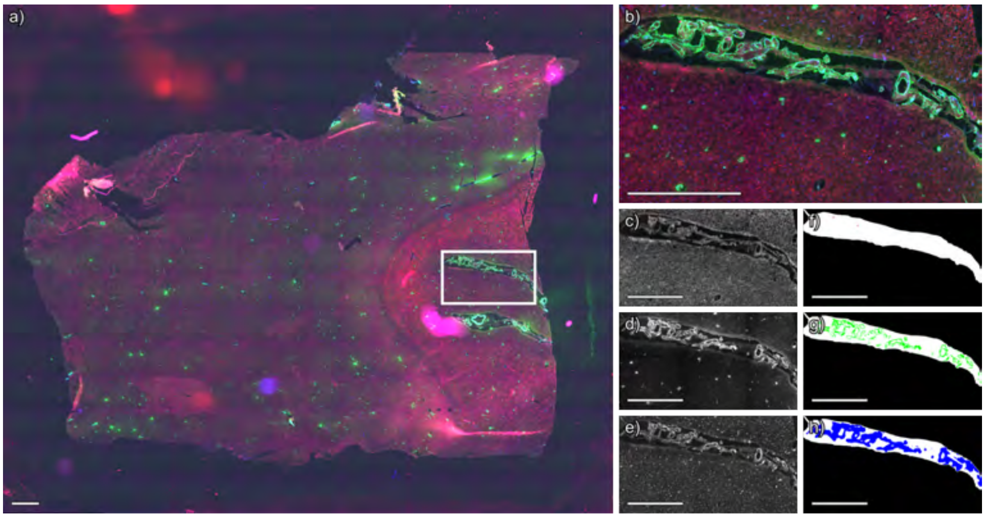


**Figure S41** Segmentation of immunofluorescence microscopy image of case 6, technical replicate 2, positive ion mode. (a) Whole slide immunofluorescence microscopy image. (b) Immunofluorescence image of region acquired by MALDI IMS, shown in a) as a white box. (c) Thiazine red stains amyloid, (d) anti-actin antibodies label smooth muscle, and (e) anti-collagen IV antibodies label vascular basement membranes. Segmentations for (f) amyloid, (g) smooth muscle cells, and (h) collagen IV. Per-channel standardization was applied across the entire dataset to ensure that color intensities are directly comparable across Figures S9-S82. Scale bar is equal to 1 mm.

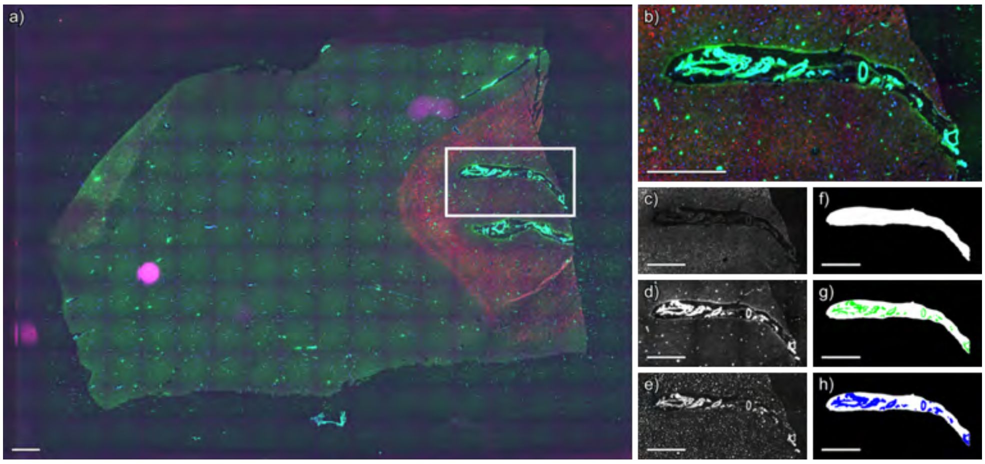


**Figure S42** Segmentation of immunofluorescence microscopy image of case 6, technical replicate 3, negative ion mode. (a) Whole slide immunofluorescence microscopy image. (b) Immunofluorescence image of region acquired by MALDI IMS, shown in a) as a white box. (c) Thiazine red stains amyloid, (d) anti-actin antibodies label smooth muscle, and (e) anti-collagen IV antibodies label vascular basement membranes. Segmentations for (f) amyloid, (g) smooth muscle cells, and (h) collagen IV. Per-channel standardization was applied across the entire dataset to ensure that color intensities are directly comparable across Figures S9-S82. Scale bar is equal to 1 mm.

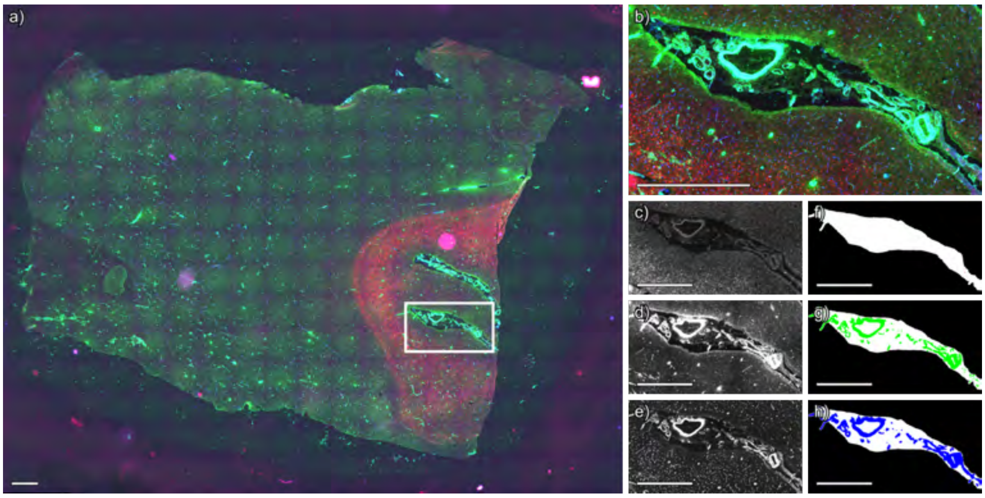


**Figure S43** Segmentation of immunofluorescence microscopy image of case 6, technical replicate 3, positive ion mode. (a) Whole slide immunofluorescence microscopy image. (b) Immunofluorescence image of region acquired by MALDI IMS, shown in a) as a white box. (c) Thiazine red stains amyloid, (d) anti-actin antibodies label smooth muscle, and (e) anti-collagen IV antibodies label vascular basement membranes. Segmentations for (f) amyloid, (g) smooth muscle cells, and (h) collagen IV. Per-channel standardization was applied across the entire dataset to ensure that color intensities are directly comparable across Figures S9-S82. Scale bar is equal to 1 mm.

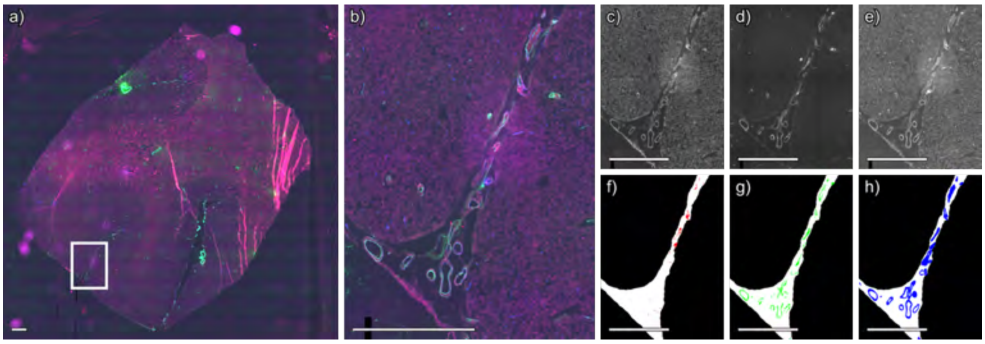


**Figure S44** Segmentation of immunofluorescence microscopy image of case 7, technical replicate 1, negative ion mode. (a) Whole slide immunofluorescence microscopy image. (b) Immunofluorescence image of region acquired by MALDI IMS, shown in a) as a white box. (c) Thiazine red stains amyloid, (d) anti-actin antibodies label smooth muscle, and (e) anti-collagen IV antibodies label vascular basement membranes. Segmentations for (f) amyloid, (g) smooth muscle cells, and (h) collagen IV. Per-channel standardization was applied across the entire dataset to ensure that color intensities are directly comparable across Figures S9-S82. Scale bar is equal to 1 mm.

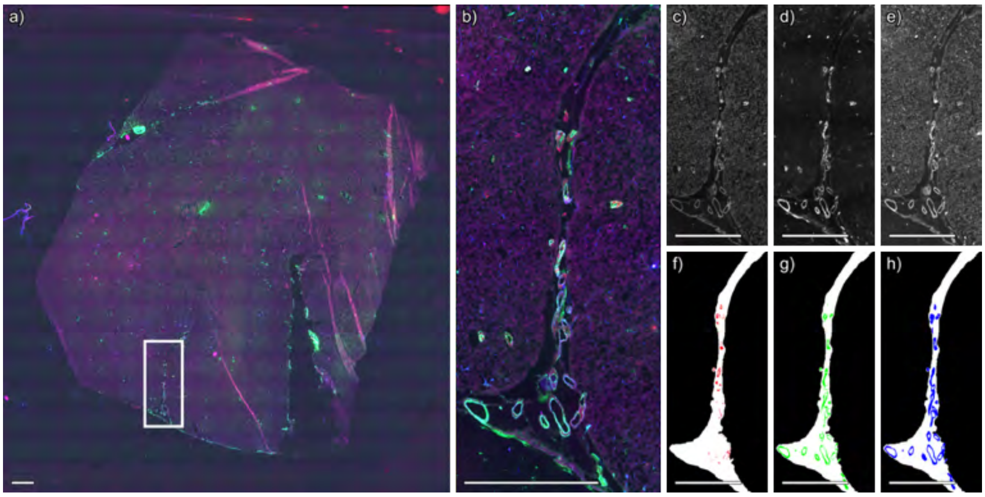


**Figure S45** Segmentation of immunofluorescence microscopy image of case 7, technical replicate 1, positive ion mode. (a) Whole slide immunofluorescence microscopy image. (b) Immunofluorescence image of region acquired by MALDI IMS, shown in a) as a white box. (c) Thiazine red stains amyloid, (d) anti-actin antibodies label smooth muscle, and (e) anti-collagen IV antibodies label vascular basement membranes. Segmentations for (f) amyloid, (g) smooth muscle cells, and (h) collagen IV. Per-channel standardization was applied across the entire dataset to ensure that color intensities are directly comparable across Figures S9-S82. Scale bar is equal to 1 mm.

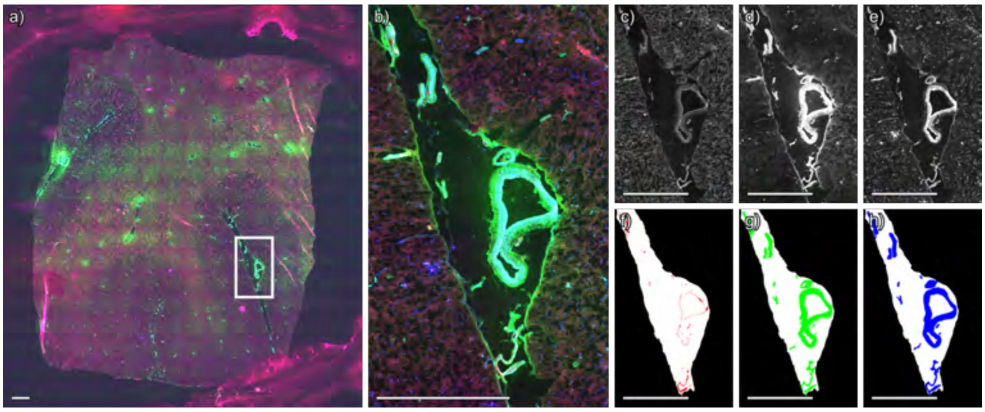


**Figure S46** Segmentation of immunofluorescence microscopy image of case 7, technical replicate 2, negative ion mode. (a) Whole slide immunofluorescence microscopy image. (b) Immunofluorescence image of region acquired by MALDI IMS, shown in a) as a white box. (c) Thiazine red stains amyloid, (d) anti-actin antibodies label smooth muscle, and (e) anti-collagen IV antibodies label vascular basement membranes. Segmentations for (f) amyloid, (g) smooth muscle cells, and (h) collagen IV. Per-channel standardization was applied across the entire dataset to ensure that color intensities are directly comparable across Figures S9-S82. Scale bar is equal to 1 mm.

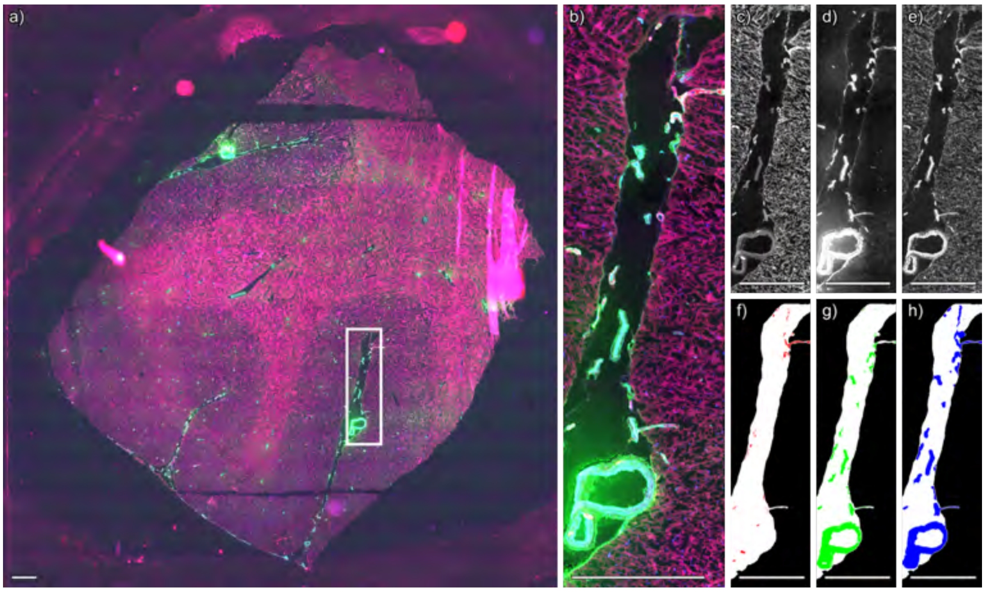


**Figure S47** Segmentation of immunofluorescence microscopy image of case 7, technical replicate 2, positive ion mode. (a) Whole slide immunofluorescence microscopy image. (b) Immunofluorescence image of region acquired by MALDI IMS, shown in a) as a white box. (c) Thiazine red stains amyloid, (d) anti-actin antibodies label smooth muscle, and (e) anti-collagen IV antibodies label vascular basement membranes. Segmentations for (f) amyloid, (g) smooth muscle cells, and (h) collagen IV. Per-channel standardization was applied across the entire dataset to ensure that color intensities are directly comparable across Figures S9-S82. Scale bar is equal to 1 mm.

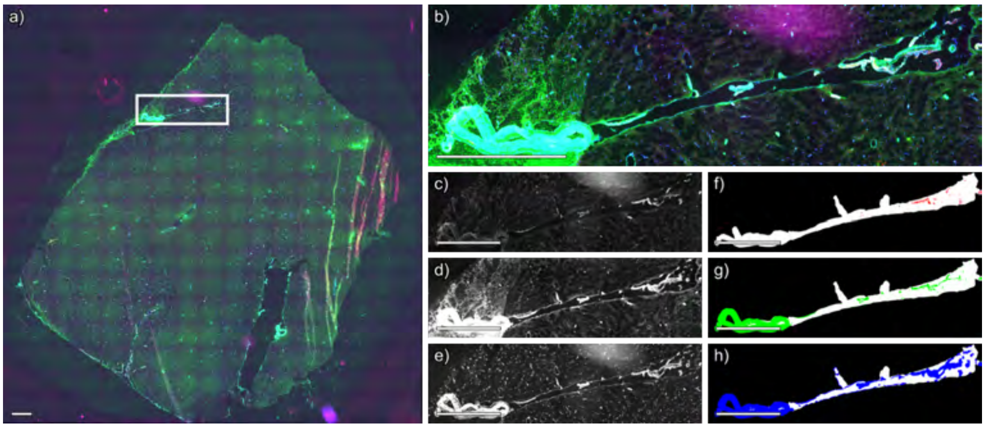


**Figure S48** Segmentation of immunofluorescence microscopy image of case 7, technical replicate 3, negative ion mode. (a) Whole slide immunofluorescence microscopy image. (b) Immunofluorescence image of region acquired by MALDI IMS, shown in a) as a white box. (c) Thiazine red stains amyloid, (d) anti-actin antibodies label smooth muscle, and (e) anti-collagen IV antibodies label vascular basement membranes. Segmentations for (f) amyloid, (g) smooth muscle cells, and (h) collagen IV. Per-channel standardization was applied across the entire dataset to ensure that color intensities are directly comparable across Figures S9-S82. Scale bar is equal to 1 mm.

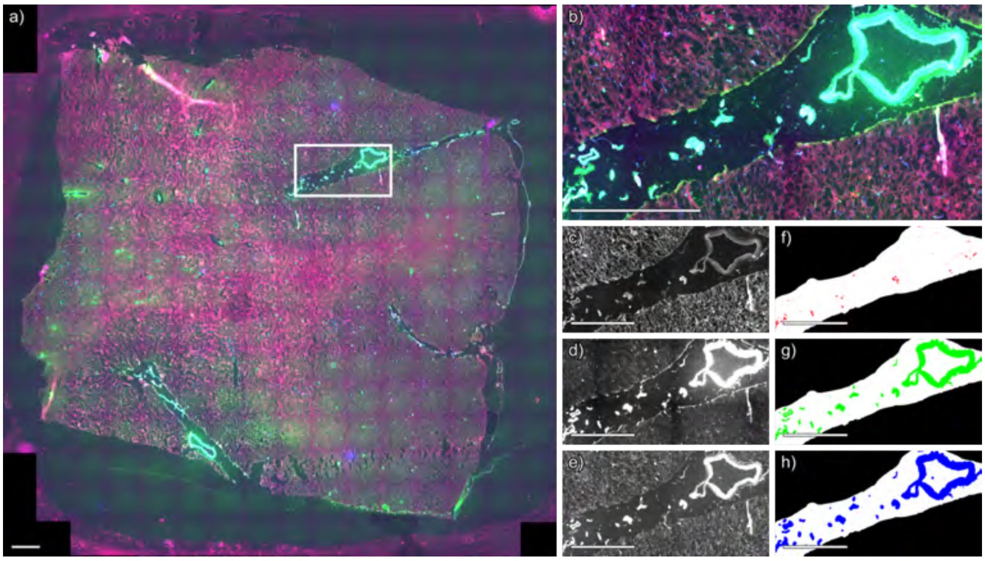


**Figure S49** Segmentation of immunofluorescence microscopy image of case 7, technical replicate 3, positive ion mode. (a) Whole slide immunofluorescence microscopy image. (b) Immunofluorescence image of region acquired by MALDI IMS, shown in a) as a white box. (c) Thiazine red stains amyloid, (d) anti-actin antibodies label smooth muscle, and (e) anti-collagen IV antibodies label vascular basement membranes. Segmentations for (f) amyloid, (g) smooth muscle cells, and (h) collagen IV. Per-channel standardization was applied across the entire dataset to ensure that color intensities are directly comparable across Figures S9-S82. Scale bar is equal to 1 mm.

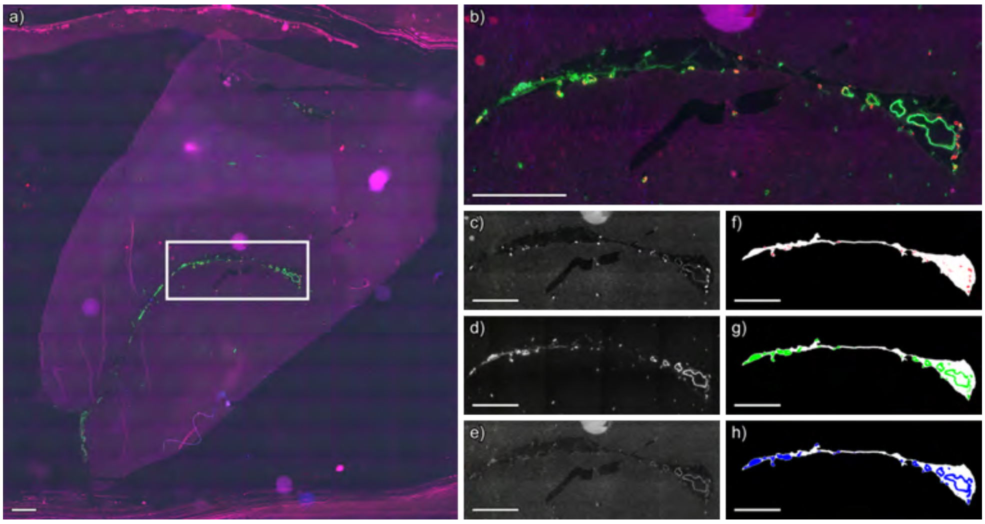


**Figure S50** Segmentation of immunofluorescence microscopy image of case 8, technical replicate 1, negative ion mode. (a) Whole slide immunofluorescence microscopy image. (b) Immunofluorescence image of region acquired by MALDI IMS, shown in a) as a white box. (c) Thiazine red stains amyloid, (d) anti-actin antibodies label smooth muscle, and (e) anti-collagen IV antibodies label vascular basement membranes. Segmentations for (f) amyloid, (g) smooth muscle cells, and (h) collagen IV. Per-channel standardization was applied across the entire dataset to ensure that color intensities are directly comparable across Figures S9-S82. Scale bar is equal to 1 mm.

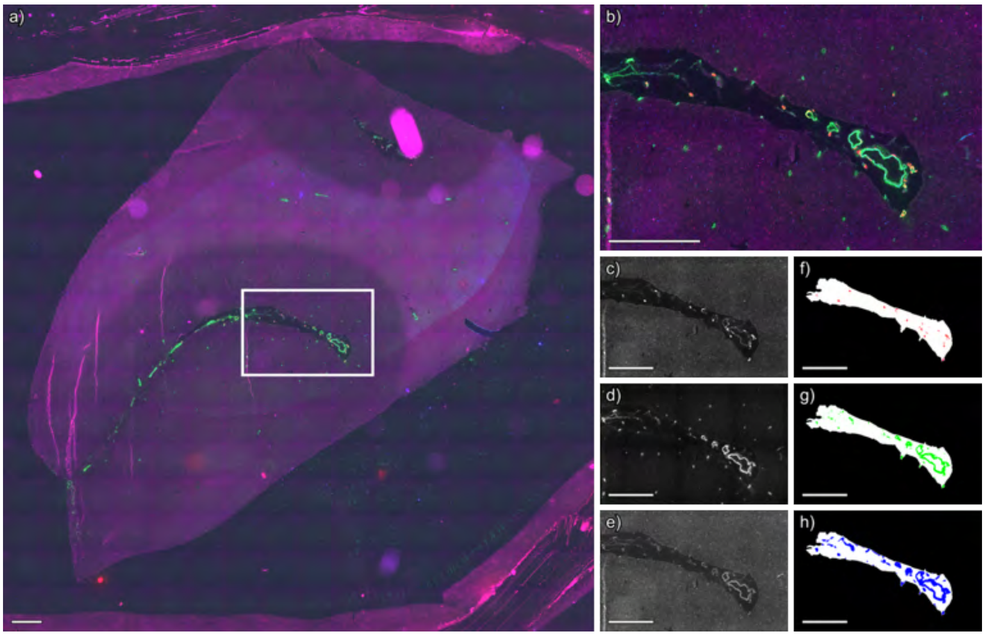


**Figure S51** Segmentation of immunofluorescence microscopy image of case 8, technical replicate 1, positive ion mode. (a) Whole slide immunofluorescence microscopy image. (b) Immunofluorescence image of region acquired by MALDI IMS, shown in a) as a white box. (c) Thiazine red stains amyloid, (d) anti-actin antibodies label smooth muscle, and (e) anti-collagen IV antibodies label vascular basement membranes. Segmentations for (f) amyloid, (g) smooth muscle cells, and (h) collagen IV. Per-channel standardization was applied across the entire dataset to ensure that color intensities are directly comparable across Figures S9-S82. Scale bar is equal to 1 mm.

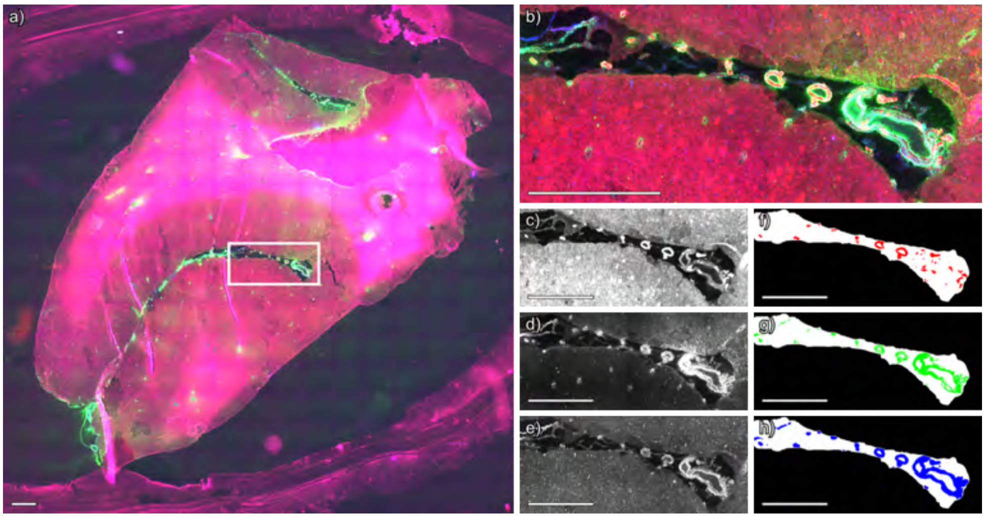


**Figure S52** Segmentation of immunofluorescence microscopy image of case 8, technical replicate 2, negative ion mode. (a) Whole slide immunofluorescence microscopy image. (b) Immunofluorescence image of region acquired by MALDI IMS, shown in a) as a white box. (c) Thiazine red stains amyloid, (d) anti-actin antibodies label smooth muscle, and (e) anti-collagen IV antibodies label vascular basement membranes. Segmentations for (f) amyloid, (g) smooth muscle cells, and (h) collagen IV. Per-channel standardization was applied across the entire dataset to ensure that color intensities are directly comparable across Figures S9-S82. Scale bar is equal to 1 mm.

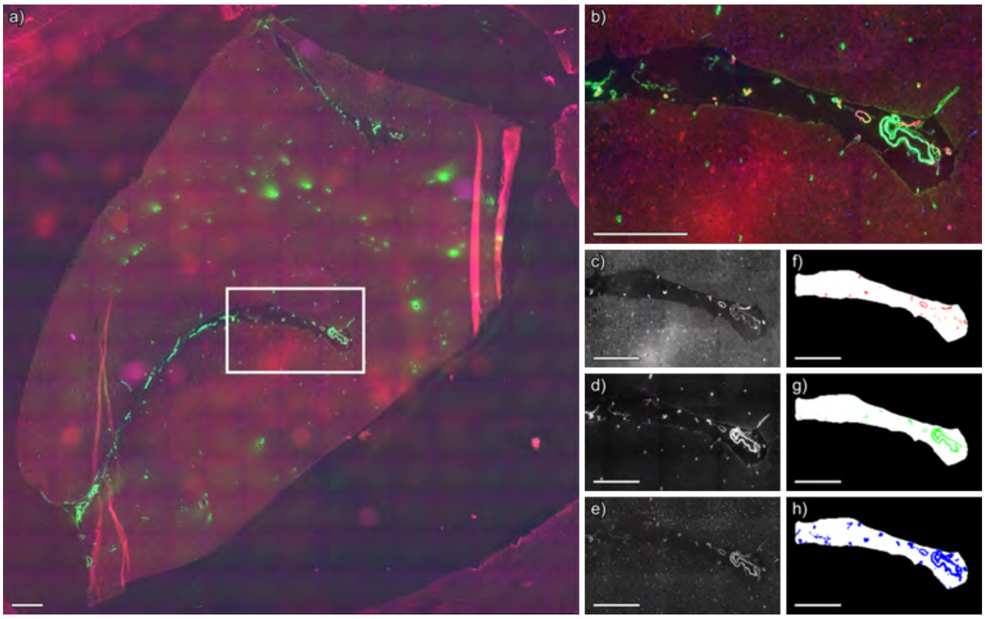


**Figure S53** Segmentation of immunofluorescence microscopy image of case 8, technical replicate 2, positive ion mode. (a) Whole slide immunofluorescence microscopy image. (b) Immunofluorescence image of region acquired by MALDI IMS, shown in a) as a white box. (c) Thiazine red stains amyloid, (d) anti-actin antibodies label smooth muscle, and (e) anti-collagen IV antibodies label vascular basement membranes. Segmentations for (f) amyloid, (g) smooth muscle cells, and (h) collagen IV. Per-channel standardization was applied across the entire dataset to ensure that color intensities are directly comparable across Figures S9-S82. Scale bar is equal to 1 mm.

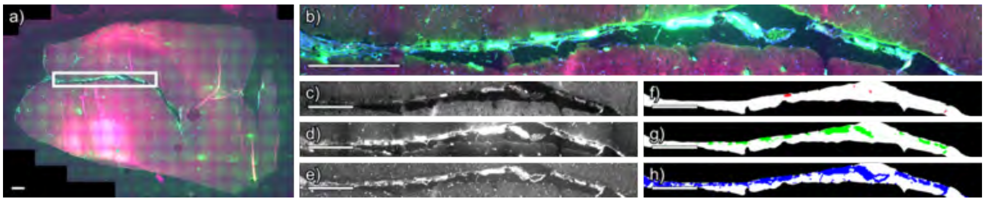


**Figure S54** Segmentation of immunofluorescence microscopy image of case 8, technical replicate 3, negative ion mode. (a) Whole slide immunofluorescence microscopy image. (b) Immunofluorescence image of region acquired by MALDI IMS, shown in a) as a white box. (c) Thiazine red stains amyloid, (d) anti-actin antibodies label smooth muscle, and (e) anti-collagen IV antibodies label vascular basement membranes. Segmentations for (f) amyloid, (g) smooth muscle cells, and (h) collagen IV. Per-channel standardization was applied across the entire dataset to ensure that color intensities are directly comparable across Figures S9-S82. Scale bar is equal to 1 mm.

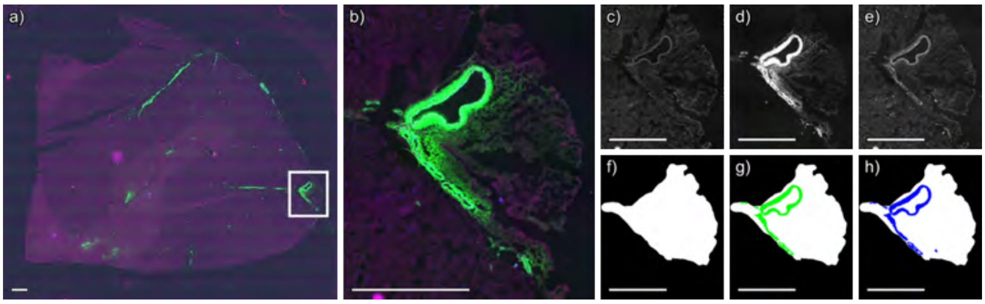


**Figure S55** Segmentation of immunofluorescence microscopy image of case 9, technical replicate 1, negative ion mode. (a) Whole slide immunofluorescence microscopy image. (b) Immunofluorescence image of region acquired by MALDI IMS, shown in a) as a white box. (c) Thiazine red stains amyloid, (d) anti-actin antibodies label smooth muscle, and (e) anti-collagen IV antibodies label vascular basement membranes. Segmentations for (f) amyloid, (g) smooth muscle cells, and (h) collagen IV. Per-channel standardization was applied across the entire dataset to ensure that color intensities are directly comparable across Figures S9-S82. Scale bar is equal to 1 mm.

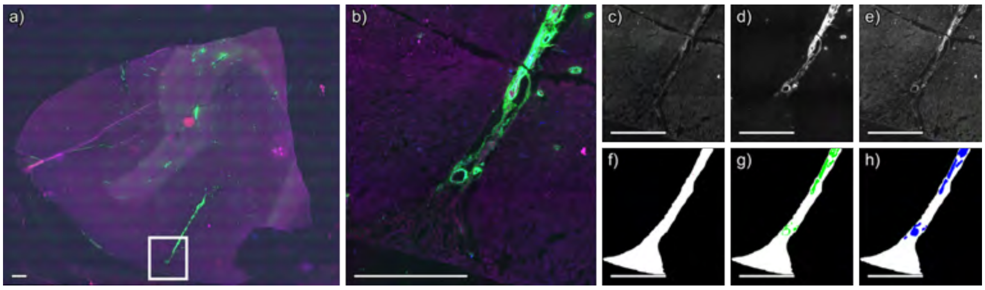


**Figure S56** Segmentation of immunofluorescence microscopy image of case 9, technical replicate 1, positive ion mode. (a) Whole slide immunofluorescence microscopy image. (b) Immunofluorescence image of region acquired by MALDI IMS, shown in a) as a white box. (c) Thiazine red stains amyloid, (d) anti-actin antibodies label smooth muscle, and (e) anti-collagen IV antibodies label vascular basement membranes. Segmentations for (f) amyloid, (g) smooth muscle cells, and (h) collagen IV. Per-channel standardization was applied across the entire dataset to ensure that color intensities are directly comparable across Figures S9-S82. Scale bar is equal to 1 mm.

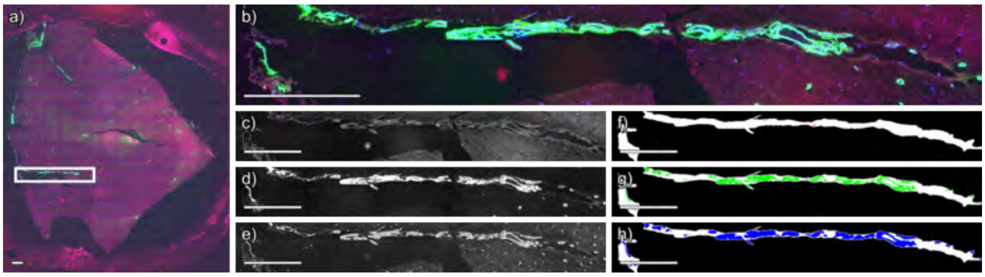


**Figure S57** Segmentation of immunofluorescence microscopy image of case 9, technical replicate 2, negative ion mode. (a) Whole slide immunofluorescence microscopy image. (b) Immunofluorescence image of region acquired by MALDI IMS, shown in a) as a white box. (c) Thiazine red stains amyloid, (d) anti-actin antibodies label smooth muscle, and (e) anti-collagen IV antibodies label vascular basement membranes. Segmentations for (f) amyloid, (g) smooth muscle cells, and (h) collagen IV. Per-channel standardization was applied across the entire dataset to ensure that color intensities are directly comparable across Figures S9-S82. Scale bar is equal to 1 mm.

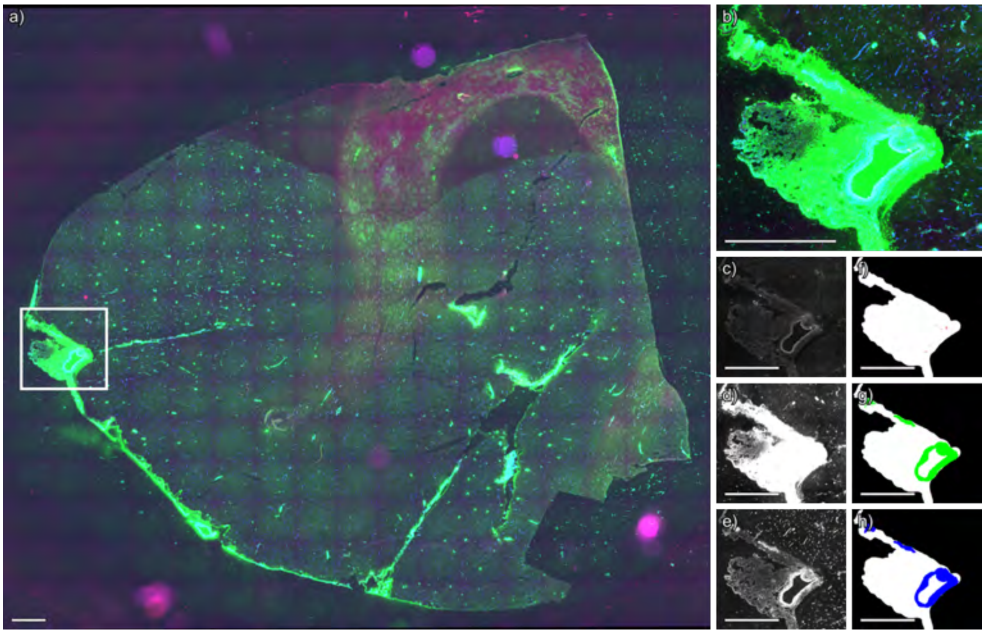


**Figure S58** Segmentation of immunofluorescence microscopy image of case 9, technical replicate 3, negative ion mode. (a) Whole slide immunofluorescence microscopy image. (b) Immunofluorescence image of region acquired by MALDI IMS, shown in a) as a white box. (c) Thiazine red stains amyloid, (d) anti-actin antibodies label smooth muscle, and (e) anti-collagen IV antibodies label vascular basement membranes. Segmentations for (f) amyloid, (g) smooth muscle cells, and (h) collagen IV. Per-channel standardization was applied across the entire dataset to ensure that color intensities are directly comparable across Figures S9-S82. Scale bar is equal to 1 mm.

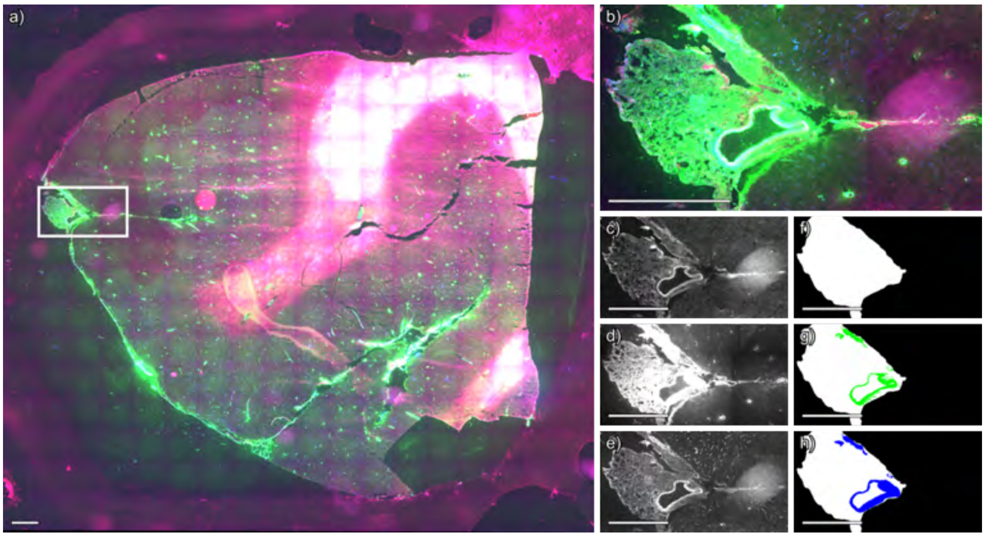


**Figure S59** Segmentation of immunofluorescence microscopy image of case 9, technical replicate 3, positive ion mode. (a) Whole slide immunofluorescence microscopy image. (b) Immunofluorescence image of region acquired by MALDI IMS, shown in a) as a white box. (c) Thiazine red stains amyloid, (d) anti-actin antibodies label smooth muscle, and (e) anti-collagen IV antibodies label vascular basement membranes. Segmentations for (f) amyloid, (g) smooth muscle cells, and (h) collagen IV. Per-channel standardization was applied across the entire dataset to ensure that color intensities are directly comparable across Figures S9-S82. Scale bar is equal to 1 mm.

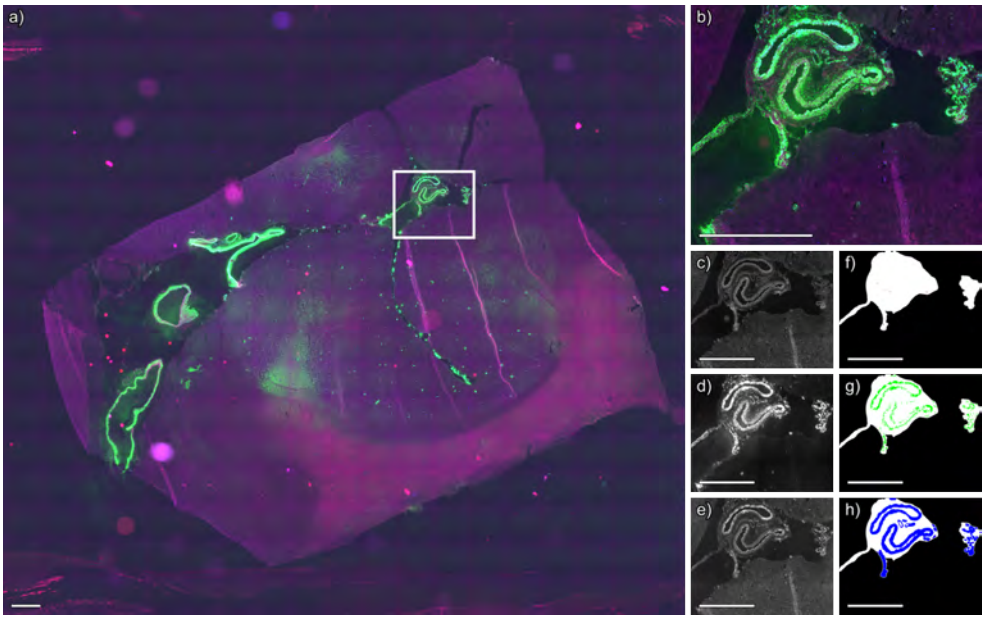


**Figure S60** Segmentation of immunofluorescence microscopy image of case 10, technical replicate 1, negative ion mode. (a) Whole slide immunofluorescence microscopy image. (b) Immunofluorescence image of region acquired by MALDI IMS, shown in a) as a white box. (c) Thiazine red stains amyloid, (d) anti-actin antibodies label smooth muscle, and (e) anti-collagen IV antibodies label vascular basement membranes. Segmentations for (f) amyloid, (g) smooth muscle cells, and (h) collagen IV. Per-channel standardization was applied across the entire dataset to ensure that color intensities are directly comparable across Figures S9-S82. Scale bar is equal to 1 mm.

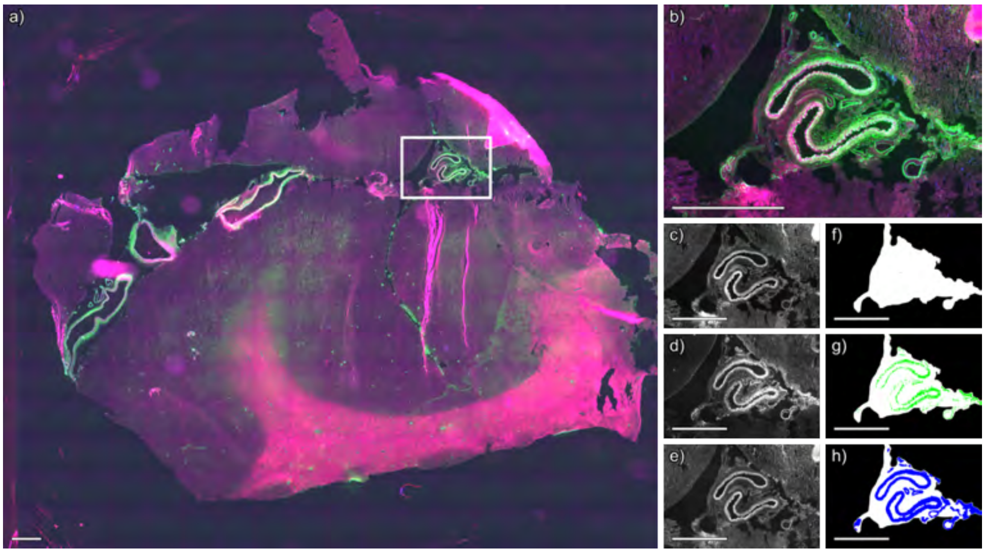


**Figure S61** Segmentation of immunofluorescence microscopy image of case 10, technical replicate 1, positive ion mode. (a) Whole slide immunofluorescence microscopy image. (b) Immunofluorescence image of region acquired by MALDI IMS, shown in a) as a white box. (c) Thiazine red stains amyloid, (d) anti-actin antibodies label smooth muscle, and (e) anti-collagen IV antibodies label vascular basement membranes. Segmentations for (f) amyloid, (g) smooth muscle cells, and (h) collagen IV. Per-channel standardization was applied across the entire dataset to ensure that color intensities are directly comparable across Figures S9-S82. Scale bar is equal to 1 mm.

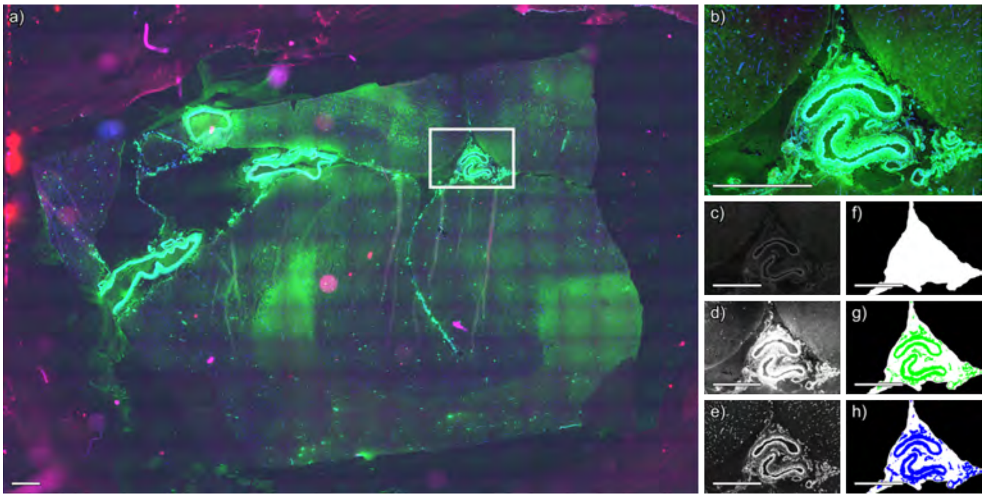


**Figure S62** Segmentation of immunofluorescence microscopy image of case 10, technical replicate 2, negative ion mode. (a) Whole slide immunofluorescence microscopy image. (b) Immunofluorescence image of region acquired by MALDI IMS, shown in a) as a white box. (c) Thiazine red stains amyloid, (d) anti-actin antibodies label smooth muscle, and (e) anti-collagen IV antibodies label vascular basement membranes. Segmentations for (f) amyloid, (g) smooth muscle cells, and (h) collagen IV. Per-channel standardization was applied across the entire dataset to ensure that color intensities are directly comparable across Figures S9-S82. Scale bar is equal to 1 mm.

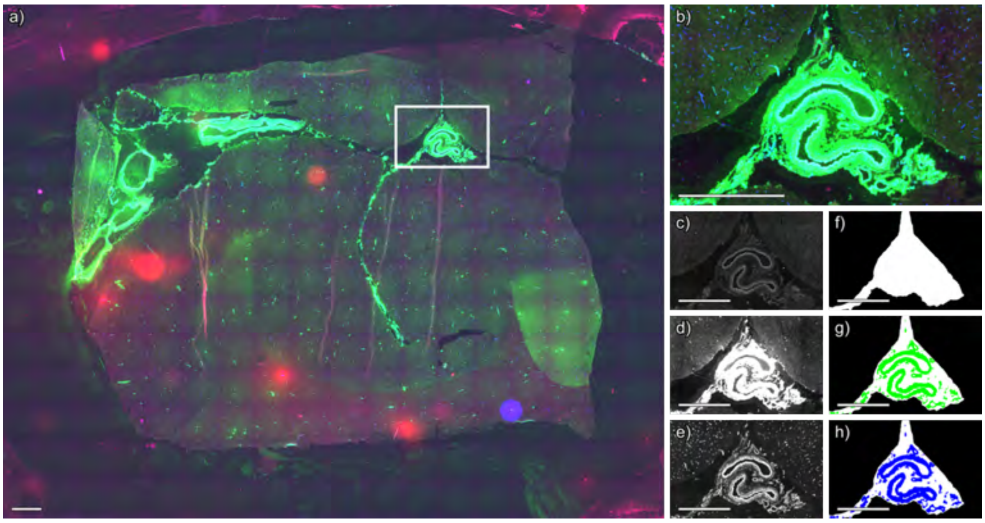


**Figure S63** Segmentation of immunofluorescence microscopy image of case 10, technical replicate 2, positive ion mode. (a) Whole slide immunofluorescence microscopy image. (b) Immunofluorescence image of region acquired by MALDI IMS, shown in a) as a white box. (c) Thiazine red stains amyloid, (d) anti-actin antibodies label smooth muscle, and (e) anti-collagen IV antibodies label vascular basement membranes. Segmentations for (f) amyloid, (g) smooth muscle cells, and (h) collagen IV. Per-channel standardization was applied across the entire dataset to ensure that color intensities are directly comparable across Figures S9-S82. Scale bar is equal to 1 mm.

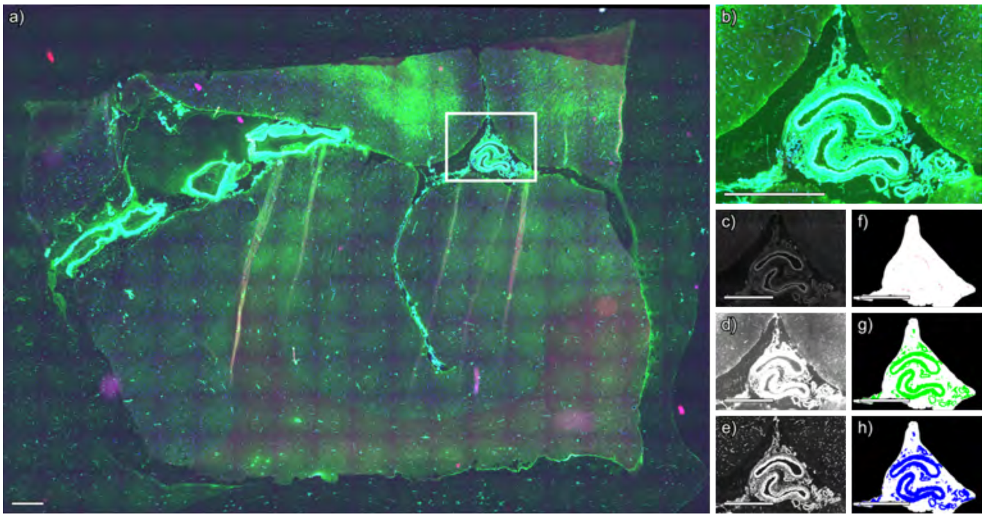


**Figure S64** Segmentation of immunofluorescence microscopy image of case 10, technical replicate 3, negative ion mode. (a) Whole slide immunofluorescence microscopy image. (b) Immunofluorescence image of region acquired by MALDI IMS, shown in a) as a white box. (c) Thiazine red stains amyloid, (d) anti-actin antibodies label smooth muscle, and (e) anti-collagen IV antibodies label vascular basement membranes. Segmentations for (f) amyloid, (g) smooth muscle cells, and (h) collagen IV. Per-channel standardization was applied across the entire dataset to ensure that color intensities are directly comparable across Figures S9-S82. Scale bar is equal to 1 mm.

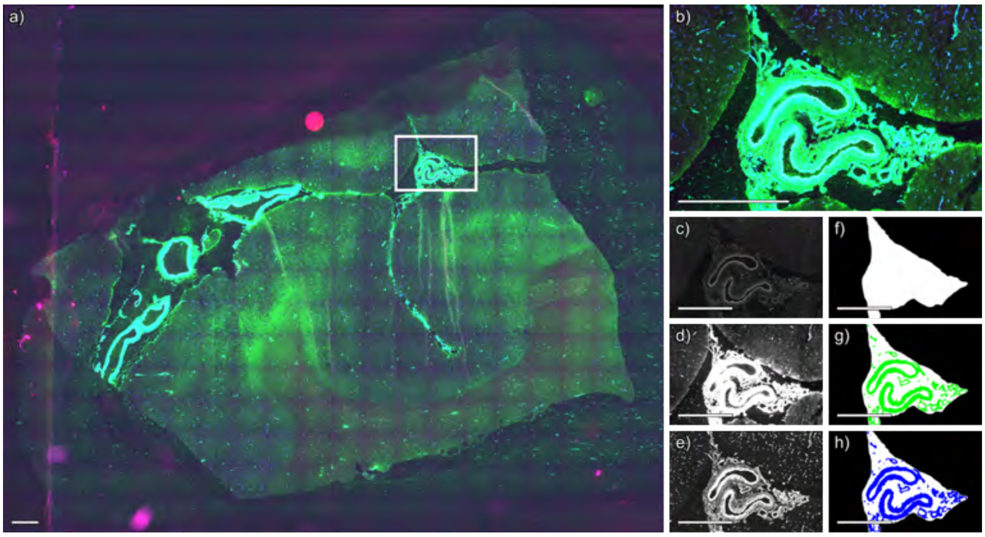


**Figure S65** Segmentation of immunofluorescence microscopy image of case 10, technical replicate 3, positive ion mode. (a) Whole slide immunofluorescence microscopy image. (b) Immunofluorescence image of region acquired by MALDI IMS, shown in a) as a white box. (c) Thiazine red stains amyloid, (d) anti-actin antibodies label smooth muscle, and (e) anti-collagen IV antibodies label vascular basement membranes. Segmentations for (f) amyloid, (g) smooth muscle cells, and (h) collagen IV. Per-channel standardization was applied across the entire dataset to ensure that color intensities are directly comparable across Figures S9-S82. Scale bar is equal to 1 mm.

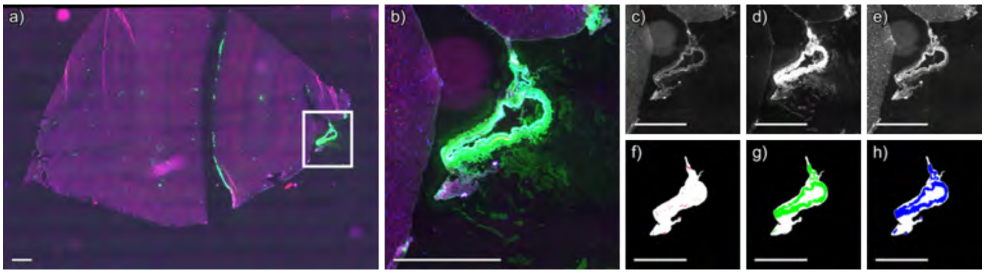


**Figure S66** Segmentation of immunofluorescence microscopy image of case 11, technical replicate 1, positive ion mode. (a) Whole slide immunofluorescence microscopy image. (b) Immunofluorescence image of region acquired by MALDI IMS, shown in a) as a white box. (c) Thiazine red stains amyloid, (d) anti-actin antibodies label smooth muscle, and (e) anti-collagen IV antibodies label vascular basement membranes. Segmentations for (f) amyloid, (g) smooth muscle cells, and (h) collagen IV. Per-channel standardization was applied across the entire dataset to ensure that color intensities are directly comparable across Figures S9-S82. Scale bar is equal to 1 mm.

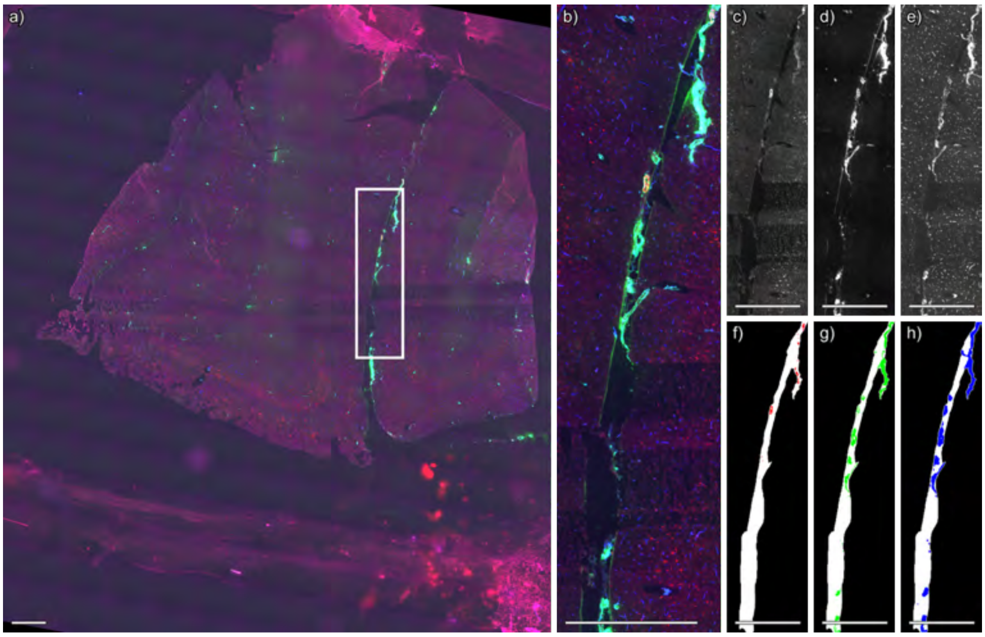


**Figure S67** Segmentation of immunofluorescence microscopy image of case 11, technical replicate 2, negative ion mode. (a) Whole slide immunofluorescence microscopy image. (b) Immunofluorescence image of region acquired by MALDI IMS, shown in a) as a white box. (c) Thiazine red stains amyloid, (d) anti-actin antibodies label smooth muscle, and (e) anti-collagen IV antibodies label vascular basement membranes. Segmentations for (f) amyloid, (g) smooth muscle cells, and (h) collagen IV. Per-channel standardization was applied across the entire dataset to ensure that color intensities are directly comparable across Figures S9-S82. Scale bar is equal to 1 mm.

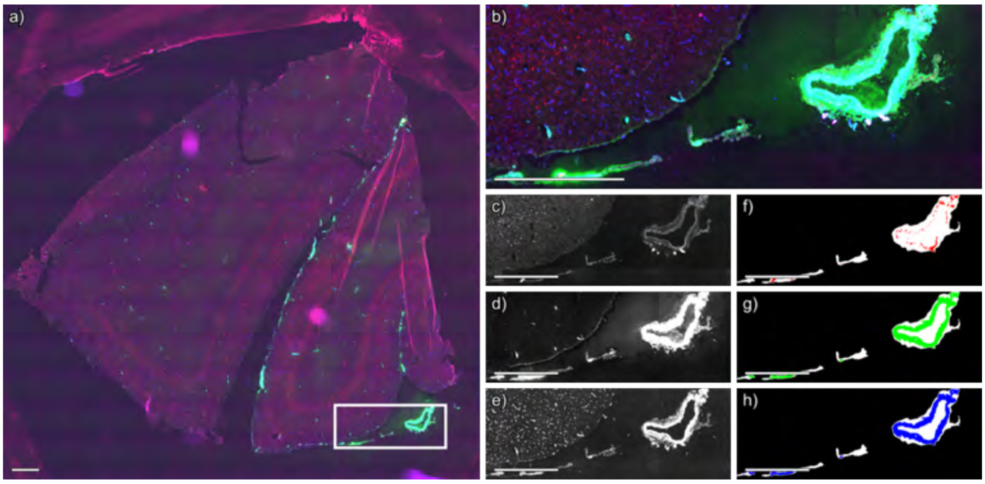


**Figure S68** Segmentation of immunofluorescence microscopy image of case 11, technical replicate 2, positive ion mode. (a) Whole slide immunofluorescence microscopy image. (b) Immunofluorescence image of region acquired by MALDI IMS, shown in a) as a white box. (c) Thiazine red stains amyloid, (d) anti-actin antibodies label smooth muscle, and (e) anti-collagen IV antibodies label vascular basement membranes. Segmentations for (f) amyloid, (g) smooth muscle cells, and (h) collagen IV. Per-channel standardization was applied across the entire dataset to ensure that color intensities are directly comparable across Figures S9-S82. Scale bar is equal to 1 mm.

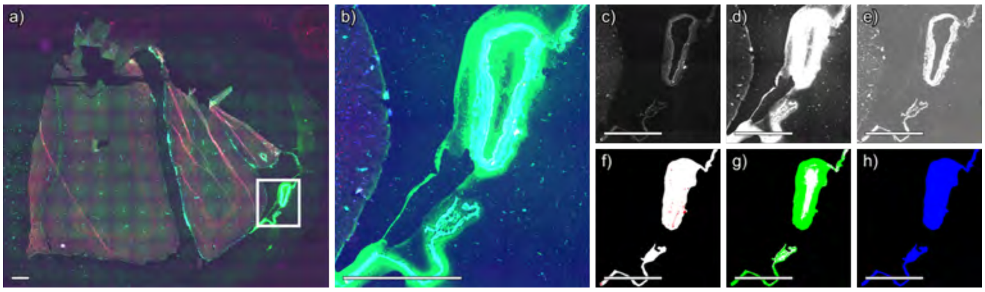


**Figure S69** Segmentation of immunofluorescence microscopy image of case 11, technical replicate 3, negative ion mode. (a) Whole slide immunofluorescence microscopy image. (b) Immunofluorescence image of region acquired by MALDI IMS, shown in a) as a white box. (c) Thiazine red stains amyloid, (d) anti-actin antibodies label smooth muscle, and (e) anti-collagen IV antibodies label vascular basement membranes. Segmentations for (f) amyloid, (g) smooth muscle cells, and (h) collagen IV. Per-channel standardization was applied across the entire dataset to ensure that color intensities are directly comparable across Figures S9-S82. Scale bar is equal to 1 mm.

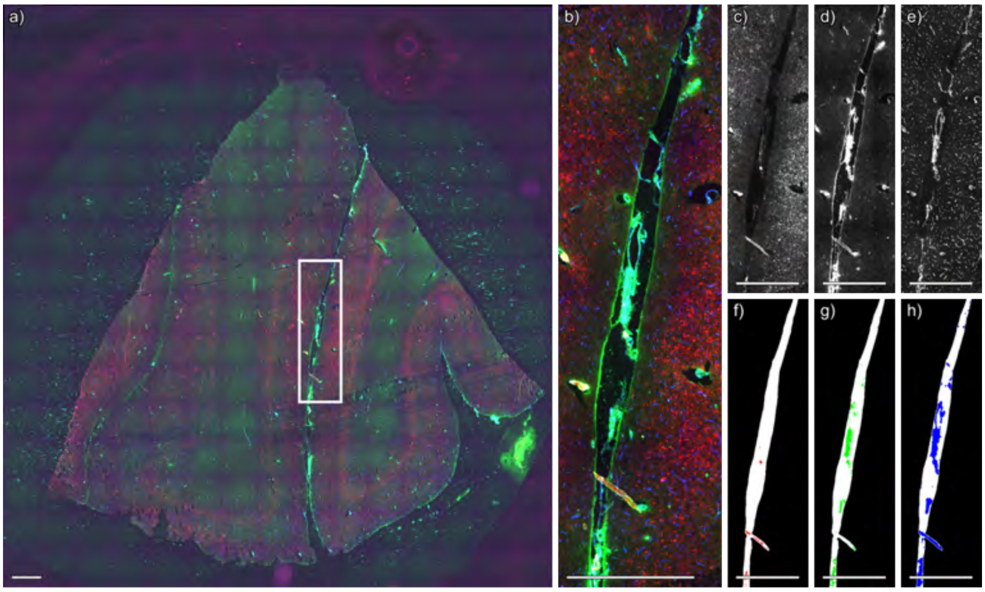


**Figure S70** Segmentation of immunofluorescence microscopy image of case 11, technical replicate 3, positive ion mode. (a) Whole slide immunofluorescence microscopy image. (b) Immunofluorescence image of region acquired by MALDI IMS, shown in a) as a white box. (c) Thiazine red stains amyloid, (d) anti-actin antibodies label smooth muscle, and (e) anti-collagen IV antibodies label vascular basement membranes. Segmentations for (f) amyloid, (g) smooth muscle cells, and (h) collagen IV. Per-channel standardization was applied across the entire dataset to ensure that color intensities are directly comparable across Figures S9-S82. Scale bar is equal to 1 mm.

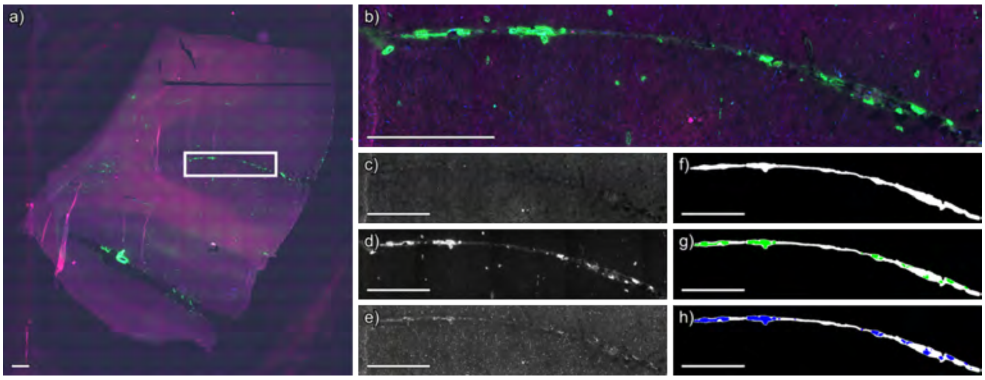


**Figure S71** Segmentation of immunofluorescence microscopy image of case 12, technical replicate 1, negative ion mode. (a) Whole slide immunofluorescence microscopy image. (b) Immunofluorescence image of region acquired by MALDI IMS, shown in a) as a white box. (c) Thiazine red stains amyloid, (d) anti-actin antibodies label smooth muscle, and (e) anti-collagen IV antibodies label vascular basement membranes. Segmentations for (f) amyloid, (g) smooth muscle cells, and (h) collagen IV. Per-channel standardization was applied across the entire dataset to ensure that color intensities are directly comparable across Figures S9-S82. Scale bar is equal to 1 mm.

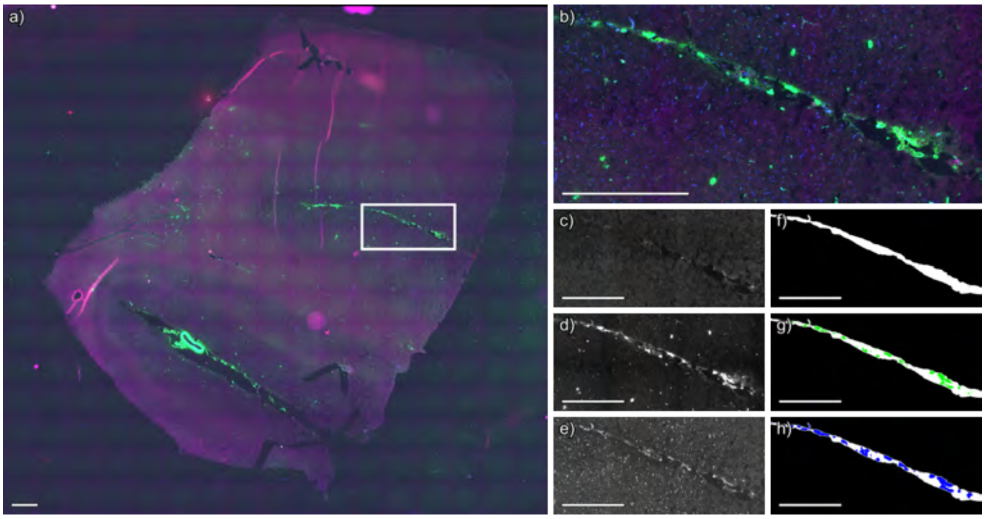


**Figure S72** Segmentation of immunofluorescence microscopy image of case 12, technical replicate 1, positive ion mode. (a) Whole slide immunofluorescence microscopy image. (b) Immunofluorescence image of region acquired by MALDI IMS, shown in a) as a white box. (c) Thiazine red stains amyloid, (d) anti-actin antibodies label smooth muscle, and (e) anti-collagen IV antibodies label vascular basement membranes. Segmentations for (f) amyloid, (g) smooth muscle cells, and (h) collagen IV. Per-channel standardization was applied across the entire dataset to ensure that color intensities are directly comparable across Figures S9-S82. Scale bar is equal to 1 mm.

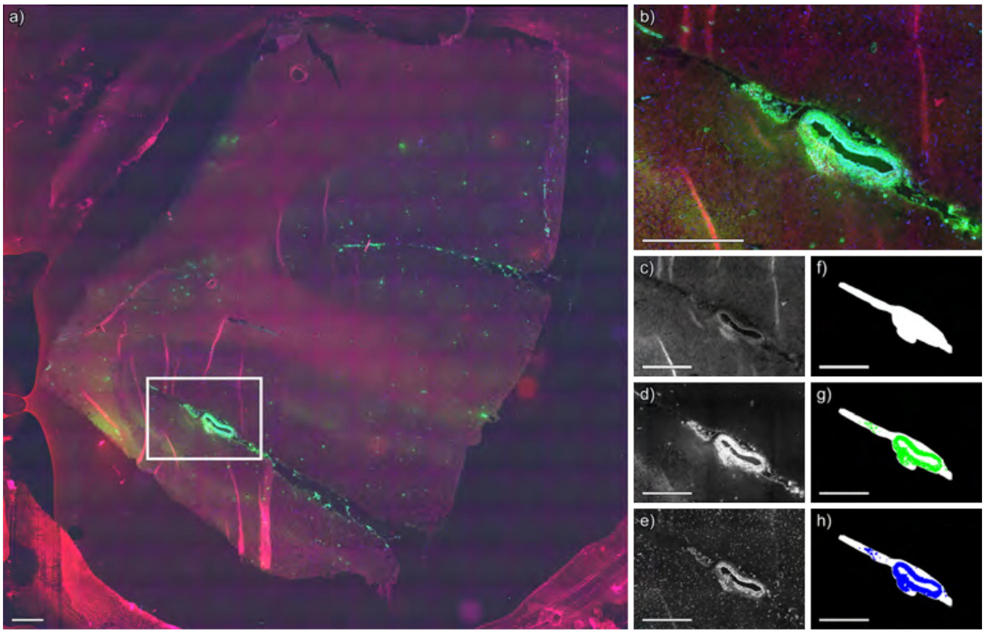


**Figure S73** Segmentation of immunofluorescence microscopy image of case 12, technical replicate 2, negative ion mode. (a) Whole slide immunofluorescence microscopy image. (b) Immunofluorescence image of region acquired by MALDI IMS, shown in a) as a white box. (c) Thiazine red stains amyloid, (d) anti-actin antibodies label smooth muscle, and (e) anti-collagen IV antibodies label vascular basement membranes. Segmentations for (f) amyloid, (g) smooth muscle cells, and (h) collagen IV. Per-channel standardization was applied across the entire dataset to ensure that color intensities are directly comparable across Figures S9-S82. Scale bar is equal to 1 mm.

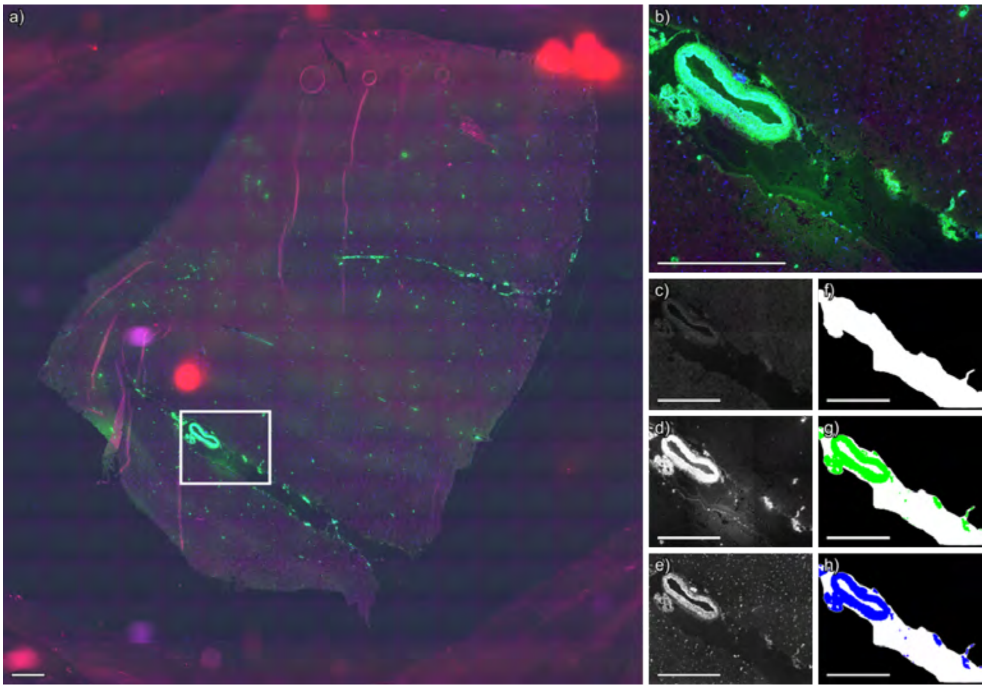


**Figure S74** Segmentation of immunofluorescence microscopy image of case 12, technical replicate 2, positive ion mode. (a) Whole slide immunofluorescence microscopy image. (b) Immunofluorescence image of region acquired by MALDI IMS, shown in a) as a white box. (c) Thiazine red stains amyloid, (d) anti-actin antibodies label smooth muscle, and (e) anti-collagen IV antibodies label vascular basement membranes. Segmentations for (f) amyloid, (g) smooth muscle cells, and (h) collagen IV. Per-channel standardization was applied across the entire dataset to ensure that color intensities are directly comparable across Figures S9-S82. Scale bar is equal to 1 mm.

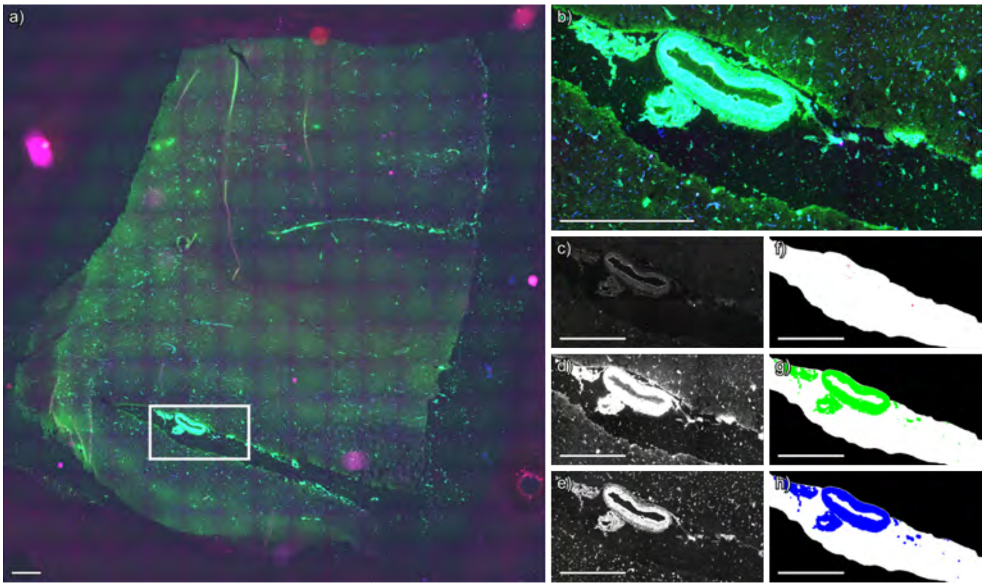


**Figure S75** Segmentation of immunofluorescence microscopy image of case 12, technical replicate 3, negative ion mode. (a) Whole slide immunofluorescence microscopy image. (b) Immunofluorescence image of region acquired by MALDI IMS, shown in a) as a white box. (c) Thiazine red stains amyloid, (d) anti-actin antibodies label smooth muscle, and (e) anti-collagen IV antibodies label vascular basement membranes. Segmentations for (f) amyloid, (g) smooth muscle cells, and (h) collagen IV. Per-channel standardization was applied across the entire dataset to ensure that color intensities are directly comparable across Figures S9-S82. Scale bar is equal to 1 mm.

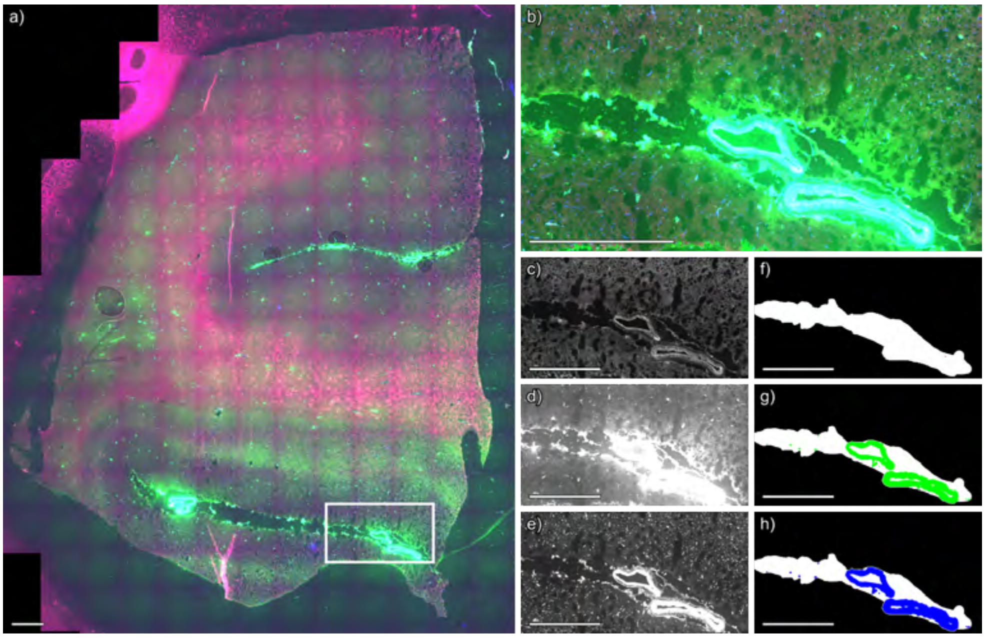


**Figure S76** Segmentation of immunofluorescence microscopy image of case 12, technical replicate 3, positive ion mode. (a) Whole slide immunofluorescence microscopy image. (b) Immunofluorescence image of region acquired by MALDI IMS, shown in a) as a white box. (c) Thiazine red stains amyloid, (d) anti-actin antibodies label smooth muscle, and (e) anti-collagen IV antibodies label vascular basement membranes. Segmentations for (f) amyloid, (g) smooth muscle cells, and (h) collagen IV. Per-channel standardization was applied across the entire dataset to ensure that color intensities are directly comparable across Figures S9-S82. Scale bar is equal to 1 mm.

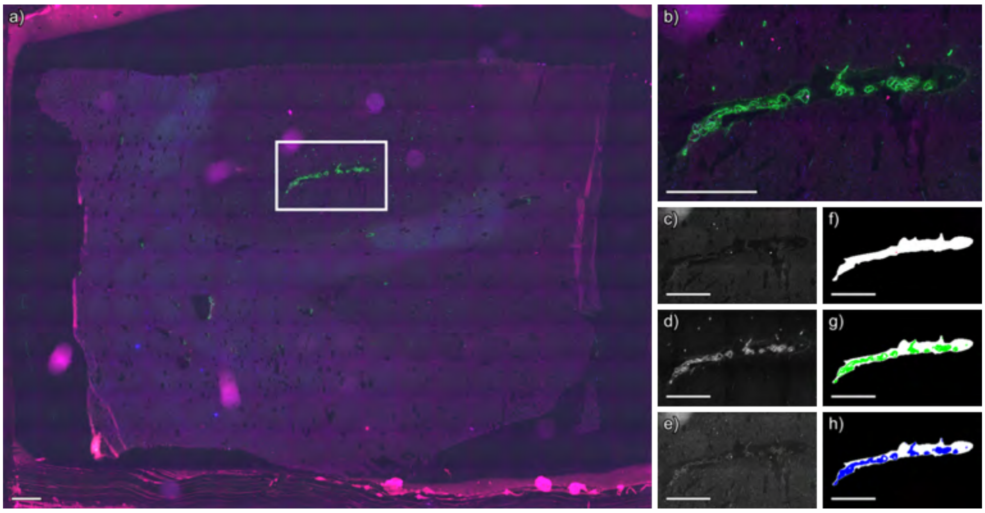


**Figure S77** Segmentation of immunofluorescence microscopy image of case 13, technical replicate 1, negative ion mode. (a) Whole slide immunofluorescence microscopy image. (b) Immunofluorescence image of region acquired by MALDI IMS, shown in a) as a white box. (c) Thiazine red stains amyloid, (d) anti-actin antibodies label smooth muscle, and (e) anti-collagen IV antibodies label vascular basement membranes. Segmentations for (f) amyloid, (g) smooth muscle cells, and (h) collagen IV. Per-channel standardization was applied across the entire dataset to ensure that color intensities are directly comparable across Figures S9-S82. Scale bar is equal to 1 mm.

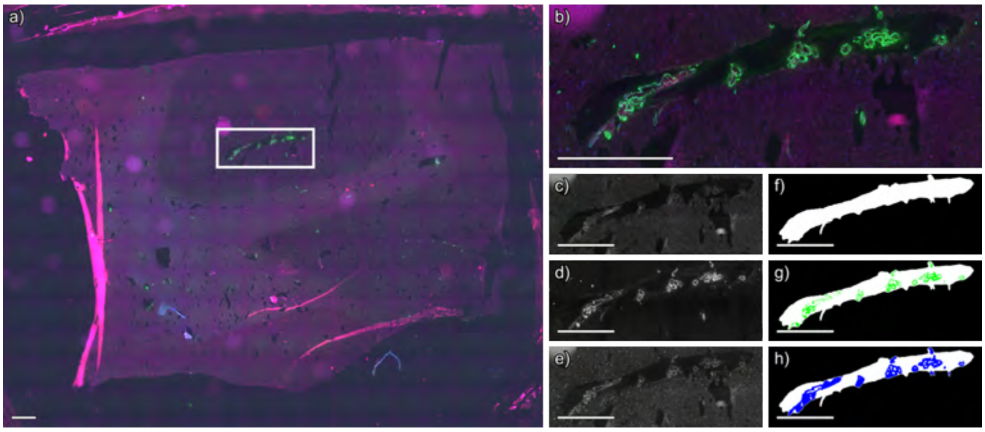


**Figure S78** Segmentation of immunofluorescence microscopy image of case 13, technical replicate 1, positive ion mode. (a) Whole slide immunofluorescence microscopy image. (b) Immunofluorescence image of region acquired by MALDI IMS, shown in a) as a white box. (c) Thiazine red stains amyloid, (d) anti-actin antibodies label smooth muscle, and (e) anti-collagen IV antibodies label vascular basement membranes. Segmentations for (f) amyloid, (g) smooth muscle cells, and (h) collagen IV. Per-channel standardization was applied across the entire dataset to ensure that color intensities are directly comparable across Figures S9-S82. Scale bar is equal to 1 mm.

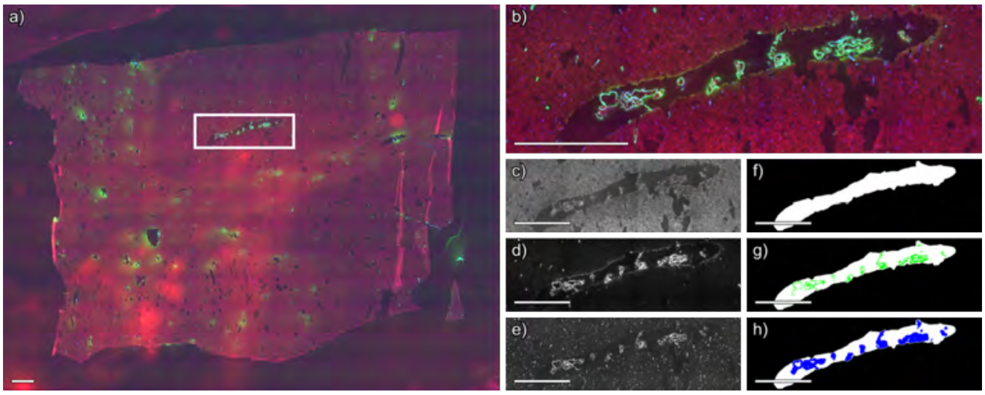


**Figure S79** Segmentation of immunofluorescence microscopy image of case 13, technical replicate 2, negative ion mode. (a) Whole slide immunofluorescence microscopy image. (b) Immunofluorescence image of region acquired by MALDI IMS, shown in a) as a white box. (c) Thiazine red stains amyloid, (d) anti-actin antibodies label smooth muscle, and (e) anti-collagen IV antibodies label vascular basement membranes. Segmentations for (f) amyloid, (g) smooth muscle cells, and (h) collagen IV. Per-channel standardization was applied across the entire dataset to ensure that color intensities are directly comparable across Figures S9-S82. Scale bar is equal to 1 mm.

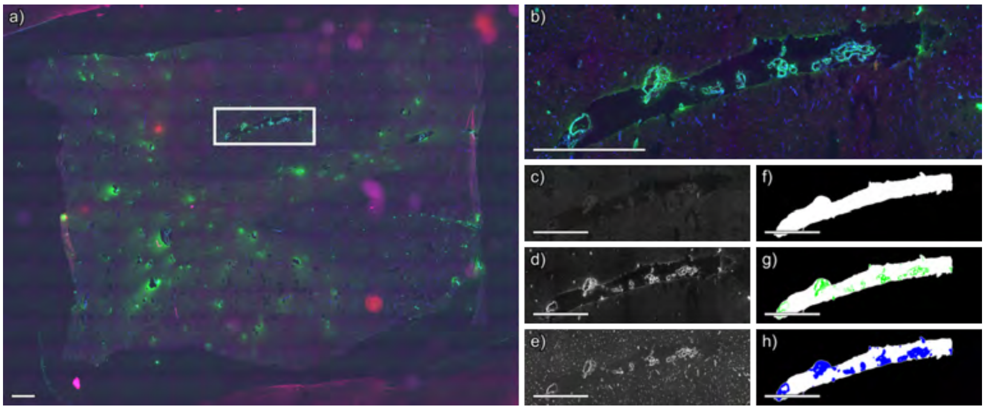


**Figure S80** Segmentation of immunofluorescence microscopy image of case 13, technical replicate 2, positive ion mode. (a) Whole slide immunofluorescence microscopy image. (b) Immunofluorescence image of region acquired by MALDI IMS, shown in a) as a white box. (c) Thiazine red stains amyloid, (d) anti-actin antibodies label smooth muscle, and (e) anti-collagen IV antibodies label vascular basement membranes. Segmentations for (f) amyloid, (g) smooth muscle cells, and (h) collagen IV. Per-channel standardization was applied across the entire dataset to ensure that color intensities are directly comparable across Figures S9-S82. Scale bar is equal to 1 mm.

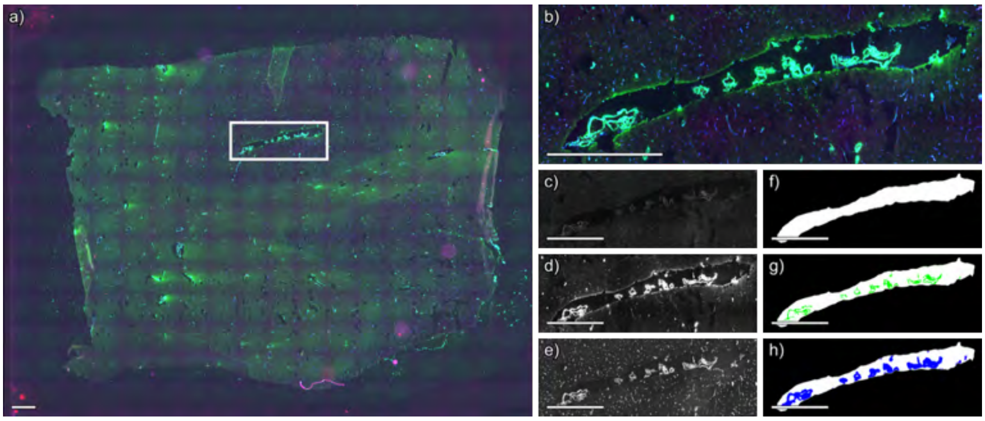


**Figure S81** Segmentation of immunofluorescence microscopy image of case 13, technical replicate 3, negative ion mode. (a) Whole slide immunofluorescence microscopy image. (b) Immunofluorescence image of region acquired by MALDI IMS, shown in a) as a white box. (c) Thiazine red stains amyloid, (d) anti-actin antibodies label smooth muscle, and (e) anti-collagen IV antibodies label vascular basement membranes. Segmentations for (f) amyloid, (g) smooth muscle cells, and (h) collagen IV. Per-channel standardization was applied across the entire dataset to ensure that color intensities are directly comparable across Figures S9-S82. Scale bar is equal to 1 mm.

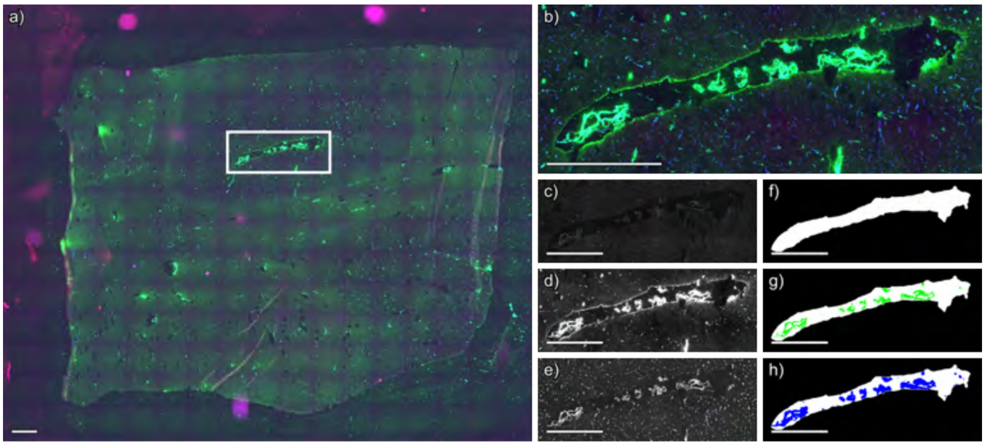


**Figure S82** Segmentation of immunofluorescence microscopy image of case 13, technical replicate 3, positive ion mode. (a) Whole slide immunofluorescence microscopy image. (b) Immunofluorescence image of region acquired by MALDI IMS, shown in a) as a white box. (c) Thiazine red stains amyloid, (d) anti-actin antibodies label smooth muscle, and (e) anti-collagen IV antibodies label vascular basement membranes. Segmentations for (f) amyloid, (g) smooth muscle cells, and (h) collagen IV. Per-channel standardization was applied across the entire dataset to ensure that color intensities are directly comparable across Figures S9-S82. Scale bar is equal to 1 mm.

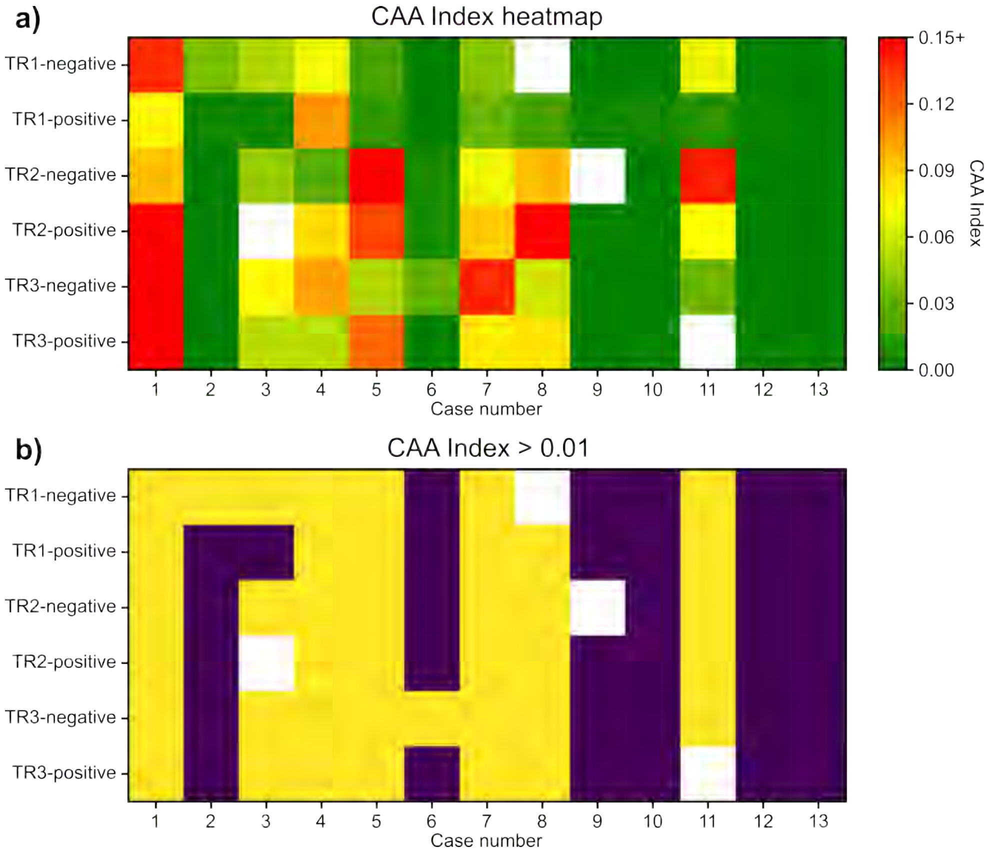


**Figure S83** Calculations of CAA index using acquired immunofluorescence microscopy segmentation masks (a). Samples marked yellow surpassed CAA index, while samples marked purple fell below CAA index (b).

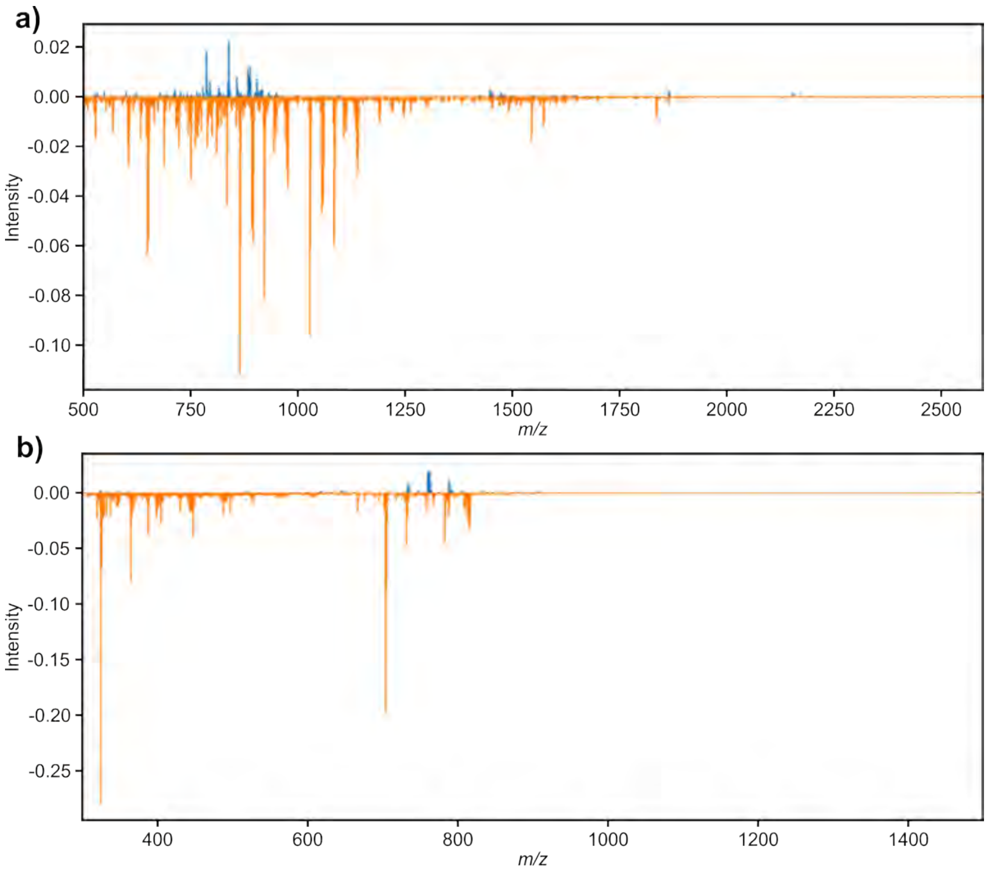


**Figure S84** Negative ion mode difference spectrum (a) and positive ion difference spectrum (b) between the average spectrum of the CAA-present group and CAA-absent group. Ions more expressed in CAA-absent are shown in blue, while ions more expressed CAA-present are shown in orange.

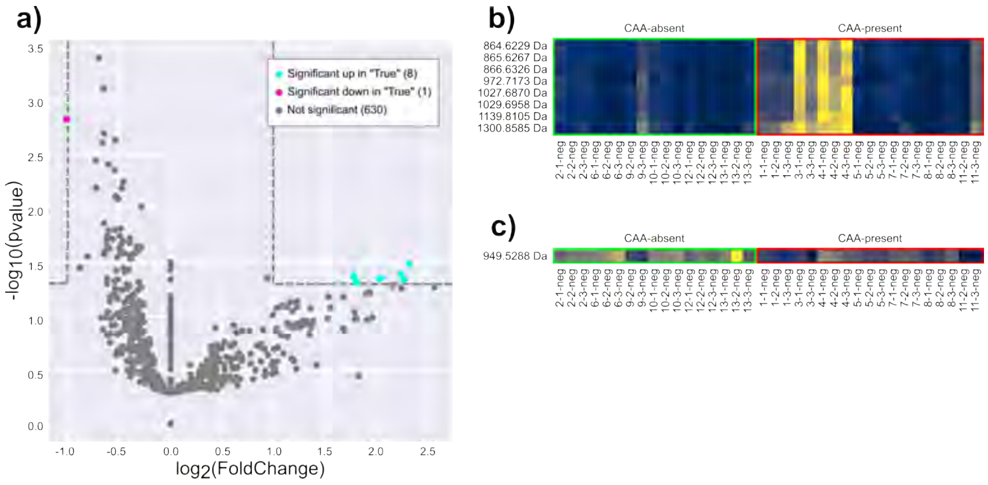


**Figure S85** Univariate comparisons of average mass spectrums in negative ion mode of each case between CAA-present and CAA-absent groups, where CAA-present is represented as True (a). Ions significantly higher in CAA-present (b) and higher in CAA-absent (c). Labels are in order of case number, technical replicate, and polarity. The tile color (blue-yellow) represents the average intensity of pixels inside the vasculature masks for a given ion.

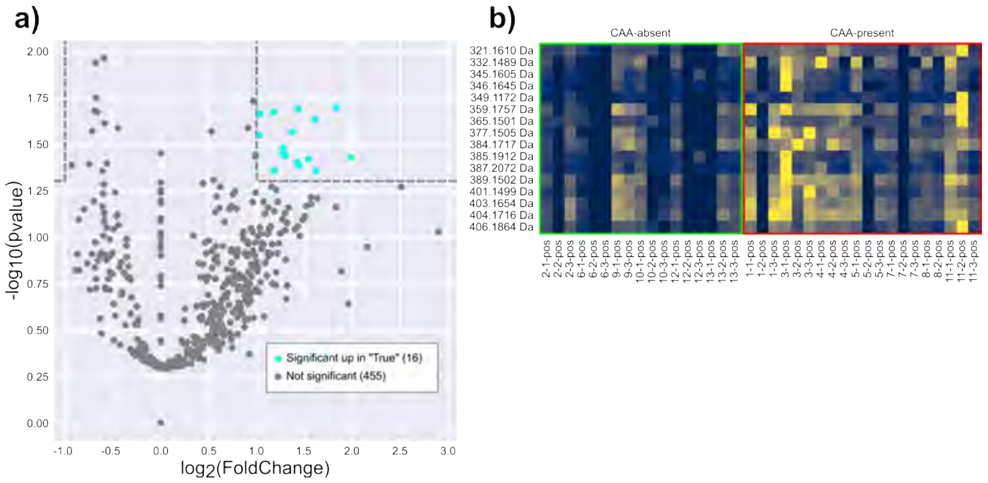


**Figure S86** Univariate comparisons of average mass spectrums in positive ion mode of each case between CAA-present and CAA-absent groups, where CAA-present is represented as True (a). Ions significantly higher in CAA-present (b) and higher in CAA-absent (c). Labels are in order of case number, technical replicate, and polarity. The tile color (blue-yellow) represents the average intensity of pixels inside the vasculature masks for a given ion.

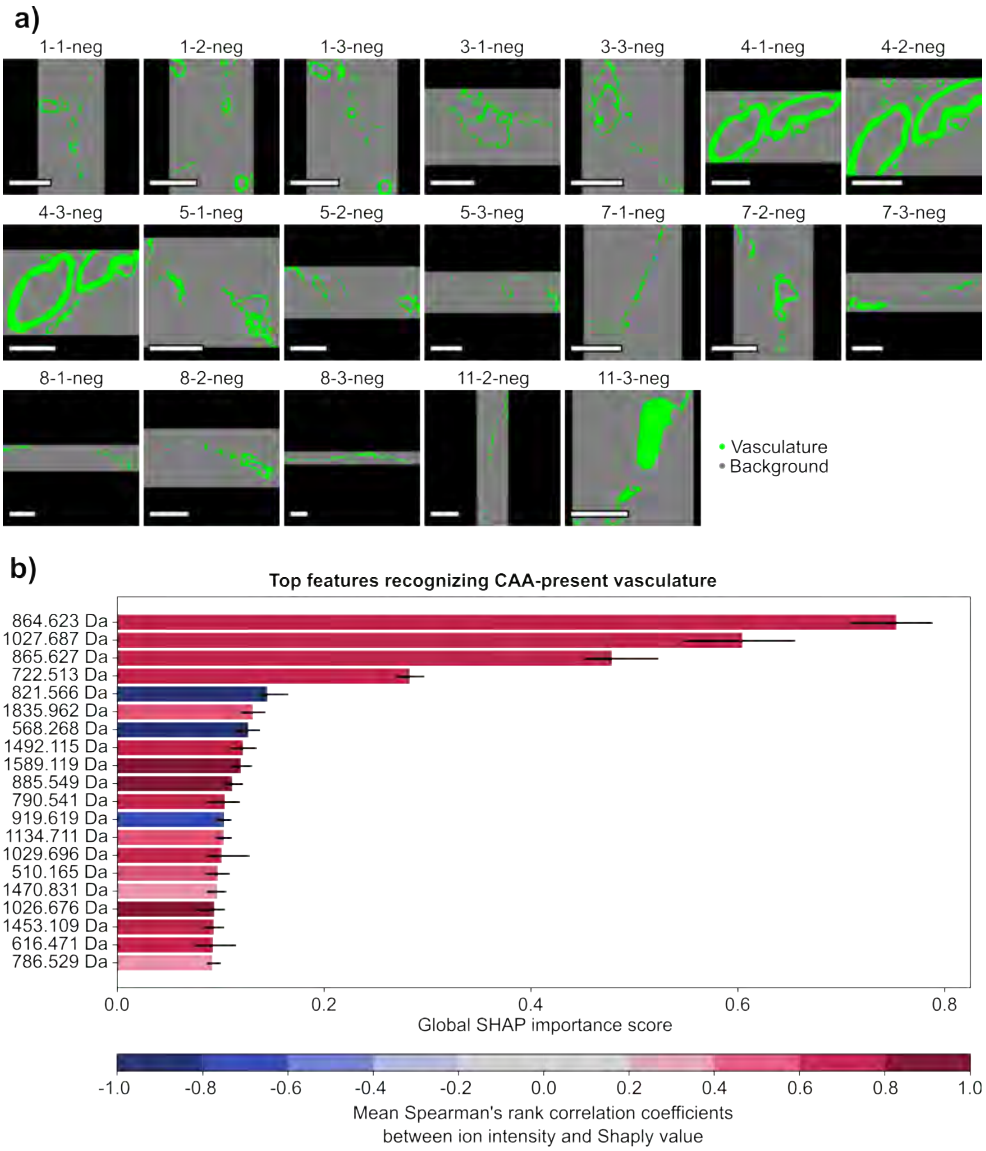


**Figure S87** Negative ion mode CAA-present vs background masks (a) and top features recognizing CAA-present masks (b). Scale bar is equal to 1 mm. Labels are in order of case number, technical replicate, and polarity.

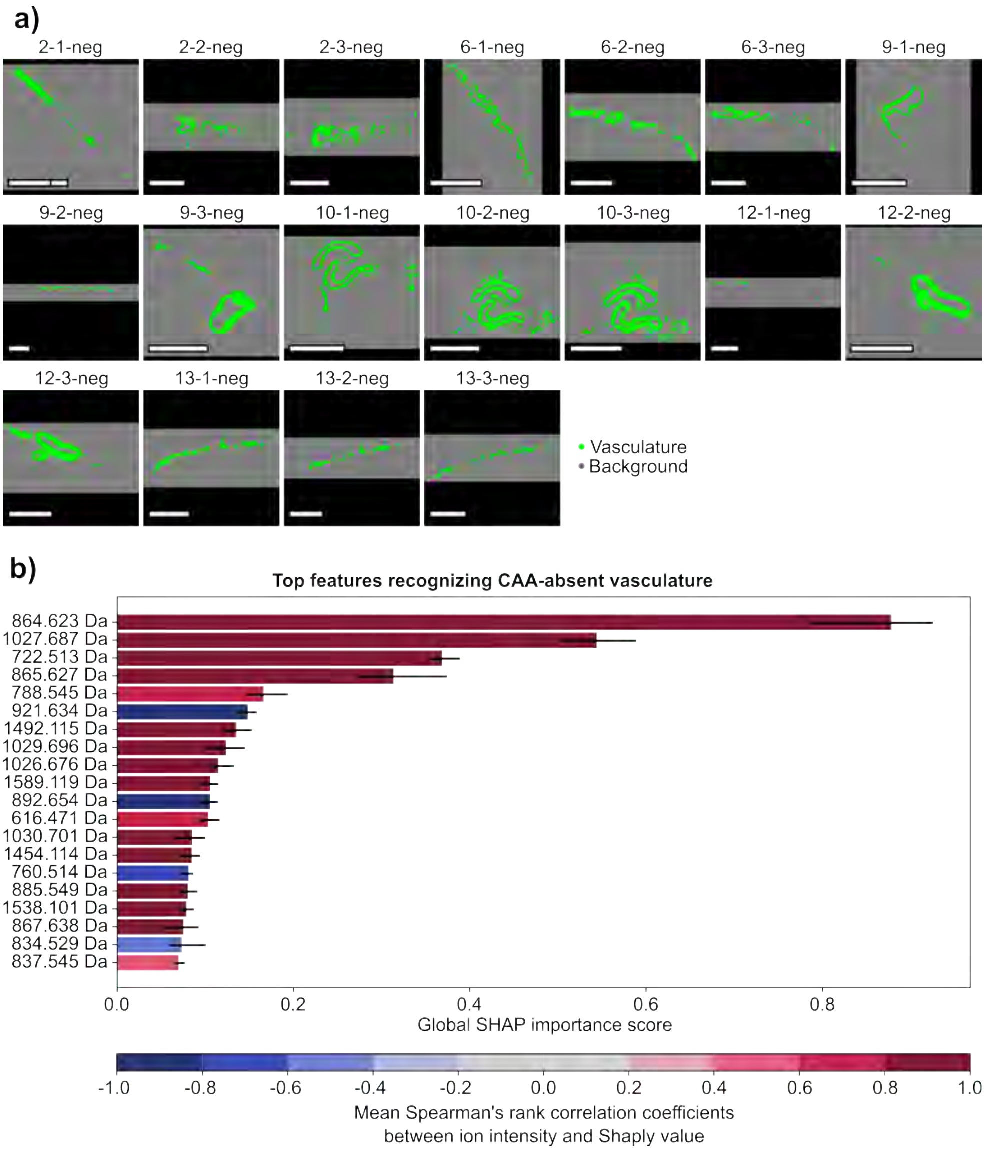


**Figure S88** Negative ion mode CAA-absent vs background masks (a) and top features recognizing CAA-absent masks (b). Scale bar is equal to 1 mm. Labels are in order of case number, technical replicate, and polarity.

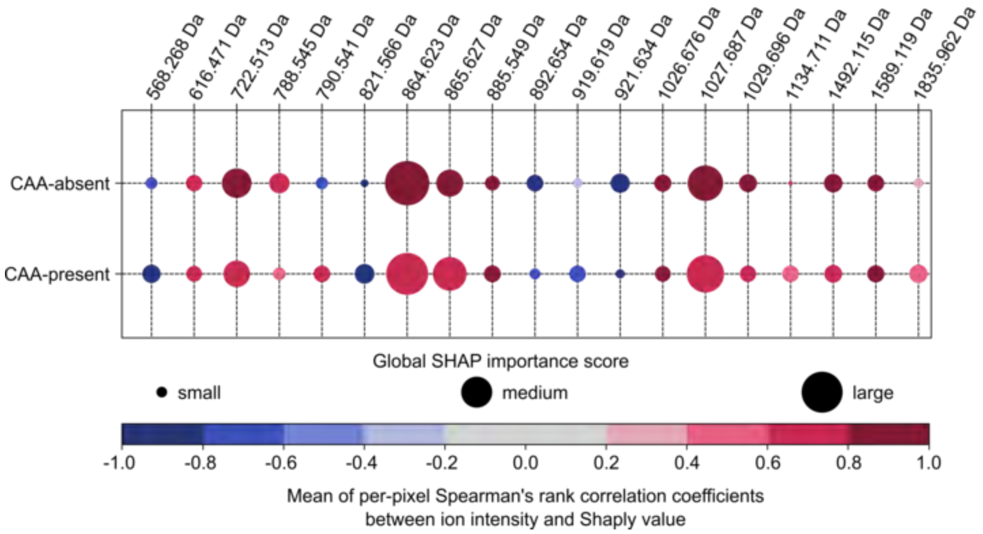


**Figure S89** Global SHAP importance score of negative ion mode CAA-present masks vs background masks compared to CAA-absent masks vs background masks. The size of each bubble indicates the global SHAP importance of a given feature (column) to recognizing CAA-absent and CAA-present samples (row), with the color indicating a positive (red) or negative (blue) correlation of the feature abundance to that class.

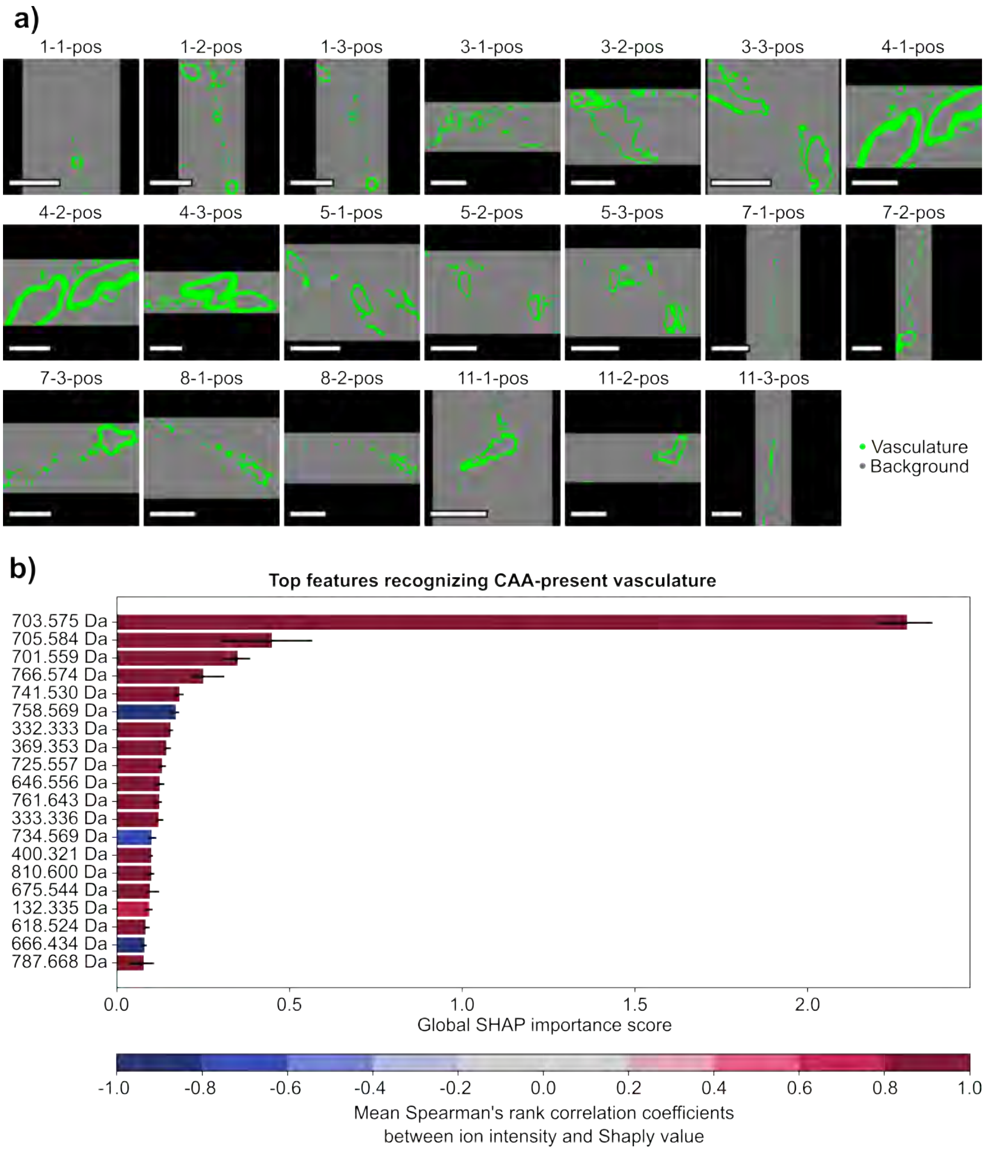


**Figure S90** Positive ion mode CAA-present vs background masks (a) and top features recognizing CAA-present masks (b). Scale bar is equal to 1 mm. Labels are in order of case number, technical replicate, and polarity.

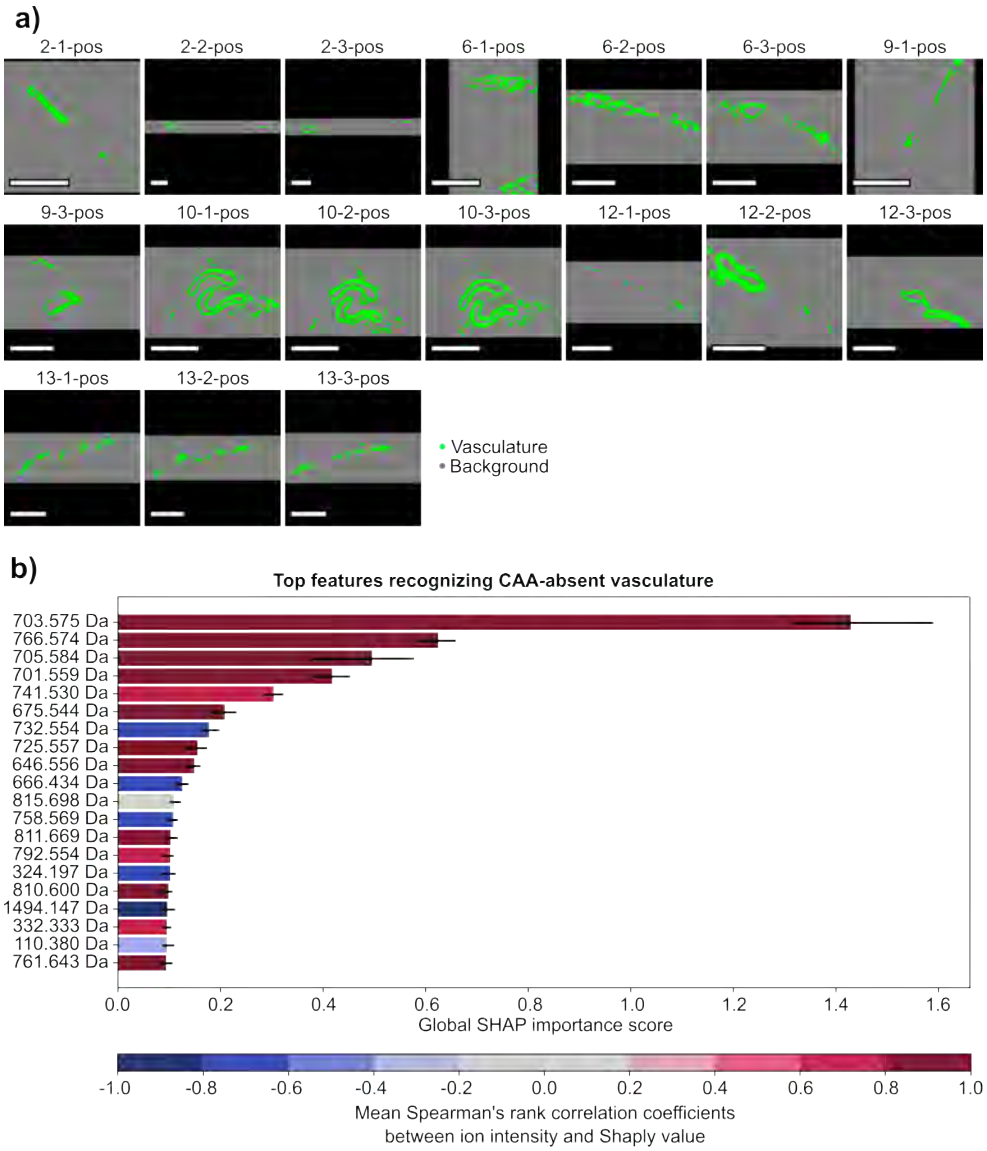


**Figure S91** Positive ion mode CAA-absent vs background masks (a) and top features recognizing CAA-absent masks (b). Scale bar is equal to 1 mm. Labels are in order of case number, technical replicate, and polarity.

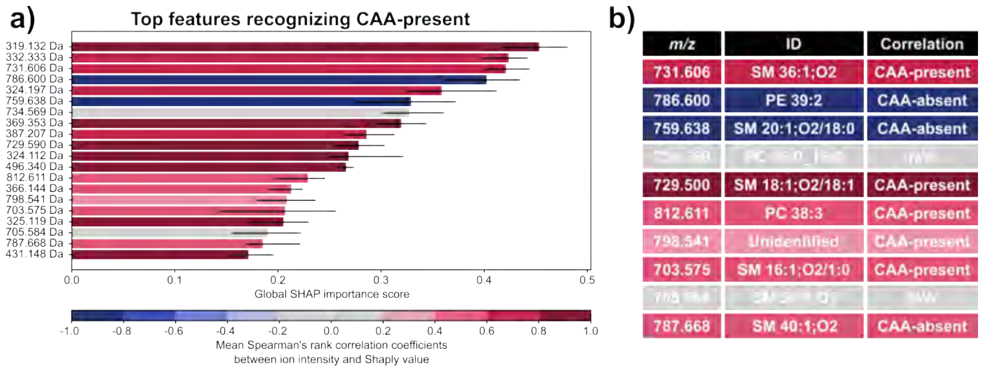


**Figure S92** Positive ion mode CAA-present vs CAA-absent. Top features recognizing CAA-present (a) and lipid annotations (b). Scale bar is equal to 1 mm. Labels are in order of case number, technical replicate, and polarity.
